# Sexually discordant selection is associated with trait specific morphological changes and a complex genomic response

**DOI:** 10.1101/2023.08.31.555745

**Authors:** Tyler Audet, Joelle Krol, Katie Pelletier, Andrew D. Stewart, Ian Dworkin

## Abstract

Sexes often have differing fitness optima, potentially generating intra-locus sexual conflict, as each sex bears a genetic ‘load’ of alleles beneficial to the other sex. One strategy to evaluate conflict in the genome is to artificially select populations discordantly, against established sexual dimorphism, reintroducing attenuated conflict. We investigate a long-term artificial selection experiment reversing sexual size dimorphism in *Drosophila melanogaster* during ∼350 generations of sexually discordant selection. We explore morphological and genomic changes to identify loci under selection between the sexes in discordantly and concordantly size selected treatments. Despite substantial changes to overall size, concordant selection maintained ancestral sexual dimorphism. However, discordant selection altered size dimorphism in a trait-specific manner. We observe multiple, possible soft selective sweeps in the genome, with size related genes showing signs of selection. Patterns of genomic differentiation between the sexes within lineages identified potential sites maintained by sexual conflict. One discordant selection lineage shows a pattern of elevated genomic differentiation on chromosome 3L, consistent with the maintenance of sexual conflict. Our results suggest measurable signs of conflict and differentially segregating alleles between the sexes due to discordant selection.

## Introduction

While in many species the sexes show phenotypic differences, they must also use nearly the same genome during development to express these phenotypes. This sexual dimorphism (SD) evolves despite the fact that the sexes have a high genetic correlation (*r_MF_*) for many traits, which can hinder the evolution of SD (Lande 1980). The extent to which SD can evolve depends on the strength and direction of selection, additive genetic variance, and the inter-sex genetic correlation (Lande 1980; Delph et al. 2011). Many organisms show high *r_MF_,* that could limit the potential for sexually discordant evolution (Lande 1980; Poissant et al. 2010). Despite this, SD is very common in nature, in particular, sexual size dimorphism (SSD; reviewed in Fairbairn et al. 2007). In many insects including the pomace fly *Drosophila melanogaster*, females are most often the larger sex (Ashburner 1989). This dimorphism is likely due to the relative contribution of increased fecundity with increased size in females (Honěk 1993; Reeve and Fairbairn 1999). Male-biased SSD occurs in some insects, often due to the relative contribution of sexual selection, and can evolve rapidly within a clade (Emlen et al. 2005; Moczek et al. 2006; Luecke and Kopp 2019). When loci impact the phenotype in a way favouring one sex, but disadvantageous to the other, it may create intra-locus sexual conflict (IASC). This IASC combined with a high *r_MF_* between the sexes, may impede the rate of evolution if genetic variation for dimorphism is not also high. Despite high *r_MF_*values, family-based artificial selection has demonstrated that sex specific responses can occur relatively rapidly (Alicchio and Palenzona 1971; Bird and Schaffer 1972; Eisen and Hanrahan 1972). Not only can changes in SSD occur rapidly with family-based artificial selection, but *r_MF_* has been directly selected upon and degraded in just a few generations (Delph et al. 2011). This reduction of *r_MF_* allowed one sex to be selected on for size with minimal response in the unselected sex, where sex specific selection in high *r_MF_* lines resulted in a strong correlated response in the unselected sex. Although kin selection experiments demonstrates additive genetic variation for a response to sexually discordant selection, these do not reflect patterns of transmission of allelic effects in most natural populations. As such, approaches based on mass artificial selection may better reflect transmission of allelic effects in natural populations. Using strong, long-term, mass artificial selection, Stewart and Rice (2018) demonstrated that response to a sex-discordant selection pressure can occur. Stewart and Rice (2018) successfully selected in a sex-discordant manner on body size in *D*. *melanogaster* over 250 generations, with measurable phenotypic changes requiring more than 100-150 generations of artificial selection. In comparison, sex-concordant selection for body size resulted in rapid phenotypic responses. Despite discordant selection responding in family and mass selection experimental designs, one previous study using *Tribolium castaneum*, found little response to sex-discordant selection (Tigreros and Lewis 2011), despite a rapid response to sexually concordant selection on pupal mass. This suggests at least some barrier to divergent response in sexual dimorphism may exist in populations, although it may reflect the modest genetic variation in the ancestral populations and the number (7) generations of artificial selected applied.

The sex-concordant (hereafter, concordant) selection lineages established by Stewart and Rice have had their genomes sequenced at generation 100 to identify candidate SNPs associated with body size variation (Turner et al. 2011). The sex-discordant (hereafter, discordant) lineages have not previously been sequenced, and are the main focus of this current study. To date, functional genetic analyses in *Drosophila* have implicated a few pathways involved with sex determination and growth, that can contribute to SSD. Manipulation of sex-specific splice variants of *transformer* (*tra)* reduces female size, reducing (but not eliminating) SSD (Rideout et al. 2015). Increasing quantities of *tra* also showed a sex-specific increase of size in females (Rideout et al. 2015). A duplication of the *diminutive* (*myc*) gene on the X chromosome resulted in males 12-14% larger, and when paired with constitutive expression of *tra*, SSD was substantially diminished (Mathews et al. 2017). Upstream of *tra*, tissue specific depletion of *sex-lethal* (*sxl*) in neurons, led to reduction of female body size (Sawala and Gould 2017). Inhibition of insulin-signalling (IIS), had sex-specific impacts on body size, largely reducing female body size. In contrast, up-regulation of IIS increased male body size (Millington et al. 2021b). These experiments demonstrate pathways involved with the phenotypic expression of sexual dimorphism, but it is unclear whether segregating variation in these pathways contribute to natural phenotypic variation. Although alleles of large effect in any of the above genes would be exciting to find in a natural population, it is not a safe assumption that genes showing a phenotypic response in a lab setting will be the genes selected upon if a population undergoes discordant selection. Further, rapid response to selection on size as demonstrated by Stewart and Rice (2018) and Bird and Schaffer (1972) among others, suggests that a polygenic response using standing genetic variation, rather than *de novo* mutations, in growth signalling pathways is the likely mechanism for short term body size evolution. In natural populations, segregating alleles contributing to variation in sexual dimorphism interest could be being maintained in the population by selection, or simply reflect mutation-selection-drift balance.

Populations of *D. melanogaster* harbour alleles with both sexually discordant and concordant effects on fitness (Rice and Chippindale 2001). Variation for body size in *D. melanogaster* is highly polygenic (Carreira et al. 2009; Turner et al. 2011). Sex limited selection on males has been shown to incur a fitness cost to females (Prasad et al. 2007). This male limited evolution also results in a change in body size in non-selected females that is closer to the male optimum (Prasad et al. 2007). A decrease in female fitness (and body size) when males are allowed to evolve toward their own optimum without parallel female evolution suggests that there is unresolved sexual conflict in the genome of *D. melanogaster* pertaining to body size. Alleles potentially under conflict in the genomes of the outbred LH_m_ population of *D. melanogaster* have been identified, via examination of male mating success and female fecundity (Ruzicka et al. 2019). Alleles under IASC influencing body size, can help understand how the genome responds to selection with divergent phenotypic optima. Selection on body size in a discordant manner has the potential to answer questions of how the shared genome overcomes high *r_MF_*when selection favors divergent phenotypes across the sexes.

Using lineages evolved under sexually discordant selection for size, first described in Stewart and Rice (2018), we demonstrate two important findings. First, despite selection for a trait-agnostic (*i.e.* selection on general size rather than an increase in mass, or thorax length) measure of size, trait specific patterns of SSD reversal are the norm. Second, despite the long term sexually discordant selection, we see relatively weak evidence for sexually antagonistic alleles being maintained. We do, however, identify one region potentially segregating differentially between the sexes, a possible sign of unresolved (or re-introduced) sexual conflict. Using an evolve and sequence approach we not only examined among lineage patterns of genomic differentiation, but within lineage among the sexes. We discuss our findings within the context of both the evolution of sexual dimorphism and the potential role of ongoing intra-locus sexual conflict.

## Materials & Methods

### Lineages

The flies used are part of long-term experimental populations for size selection (Turner et al. 2011; Stewart and Rice 2018). The selection lineages examined were started using the outbred population, LH_M_, previously adapted to the lab for over 350 generations prior to selection (and diverged from the LH_M_ population used in Ruzicka et al. (2019) for an unknown number of generations). These flies are maintained in discrete two-week generations at moderate density (approximate 200 eggs per 10ml standard molasses food vial (Table S1). The complete methodology for maintenance of the base population has been published previously (Rice et al. 2005). For the size selection, flies are anesthetized using CO_2_ and sorted using a motorized stacked sieving device in which each successive sieve is 5% smaller than the sieve above. The largest sieve used had aperture diameters of 2000μm and the smallest sieve had apertures measuring 850μm. In each generation, all flies (∼1800 individuals), from a selection lineage were sorted by sieve, and 10 vials of 16 mating pairs (320 total flies per lineage) were selected based on phenotype, to generate the next generation. One treatment used only the smallest flies of both sexes (S; concordant selection), one treatment the largest of each sex (L; concordant selection), and the reversal of dimorphism (E; discordant selection) treatment used smallest females and largest males. Finally, a control treatment was populated with flies passed through sieves, but not selected based on size, as controls (C). Two replicate lineages were maintained for each selection treatment. During subsequent selection for size with the established protocol, it was observed that selection can be more stringent on small flies than large because of the nature of the sieve sorting (hinderances due to appendages sticking out, blocking passage of flies through sieve). This would imply that in the sex-discordant selection females being selected to be small were under increased selective pressure compared to males selected to be large. This would suggest that there was a greater pressure for selection in one direction and although this does not explain all results, it may influence response to sexually discordant selection.

### Morphological measures and analysis

At generation 367 (August 19^th^, 2019) flies from each lineage were collected and stored in 70% ethanol for dissection and measurement of traits. Individuals were chosen randomly before and after the selection treatment to get an accurate estimate of size in the overall lineage. Individuals were dissected under a Leica M125 microscope and legs and thoraces were imaged at 63x using a DCF-400 camera. A minimum of 20 flies (for each sex) were dissected from each lineage by dissecting off the first (pro-thoracic) right leg, then imaging the left side of the thorax of each individual. The leg and left side of the thorax were mounted on slides (in 70% glycerol in PBS, with a small amount of phenol as a bacteriostatic agent). The femur, tibia, and first tarsal segment of each leg, and each thorax, was measured using ImageJ version 1.52q (Schneider et al. 2012). We measured thorax length as a proxy for overall size and a proxy for the general pattern of the female-biased SSD found in *Drosophila melanogaster*. We also measured each leg segment; measurement was completed from the center of the beginning of the segment to the center of the end of the segment.

Analysis of the leg and thorax measurements were completed using R v4.1.3 (R Core Team 2022). One individual measure of femur was removed as it’s measure was ∼100X smaller than the mean for the trait. For all analyses of morphology, we used log_2_ transformed trait values to facilitate inferences of proportional changes in dimorphism. We fit general linear mixed models for individual traits including sex, selection and sampling and their 2^nd^ order interactions as fixed effects. We allowed sex effects to vary as a random effect of replicate lineage (i.e. random slopes). Models were fit using lmer in the lme4 package v1.1.30 (Bates et al. 2015). We confirmed the results with a fully multivariate mixed model for all traits. For the multivariate mixed model, we allowed for random effects of sex and traits by lineage, as well as accounting for within-individual variation across traits. The initial fit of this model using lmer was singular, likely due to variance estimates getting “stuck” on a boundary (0). We employed two approaches to deal with this. First, we employed a Bayesian extension of our model using blmer in blme v1.0.5 (Chung et al. 2013). This employs weak regularizing priors (away from 0 for the variances). We confirmed the stability of fixed effects using a second approach, fitting a general latent-variable mixed model, to estimate reduced-rank covariance matrices for random effects, as implemented in glmmTMB v1.1.7 (Brooks et al. 2017; Niku et al. 2019; Kristensen and McGillycuddy 2023). Estimated marginal means, custom contrasts, and associated confidence intervals were estimated using emmeans v1.8.0 (Lenth et al. 2018). Visualization was done using ggplot2 v3.3.6 (Wickham 2018). These approaches provided very similar fixed effects estimates to each other, and to the single trait models, which were used for downstream analysis.

Despite artificial selection being “trait-agnostic” (selecting on a composite cross-sectional area, with possible hinderance from appendages), and a relatively consistent response as outlined below in the results, we examined changes in multivariate allometry among traits within each selective treatment. Commonly, the first principal component derived from a variance-covariance matrix of log transformed morphological measures captures a measure of overall size (Jolicoeur 1963; Blackith and Reyment 1971; Klingenberg 1996). It is common to assess whether the log transformed traits contribute approximately equally to size for PC1 (isometry), with expected loadings of 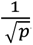, where *p* is the number of traits (dimensions). In addition to examining (and visualizing) these vectors from principal components analyses conducted by each selective treatment, we compared aspects of the orientation and structure of the variance-covariance matrices across treatments.

### Sex ratio cross

While phenotyping flies for an unrelated experiment, a deviation of the expected 1:1 sex ratio was observed in only a single cross, between our control lineage 1 and discordant lineage 1 samples. As summarized in the results, discordant lineage 1 is also the lineage with potential evidence of maintained conflict on chromosome 3L. As such we conducted experimental crosses to examine adult sex ratios to evaluate whether there is a consistent deviation in sex ratio, potentially due to genomic conflict. At generation 464, 15 males and 15 females were taken from each treatment for single pair matings to an opposite sex individual from the originating LH_m_ population. The single pair reciprocal matings were allowed to lay until larvae was visible in the food (∼5 days), before the F_0_ pair were placed in 70% ethanol. From each F1 vial, a single sibling pair was used to generate an F2, while the rest of the F1 individuals were stored in 70% ethanol after being allowed to eclose until most pupal casings were visibly empty. Adults from the F2 generation, once most eclosed, were stored in 70% ethanol. Number of male and female offspring in F1 and F2 generations were counted.

We modeled the data for the sex ratio crosses, using a logistic regression (glm in R), with counts of males and females from each cross as the unit of sampling, with lineage and cross direction and their interactions as predictors. From the model fits, we computed estimates and their confidence intervals on the response scale using emmeans, to facilitate interpretability. We did this both with and without the reciprocal direction of genetic crosses (whether or treatment individual was the sire or dam) in the model. During the experiment to examine adult sex ratios, we noticed substantial differences in number of individual offspring. While this was not a planned analysis (and should be treated as such), we examined differences in fecundity (assessed by census of adults) fitting a general linear model (fit using lm) of number of offspring regressed onto direction of cross, selection treatments and their interactions.

### Genomic sample preparation

At generation 378 (February 17^th^, 2020) flies from each lineage were collected in 70% ethanol and stored at -20°C (Figure 1). For each lineage and sex, flies were separated in to four pools of 25 and DNA was extracted using a column-based DNA extraction kit (DNeasy Qiagen kit, Cat # 69506). The extracted DNA from each of the four pools of 25 for distinct samples were combined such that the same concentration of DNA was added from each pool (equimolar). This resulted in pools of 100 individuals for each combination of sex and unique lineage that were sequenced. This 25 individual pooling was done due to the size of the columns not being capable of extracting from 100 individuals at once. Library preparation and sequencing was done by Génome Québec (Centre d’expertise et de services, Génome Québec). Library preps were prepared with IDT dual index adapters. Sequencing was done using Illumina NovaSeq 6000 to an average coverage of 200x with 151bp sequence fragments. Initial sequencing fell short of 200x coverage, so additional sequencing from the same libraries were conducted to ‘top-up’ coverage on some samples and were merged with their corresponding samples after mapping.

**Figure 1:**
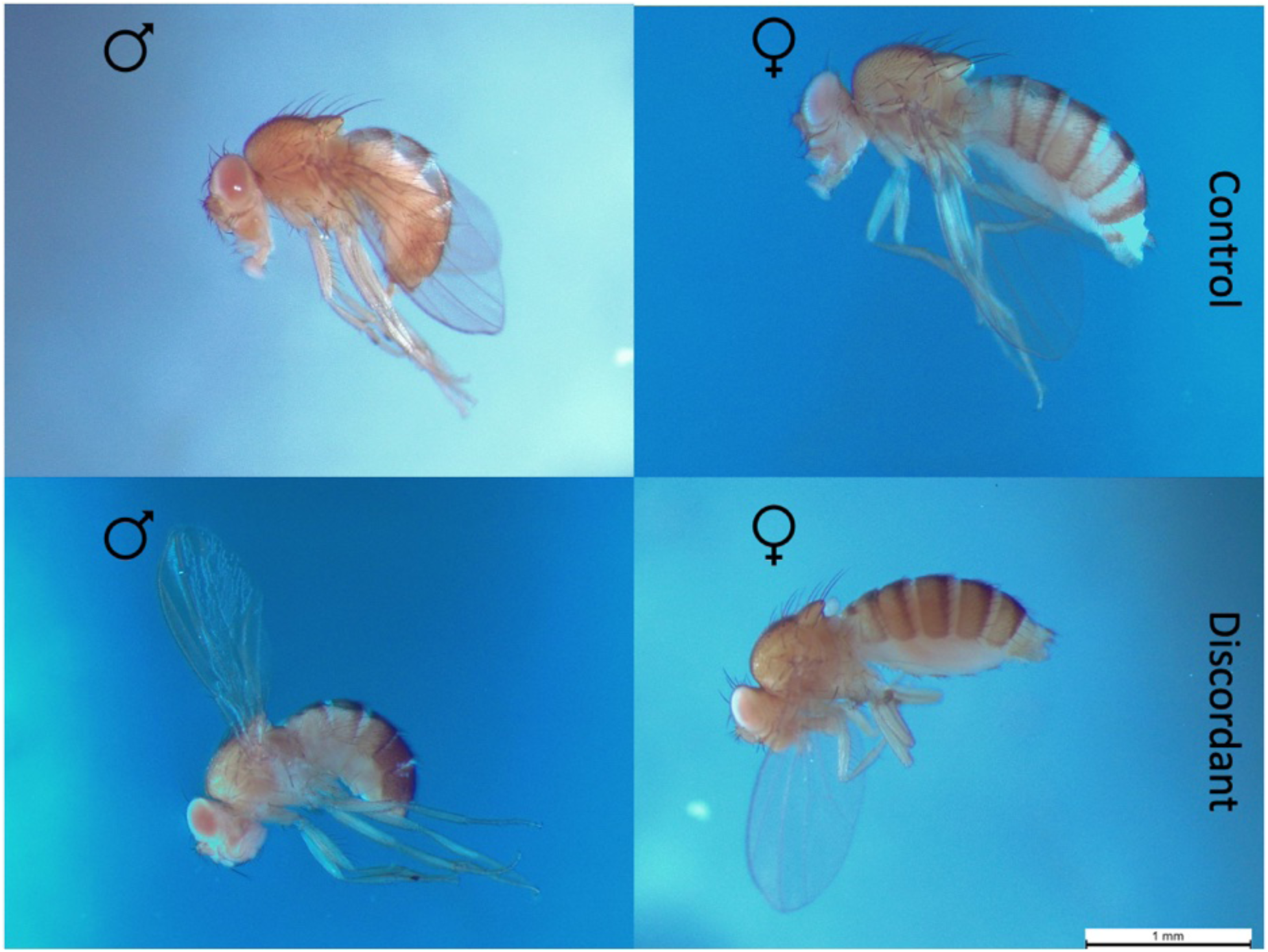
Stereomicroscope images of Control and discordant males and females (25x magnification). Flies chosen at random from populations stored in 70% ethanol and imaged in 70% ethanol.

### Bioinformatic Pipeline

Figure S1 shows a summary of the bioinformatics pipeline. A detailed summary can be found at (https://github.com/TylerAudet/SSD_workflow). Reads were trimmed using bbduk v38.90 (Bushnell 2021), aligned using BWA-mem version 0.7.8 (Li 2013), GATK v3.8 was used to mark indels, and perform local realignment. SAMtools v1.5 (Li et al. 2009) was used to convert SAM files to BAM, extract out the core genome, mark/remove read groups, and create mpileups. Three distinct mpileups were made: one with sexes and lineages separate, one with sexes merged maintaining lineages, and one where treatments were pooled together by merging both sexes and lineages together, each of these three went through the same SNP calling and file generation methods. Popoolation v1.2.2 (Kofler et al. 2011) was used to mask repetitive regions identified with Repeat Masker v4 (Smit and Hubley 2013), as well as indels for SNP calling. PoolSNP v1 (Kapun et al. 2020) was used to call variants. From this VCF file indels were masked out using custom scripts from the DrosEU pipeline (Kapun et al. 2020). Due to issues generating sync files from poolSNP VCF output files, a sync file was also generated from the mpileup using Grenedalf version 0.2.0 (Czech et al. 2023). This sync file was filtered for sites present in our SNP called VCF file to create a SNP-called sync file. For ‘sexes separate’ as well as ‘treatments pooled’ sync files, through testing, we did observe that changes in the bioinformatic pipeline such as which program was used changed the results slightly, pointing at artifacts introduced by bioinformatic programs. This suggests that great hesitancy and meticulousness must be applied when selecting programs for bioinformatic analyses, as these choices certainly change final results in, at best, a small way. Bioinformatics were done on Compute Canada servers on the Graham cluster.

### Among population Genomic Analyses

To identify variants that potentially contribute to phenotypic divergence between treatments, we examined three measurements of population differentiation between populations or genetic diversity within populations. To assess variation between populations we used two related approaches, based upon both F_ST_, and a modified Cochran–Mantel– Haenszel (CMH) statistic. To assess variation within populations to identify potential regions that have undergone a selective sweep, we performed windowed computation of nucleotide diversity (π). We calculated *F_ST_*between control treatments and discordant treatments using the Nei estimation in Grenedalf v0.2.0 in 10kb windows along each chromosome (Czech et al. 2023), which corrects for sample size and pooled sequencing size, known sources of error in parameter estimation for pooled sequencing. Although our primary goal was on the changes to SSD, not overall size, we also examined *F_ST_* between Large and Small treatments to follow up on the initial analysis in Turner et al. (2011) and see if their findings were replicated after an additional ∼275 generations of artificial selection on body size. To do so we ran the sequence data from generation 100 through our pipeline and compared genes of interest from F100 to the genes identified at F378. Effective population size (*N_e_*) was estimated as N_e_=181 for all samples using poolSeq v0.3.5 (Taus et al. 2017) using the formulas proposed in (Jónás et al. 2016) that are intended to work with pooled data as well as control for additional sources of sampling variance in allele frequencies that could bias N_e_. Using this N_e_, a CMH statistic was calculated using ACER version 1.0 (Spitzer et al. 2020) for each treatment and replicate as well. Since an ancestral sample was not preserved, drift between control and discordant treatments (or large versus small) were used, and the generations of drift was set at 700, from the control treatment (∼generations to convergence x2).

For our samples, given the large number of generations of selection, a hard selective sweep would likely result in regions of low nucleotide diversity, which we measured as π using Grenedalf v0.2.0 in 10kb windows for our control, discordant, and concordant samples. This resulted in us having: per sample π values for 10kb windows, F_ST_ values for 10kb windows between discordant and control and between large and small, and SNP-by-SNP adjusted p-values from a CMH test. Using these values, regions with a small π value (< 0.0005) were extracted as interesting, and were extracted along with the top 5% of *F_ST_*(discordant versus control cut-off was *F_ST_* 0.51 with 621 windows meeting this criteria; large vs. small cut-off was *F_ST_* 0.78 with 621 windows meeting that criteria) and SNPs with a CMH adjusted p-value < 0.01 (using *p.adjust* in R; Benjamini and Hochberg 1995; 132290 SNPs in control vs. discordant, 115747 SNPs in large vs. small) were extracted. The overlap between all windows or SNPs of interest was intersected with Bedtools v2.31.0 (Quinlan and Hall 2010) to get SNPs of interest Unfortunately, most software (including Grenedalf) does not account for hierarchical structure of populations (common in natural populations), nor the expected sources of drift and other forms of sampling variance among replicate lineages (within treatments) in evolve and resequence experiments. As such, the approach we used above (merging replicate lineages), does not account for sampling variation. To account for this, for the subset of “candidate” SNPs we identified we fit SNP specific logistic regression models, with SNP counts for each replicate lineage representing units of sampling, and with selective treatments as the predictor.

Treatment contrast between either discordant versus control, or large versus small treatments was obtained using emmeans. As this approach is relatively slow, we only did this for the sites we identified as likely candidates (discordant = 20014, large = 10649, small = 13952). Four files were created with final SNPs of interest: 1) “discordant replicate 1 region” All sites in and elevated region of *F_ST_* in replicate 1 of our discordant treatment on chromosome 3L (explained below; supplemental file 1), 2) “discordant sites” all interesting SNPs in the discordant treatment (supplemental file 2) 3) “Large sites” (supplemental file 3) 4) “Small sites” (same F_ST_ windows used as “large sites”; supplemental file 4). These sites were extracted from the SNP called VCF file using bedtools v2.30.0 (Quinlan and Hall 2010). Using this VCF file, a sync file was created, from which an allele frequency table for all sites of interest was made using Grenedalf, with the reference column set as the females from control replicate 1 major alleles.

Sites of interest found were annotated (Supplemental files 1,2,3,4). Gene ontology (GO) enrichment analyses were conducted using TopGO version 1.0 on each of these four subsets (Alexa et al. 2006). We also explored enriched GO terms using Gowinda v1.12 (Kofler and Schlötterer 2012), which controls for gene length and uses permutation to reduce false positives. Using 1000000 permutations in Gowinda we returned zero significant GO terms in our discordant and small SNPs of interest, and a single term in our large SNPs (GO:0030953: astral microtubule organization). Gene annotations were conducted using SNPeff version 5.1 (Cingolani et al. 2012).

### Within population Genomic Analyses to identify regions of genomic conflict

To identify genomic regions that may harbour variants contributing to IASC, we examined patterns of genomic differentiation between males and females from the same population and generation, for all of the evolved lineages (Cheng and Kirkpatrick 2016; Lucotte et al. 2016; Kasimatis et al. 2019). While simulation studies have suggested this approach generally has low power, the experimental design used for artificial, sexually discordant selection is well suited for this particular approach. We computed allele frequency tables for each sex within each lineage, along with FST between males and females within lineages in Grenedalf. As summarized in the results, we observed a region on chromosome 3L showing elevated *F_ST_*, in a single discordant treatment lineage (E1). To assess whether this elevated region of *F_ST_* could be accounted for by sampling variation, we simulated 1000000 sites, sampling the full range of allele frequencies used in our analyses, and simulating allele frequencies for male and female samples drawn from a common allele frequency in each simulation. We then allowed sequencing depth (based on the approximate empirical distribution) to vary for each sex. We plotted simulated allele frequencies for males and females, and over-plotted observed allele frequencies for each lineage, to determine the male-female differences in allele frequencies in this chromosomal region were extreme relative to distributions under our simulations that modeled sources of sampling variance. Additionally, we conducted a logistic regression (glm) of major and minor allele counts between the sexes. From this we obtained contrasts and confidence intervals using emmeans.

## Results

### Selection for size resulted in a trait dependent dimorphism when selection was sex-discordant

Consistent with the previous results for overall “body size” (Stewart and Rice 2018), all measured traits responded to artificial selection in the expected directions. Thorax length was measured as a proxy for body size and had a significant relationship between size and selection lineage (χ^2^ = 242.72 p <0.0001; Figure 2; Table S2; Figure S2; Figure S3) with some lineage specific variation (Figure S3). Concordant artificial selection for decreased size reduced thorax length relative to control lineages by ∼25% in females (26% in males), while selection for increased size increased thorax length by ∼9.5% for both sexes. In the discordant selection lineages, we found a ∼10% decrease in female thorax length, and minimal (∼0%) change in males. We observed similar patterns for length of leg segments. Small treatment males and females decreased in length relative to controls, to a similar degree (∼30% for femur, ∼24% for tibia and tarsus; Figure 3). Large treatment males and females increased in size somewhat more modestly relative to controls (∼2% for femur, 6% for tibia, 8% for tarsus). For sexually discordant selection, females decreased in size relative to controls (∼14% for femur, ∼7% tibia and tarsus). Males from the discordant lineage decreased more modestly relative to controls (∼8% for femur, < 1% for tibia, ∼3.5% tarsus). While some trait changes have confidence intervals that overlap zero, all traits and treatments seemed to respond in size in the expected direction of effect (Figure 2;3; Table S3; S4; S5).

**Figure 2:**
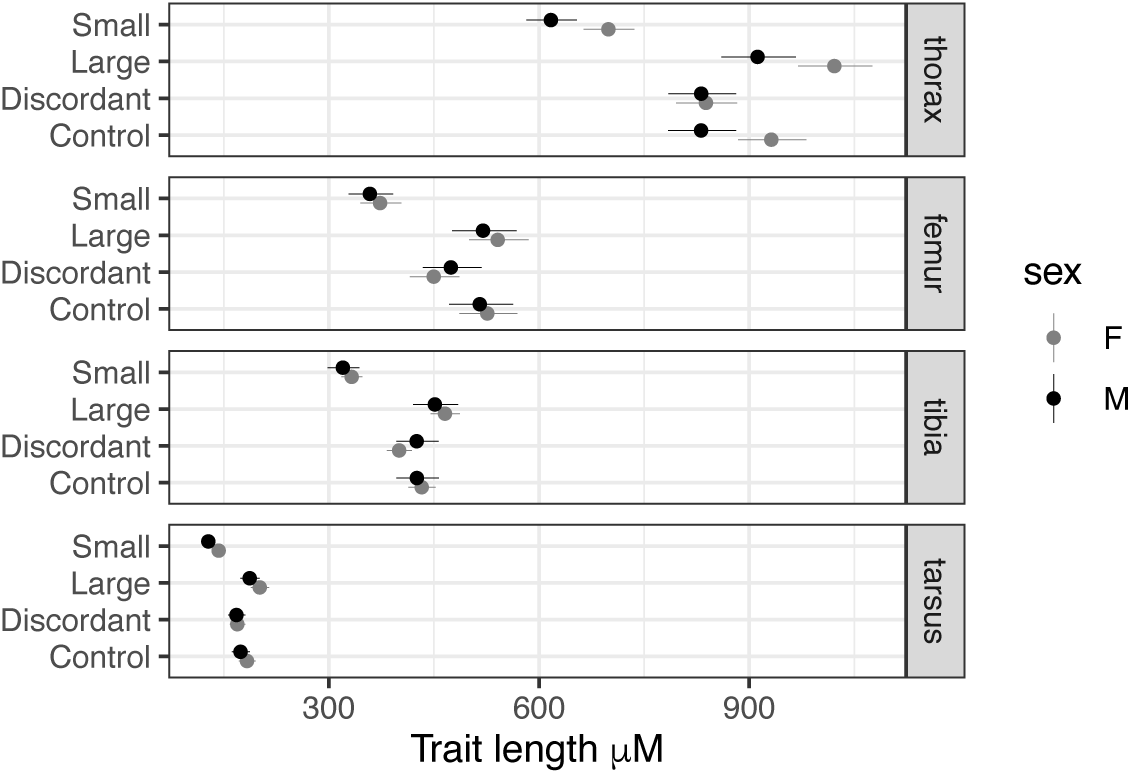
Model estimates for each trait, across treatments and sex. Measurements were log_2_ transformed for model fit, and back-transformed only for plotting. Error bars represent 95% confidence intervals.

**Figure 3:**
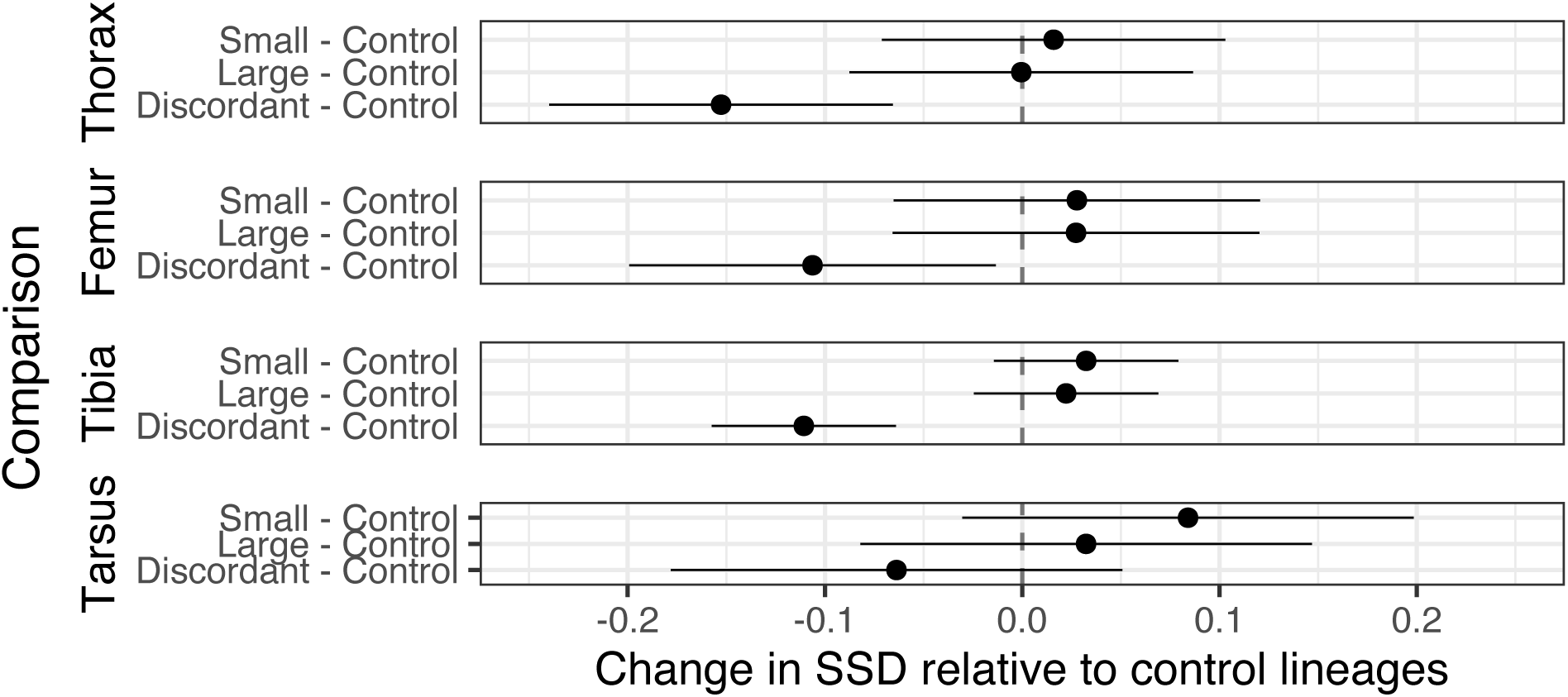
Substantial changes in sexual size dimorphism only occurs under discordant selection on size. Contrasts represent change in the amount of sexual size dimorphism between each artificially selected treatment, in comparison to controls, by trait. Modeled using log2 transformed length measures (μm), facilitating comparisons of proportional changes. Error bars are 95% confidence intervals for contrasts.

### Sexual dimorphism and multivariate allometry diverged only under discordant, sex-specific selection

Despite substantial changes in overall size, SSD remained female biased in all concordant selection treatment groups (Figure 2; 3), showing little change in dimorphism relative to control (Figure 3). In contrast, the sexually discordant selection treatment, resulted in a substantial reduction in the amount of ancestral female biased dimorphism, relative to the controls (Figure 3). Specifically, thorax and tarsus lengths have evolved to be essentially monomorphic, while femur and tibia lengths are now male-biased (∼5% increase) in size. The change in the amount of dimorphism was significant for all traits but tarsus length (Figure 3).

Given the results described above, it is not surprising we observed substantial changes in patterns of multivariate allometry (Figure 4; Figure S2, S3, S4). We observe a substantial change in the sex-discordant lineages, which deviates from the isometry vector (Table 1) that is observed for other *Drosophila melanogaster* populations (Shingleton et al. 2009). We further compared covariance matrices across selective treatments using the Krzanowski correlation. The discordant selection lineages show a reduced correlation to the control lineages (*r_Krz_* = 0.86) compared with the patterns observed for both concordant selection treatments (*r_Krz_* = 0.97 and *r_Krz_ = 0.99* for large and small respectively).

**Figure 4:**
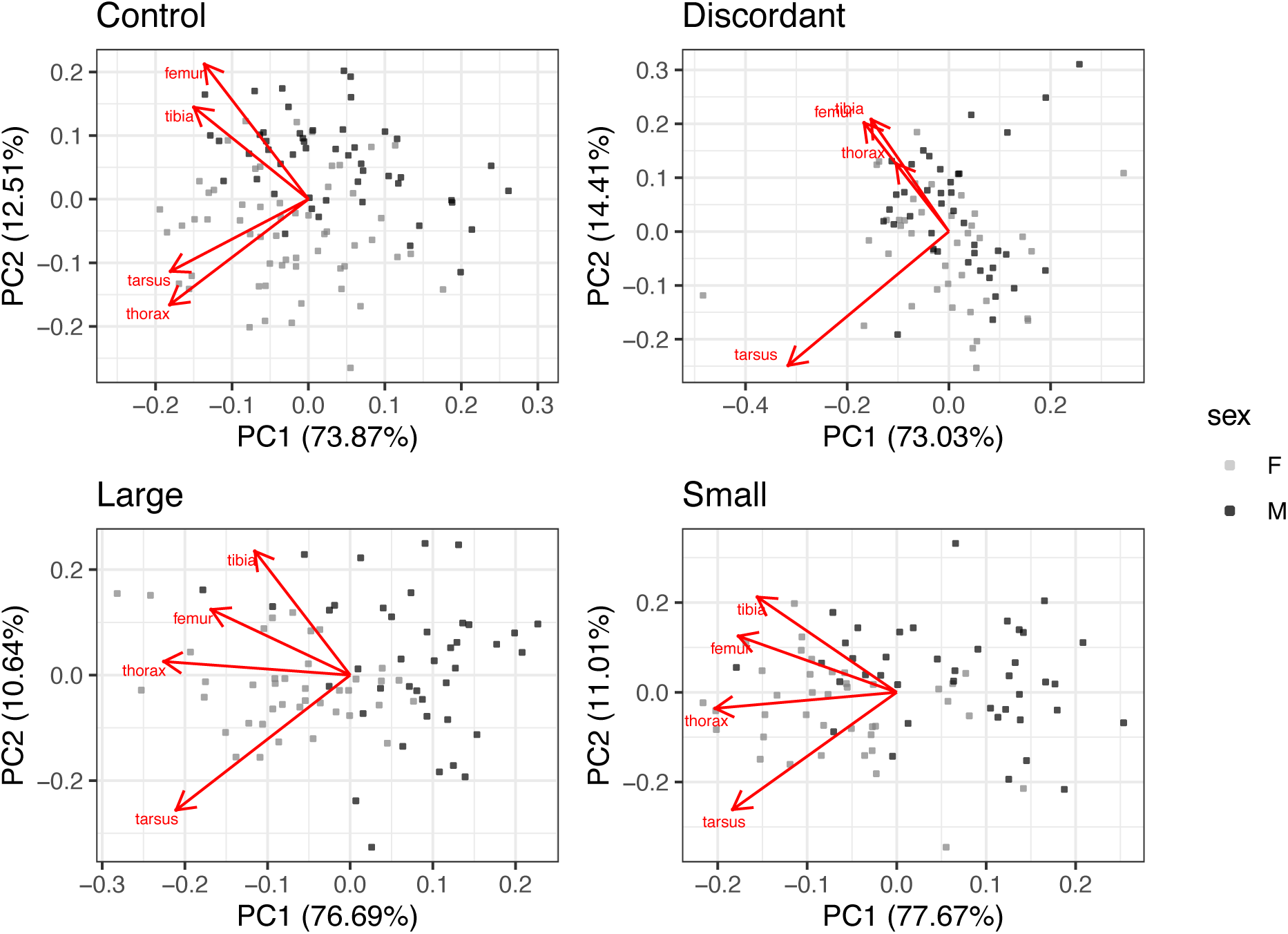
Sexually discordant selection alters patterns of multivariate allometry across sex. Represented as biplots, magnitudes and direction of the loadings for traits are superimposed onto PC1 and PC2. *Log_2_* transformed length measures were used.

**Table 1:**
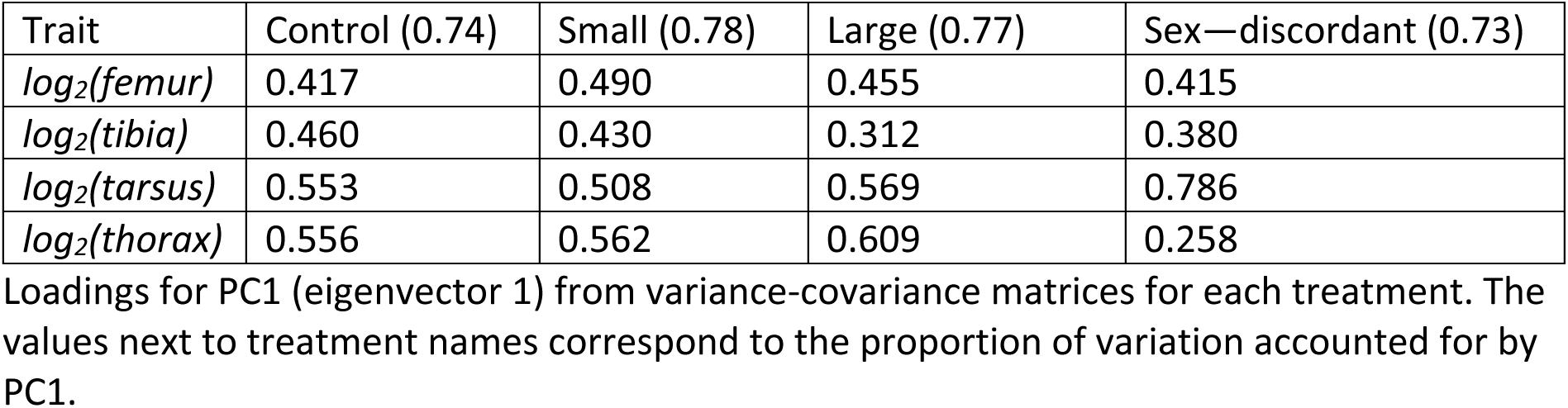
loadings for PC1, by treatment.

| Trait | Control (0.74) | Small (0.78) | Large (0.77) | Sex—discordant (0.73) |
| --- | --- | --- | --- | --- |
| $\log_2(\text{femur})$ | 0.417 | 0.490 | 0.455 | 0.415 |
| $\log_2(\text{tibia})$ | 0.460 | 0.430 | 0.312 | 0.380 |
| $\log_2(\text{tarsus})$ | 0.553 | 0.508 | 0.569 | 0.786 |
| $\log_2(\text{thorax})$ | 0.556 | 0.562 | 0.609 | 0.258 |

Loadings for PC1 (eigenvector 1) from variance-covariance matrices for each treatment. The values next to treatment names correspond to the proportion of variation accounted for by PC1.

### Genomic differentiation and nucleotide diversity (π) between selection treatments

*F_ST_* between control and discordant selection treatments was highly variable, but included substantial differentiation across the genome, and many regions at, or near, fixation (Figure 5A). Notably, average *F_ST_* is quite high across the chromosomes, consistent with a substantial impact of genetic drift on allele frequencies. When *F_ST_* is calculated between discordant and either of the concordant (small or large) treatments, we similarly see elevated *F_ST_* and multiple regions at, or near fixation (Figure S5; S6). In our discordant selection versus controls, the highest mean *F_ST_* is found on chromosome 2L (mean *F_ST_* per chromosome, 2L = 0.215, 2R = 0.177, 3L = 0.144, 3R = 0.171, X = 0.205). In the Large versus Small *F_ST_* comparison, again there were many regions showing high differentiation (Figure 5B). The chromosome with the highest mean *F_ST_* in the large versus small comparison was the X chromosome (2L = 0.277, 2R = 0.312, 3L = 0.207, 3R = 0.183, X = 0.354). The highest 95% of *F_ST_* windows (window size = 10000bp) for discordant vs. control, as well as large vs. small, were intersected with windows for nucleotide diversity under 0.0005 (windows size = 10000bp) and with SNPs with a CMH (Figure S7; S8) an fdr adjusted p-value less than 0.01. To confirm the effects for SNPs of interest, we modelled allele frequency changes with logistic regression between treatments, examining odds ratio vs. *F_ST_*, verifying high *F_ST_* correlated to large odds ratios (Figure S9; S10; S11). All candidate SNPs were annotated (Supplemental files 1;2;3;4). After manual curation, several genes with known sex-specific size effects were identified including: *dMyc* (*myc*), *Hairless* (*H*), *Insulin-like receptor* (*InR*), *Regulator of cyclin A1* (*Rca1*), and *stunted* (*sun*). Of our candidate discordant genes, 11/295 (excluding inter-genic SNPs, and lncRNA) have both a known size phenotype as well as a sex-limited phenotype. Many other genes with known effects on aspects of body or trait size were also observed for both the discordant and concordant comparisons (supplemental files 1;2;3;4). For the concordant selective lineages, we examined genes in our list overlapping with those identified from generation 100 of selection Turner et al. (2011). The only gene that overlapped between the analysis of generation 100 and 378 was *Nop1-like* (*Nop17l*) in the small vs. large selective treatments. Modulation of expression of *Nop17l* in developing wing tissues reduces size of the wing (Bennett et al. 2006). If we examine overlapping genes excluding the logistic regression analysis, additional genes overlap between F100 and F378 concordant treatments (Supplemental file 5;6).

**Figure 5:**
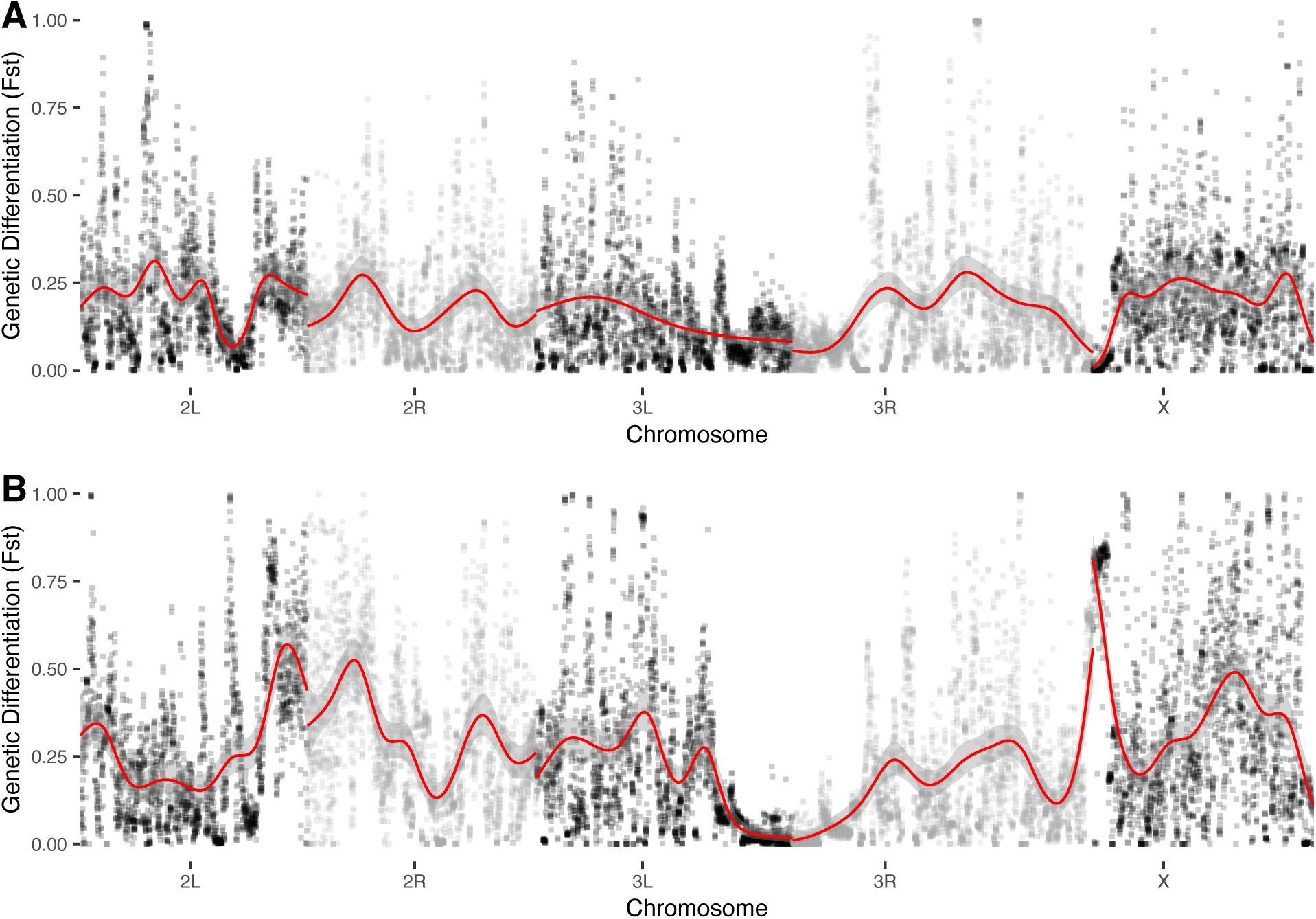
Genome-wide *F_ST_* (10000bp windows). Chromosomal trends for F_ST_ (binomial, gamm) in red. A) Discordant selection compared to control treatments. B) Large compared to small treatments.

### Both biological process and molecular function Gene ontology term enrichment was varied

The candidate genomic regions identified above were then examined for enrichment of GO terms across biological process and molecular function. For the discordant sites of interest, the top biological process GO terms pertained to perception of sweet taste, eye development, and sodium ion transport (Table S6). Within the top 20 GO terms there were also terms associated with development (positive regulation of imaginal disc growth), as well as sex-specific terms (female germ-line stem cell population maintenance). For molecular function, top results included cation:cation antiporter activity, lipid transporter activity, and sweet taste receptor activity (Table S9).

For the large selection treatment, circadian behavior, pupal development, and synaptic growth at neuromuscular junctions were the top biological processes (Table S7). Within the top 20 results, *Wnt* signalling also appeared twice (canonical *Wnt* signaling pathway, positive regulation of *Wnt* signaling pathway). *Wnt* signalling was also in the top 20 molecular function in the large treatment regions of interest (Table S10). The Small treatment regions of interest returned similar GO terms, with both circadian behavior, and pupal development being at the top of the list as well (Table S8). *Wnt* signaling also appeared in the small treatment regions of interest for biological processes as well as molecular functions (Table S11). It is worth noting, permuted GO enrichment analysis with Gowinda did not return significant terms, except for our large treatment which only returned astral microtubule organization as an enriched GO term in our large SNP file.

### Intersexual genetic differentiation in one sexually discordant lineage may suggest the maintenance of intra-locus sexual conflict

Given the long-term nature of the discordant selection, these lineages may be especially useful to identify signatures of IASC. We examined *F_ST_* between males and females within each lineage. As autosomes spend equal time in males and females it is expected intersexual *F_ST_* should be close to zero in most circumstances, absent strong sexual conflict (Cheng and Kirkpatrick 2016; Kasimatis et al. 2019). For all concordant selection treatments (Control, Large and Small), we found mean chromosomal *F_ST_* to be near zero (mean of C1 = 0.0015, C2 = 0.0019, Figure 6A;S12; L1 = 0.0012, L2 = 0.0015 Figure S13;S14; S1 = 0.0023, S2 = 0.0014, Figure S15;S16). For the discordant selection treatment replicate 2, we also find a mean chromosomal *F_ST_*to be near zero (E2 = 0.0011, Figure S17). For replicate 1 of our discordant selection treatment, however, all chromosomal mean *F_ST_* is near zero (E1 = 0.0019) except for an elevated section of ∼3.4Mb on chromosome 3L between position 18100000 and 21600000, where mean *F_ST_* for the rises to 0.005, with elevated SNPs showing a distinct peak of *F_ST_* nearing 0.1, with a couple of windows reaching *F_ST_* of 0.25 (Figure 6B). We did not identify any common samples with inversions on chromosome 3L (*In(3L)P, 3L133in, 3L165in, 3L096, 3L105, and 3L058)*, nor novel structural usingl DELLY (Rausch et al. 2012) contributing to this elevated region of F_ST_. We identified SNPs in this E1 lineage on chromosome 3L that were 3 standard deviations about the mean between sex *F_ST_* (Supplemental file 1).

**Figure 6:**
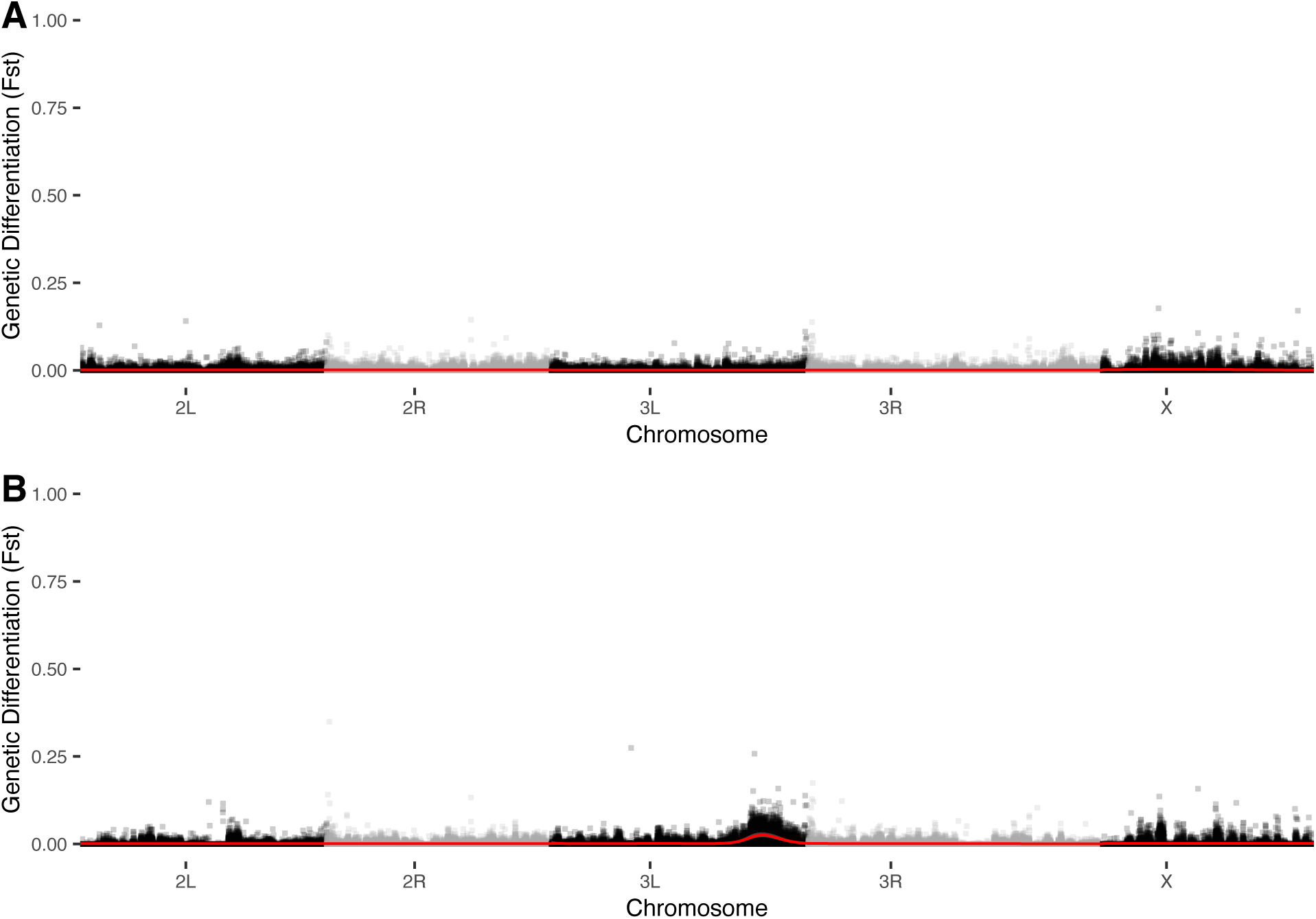
Genome-wide F_ST_ (10000bp windows). Chromosomal trends for *F_ST_* (binomial, gamm) in red. A) Male vs. Female for control treatment (replicate 1), showing a trend ≅ 0, across the genome. B) Male vs. Female for discordant treatment (replicate 1) with a small region on chromosome 3L with elevated *F_ST_* values.

We confirmed that this region of elevated intersex *F_ST_*in discordant replicate 1 was indeed an outlier using custom simulations designed to account for various sources of sampling variation (Figure S18-S25). All male—female comparisons follow a similar pattern to those of the simulations, with the exception of discordant replicate 1, which appears extreme relative to simulated values (Figure S18). We further modelled all SNPs on chromosome 3L to look for clusters of significant SNPs within our elevated *F_ST_* region (Figure S26-S33). Our discordant lineage replicate 1 shows a high number of significant SNPs within our elevated region when modelled (Figure S28), while no other lineage has a clear cluster of significant SNPs in this region. We also looked for genes that overlap with previously identified conflict genes in the establishing population of LH_m_ (Ruzicka et al. 2020), however we found no overlap. If we exclude our modelled SNPs and look for overlap in genes that overlap without identified high *F_ST_* we find a single named gene, Formin-like (*Frl*), which does not have any known body size or sex-limited phenotypes.

### Sex ratio and fecundity may suggest conflict in the discordant lineages genomes

Crossing individuals from all treatment groups in reciprocal single-pair matings to the LH_m_ ‘ancestor’ resulted in ∼1:1 sex ratio in the F_1_. The exception to this was Control replicate 1 males crossed to LH_m_ females (C1: F-M = 0.455, CI = 0.426-0.485; Figure 7), as well as within lineage crosses within both lineages of the discordant treatment (E1: F-M 0.550, CI = 0.517-0.582; E2: F-M 0.550, CI = 0.509-0.589; Figure 7). The deviation in sex ratio in control replicate 1 is also observed in the F2 (C1: F-M = 0.457, CI = 0.427-0.487; Figure 8). The sex ratios of the F_2_s from crosses within both discordant lineages both show male bias, althougwithh the confidence intervals for discordant replicate 1 cross (E1: F-M = 0.557, CI = 0.521-0.592; Figure 8) not overlapping the 1:1 expectation. We also observed a deviation from the expected sex ratio in our large replicate 1 male to LH_m_ female cross (L1: M-F = 0.459, CI = 0.429-0.489; Figure 8), and our discordant replicate 1 male to LH_m_ female cross (E1: M-F = 0.453, CI = 0.426-0.480; Figure 8). This effect in the discordant cross was in the opposite direction of the discordant replicate 1 pure cross.

**Figure 7:**
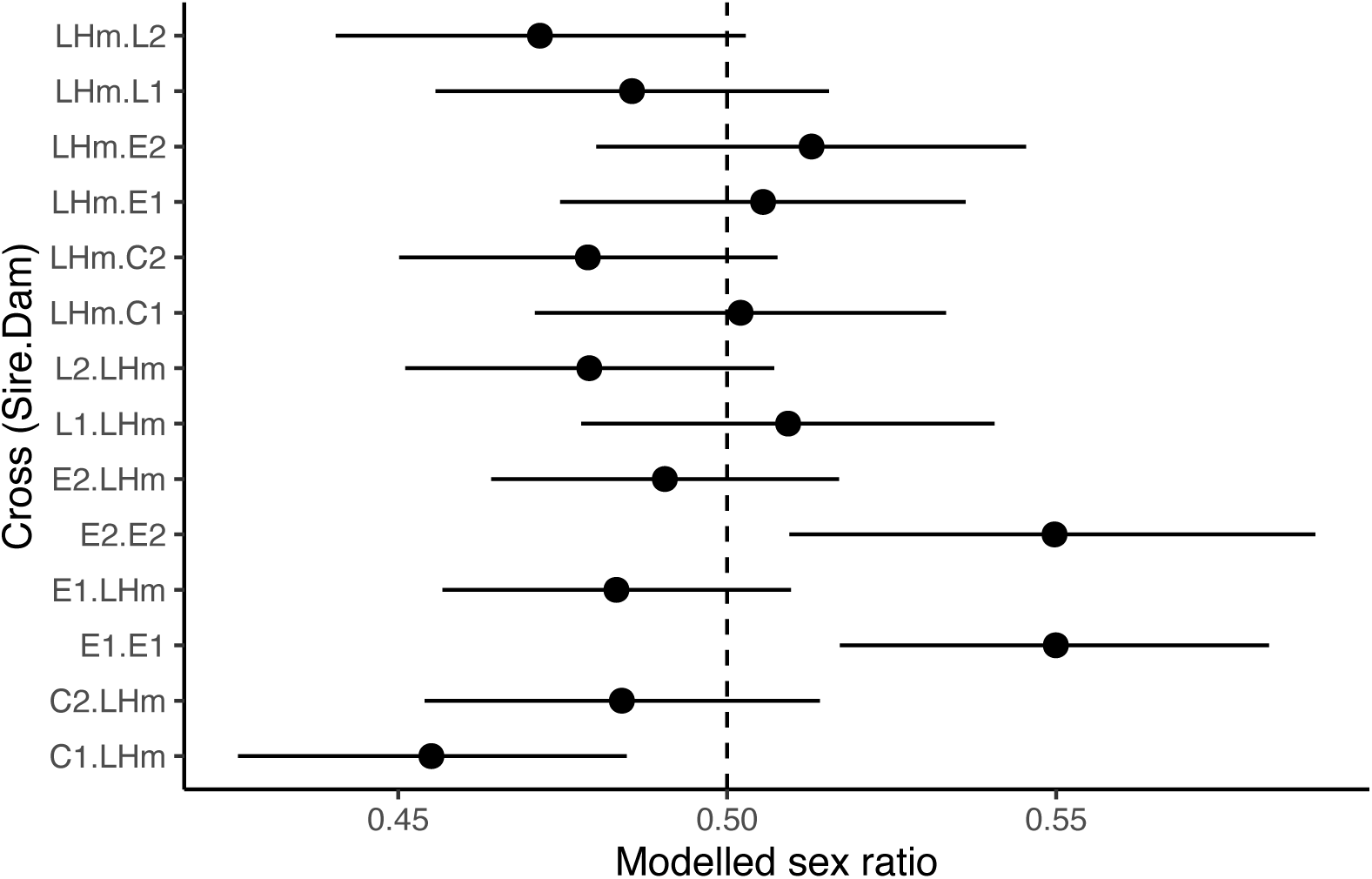
F1 offspring sex ratio from all treatments crossed to the founder population as well as both discordant lineages crossed ‘pure’. Cross label has Sire on the left and Dam on the right of the cross identifier. Dashed line marks expected 1:1 sex ratio.

**Figure 8.**
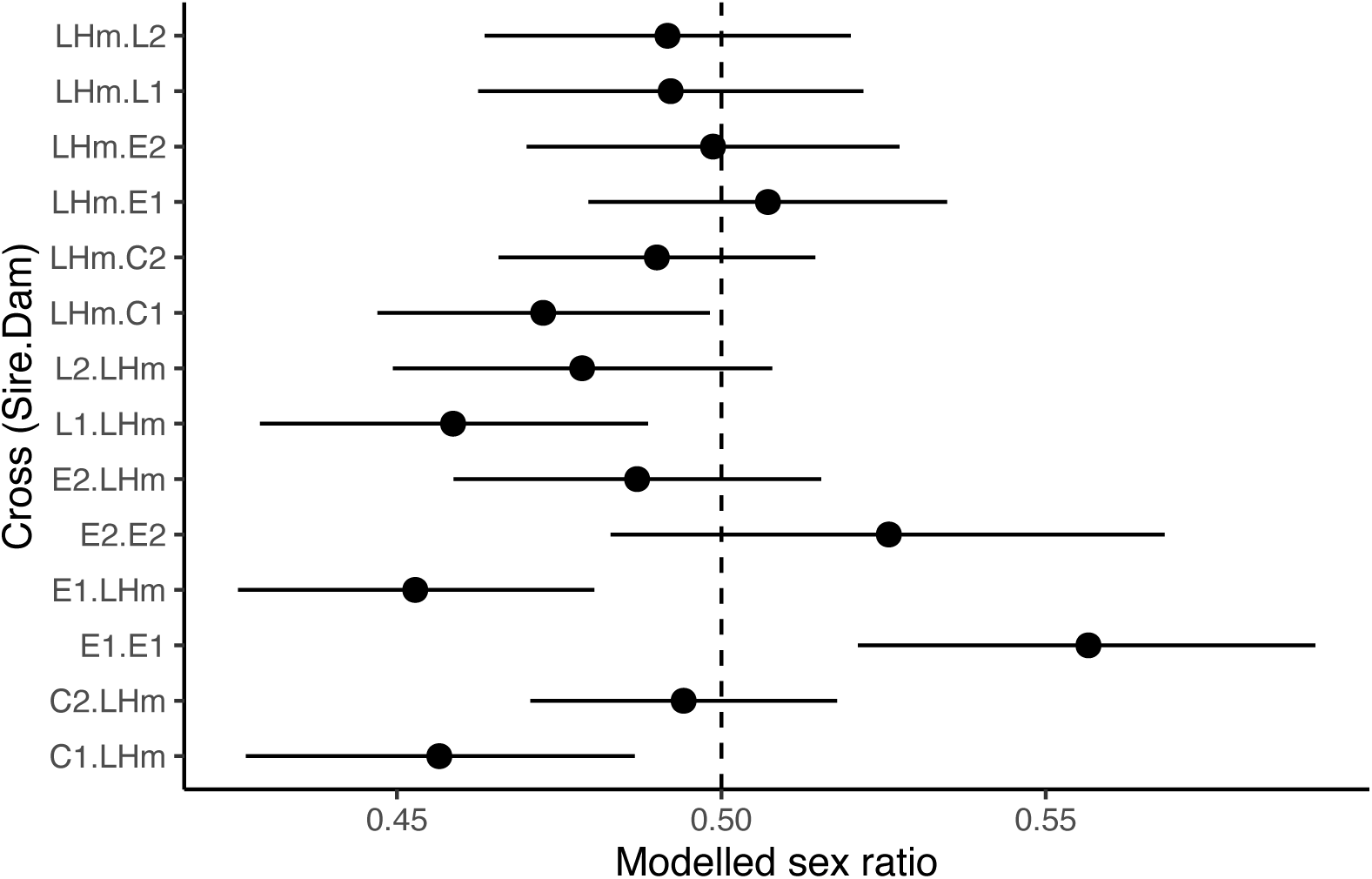
F2 offspring sex ratio from all treatments crossed to the founder population as well as both discordant lineages crossed ‘pure’. Cross label has Sire on the left and Dam on the right of the cross identifier. Dashed line marks the expected 50/50 sex ratio.

We examined whether the direction of the cross influenced the sex ratio, by adding cross direction as a predictor in the model. In the F_1_ generation, overall direction of cross had very modest impacts on sex ratios (χ^2^ = 1.13, df = 1, p = 0.29). There was some evidence of deviation from the expected sex ratio when direction was accounted for when control 1 sired the cross (C1: M-F = 0.455, CI = 0.426 -0.485). In the F_2_ generation, the direction of the F_0_ cross (sire vs dam) had a modest impact on sex ratios (χ^2^ = 5.67, df = 1 p = 0.017). Specifically, when the treatment lineage was the sire in the initial cross for discordant replicate 1, and large replicate 1 (E1: M-F = 0.453, CI = 0.426-0.480; L1: M-F = 0.459, CI = 0.429-0.489). Control replicate 1 crosses appeared to deviate from expected sex ratios regardless of which parent served as sire vs dam.

While it was not a planned experiment, we observed a possible difference in fecundity between treatments. The preliminary results from the analysis from the F_1_, when the dam was from either discordant lineage, showed reduced fecundity (Figure S34). In the F_2_ generation there didn’t appear to be an influence of direction of cross, and discordant lineages had the lowest fecundity (Figure S35).

## Discussion

Sexual dimorphism (SD) evolves frequently, despite *r_MF_*generally being high within species, in particular for morphological traits (Lande 1980; Poissant et al. 2010). In the presence of sex-specific optimal phenotypes, this *r_MF_*has the potential to generate intra-locus sexual conflict through the ‘load’ on the opposite sex (Fairbairn et al. 2007). Species that evolve changes in SD must overcome any hurdles due to a high *r_MF_*as well as ensuing genomic conflict. In the long-term, this conflict may reach an equilibrium, with sex-biased alleles creating as near an optimum phenotype for each sex as possible or be resolved entirely. However, the reintroduction of sex-discordant selection should disrupt this equilibrium and generate additional genomic conflict. Examining the response to sexually discordant selection for body size across the genome after selection provides an opportunity to identify genomic regions undergoing conflict. In this study we utilize long-term artificially selected lineages (Stewart and Rice 2018), selected for body size either in sexually concordant or discordant manner. As discussed in detail below, in addition to observing trait specific changes in patterns of sexual size dimorphism, we see potential evidence for the maintenance of polymorphisms consistent with unresolved conflict.

### Concordant selection lineages maintained ancestral patterns of SSD despite selection on size, while discordant selection lineages responded in a trait specific manner

The lineages under concordant artificial selection for size responded in the expected directions, for all traits, and changed size proportionally maintaining SD for all traits. This proportional response is consistent with relatively high *r_MF_* for morphological traits between the sexes, as previously observed (Cowley and Atchley 1988; Reeve and Fairbairn 1996). If *r_MF_* had been low in the starting population, we may have observed variation in responses between the sexes, with one sex responding to selection faster than the other, and the overall SSD changing from the ‘baseline’, as has been shown when *r_MF_* is intentionally degraded (Delph et al. 2011).

Instead, within concordant treatments, SSD was largely maintained despite substantial changes in size, in particular, under selection for smaller body sizes. For the discordant artificial selection treatment, SD changed trait-specifically (Figure 2). Of the four traits measured for this study, one (thorax length) showed a substantial degree of female biased dimorphism in control lineages (difference of ∼0.16, in log_2_ μm, or 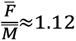), while leg traits (femur, tibia and basi-tarsus lengths) were closer to sexually monomorphic (differences of 0.029, 0.023, 0.074, or ratios of 1.02, 1.02, 1.05 respectively). Under discordant artificial selection, all traits saw a reduction in female biased dimorphism, with the magnitude of change varying by trait (Figure 3).

Dimorphism for thorax length reduced to the greatest degree (difference of ∼0.012, in log_2_ μm, or 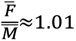), compared to dimorphism for leg lengths (-0.076, -0.087, 0.011, or ratios of 0.95, 0.94, 1.01). This is perhaps most clearly illustrated in the changes in patterns of multivariate allometry observed in the sexually discordant lineages (Figure 4). Traits with higher levels of sexual dimorphism tend to have lower *r_MF_* than non-dimorphic traits (Poissant et al. 2010), which may partially account for this. However, this may also reflect the manner in which artificial selection was applied (via a sieve), which likely results in a complex composite “size measure” for selection on cross-sectional area, of which a trait like thorax length may contribute substantially more than any individual leg measure. Alternatively, the genetic correlations between the sexes may differ across traits. This might explain why traits like femur and tibia reversed sexual dimorphism when selected on indirectly, however, tarsus and thorax (thorax being the most directly selected on) did not fully reverse dimorphism.

### Within-lineage, across-sex comparison provides possible evidence of intra-locus sexual conflict in the genome of one discordant lineage

To identify regions under possible intra-locus sexual conflict we compared male-female *F_ST_* within each lineage, and conducted several follow up analyses, to identify possible SNPs showing subtle, intersexual, within-generation distortions in allele frequency. There may be small genomic regions or possibly single SNPs showing signs of different allele frequencies between the sexes, which may be evidence of unresolved conflict in the genome. Neither control lineages, nor the sexually concordant lineages showed any evidence for elevation in *F_ST_* (Figure 6A;S12-S17). However, one discordantly selected lineage (E1; Figure 6B), has a region on chromosome 3L with *F_ST_* reaching as high as >0.2. Other comparisons of this nature have yielded smaller male-female *F_ST_* estimates, which led us to evaluate whether it was an artifact of our pipeline (Cheng & Kirkpatrick 2016; Lucotte et al. 2016; Flanagan & Jones 2016; Dutoit et al. 2017; however see Sylvestre et al. 2023). Modifying SNP calling methods, *F_ST_* estimation procedures, and filtering methods all yielded the same spike in *F_ST_* in the E1 lineage. This elevated region of allele frequency differentiation was also extreme for only this lineage in comparisons to simulations accounting for a number of sources of sampling variation and overall allele frequencies differences. If it was a spurious peak of elevated *F_ST_* as a result of technical aspect of the pipeline, or mis-mapping of reads from a sex chromosome, other replicate lineages would likely also show elevated regions. We consider the most likely explanation to be due to the sustained nature of sexually discordant selection in our experiment. Compared with most other studies examining M-F differences in allele frequencies to identify potential loci under conflict, the selection applied to the discordant lineages in this experiment is both strong, and sustained across hundreds of generations. Kasimatis et al. (2019) suggests that for a locus to show this degree of asymmetry it would require an exceedingly high magnitude of selection. In their model, this would require near 40% mortality each generation to maintain the distortion in M-F allele frequencies at a locus. The nature of artificial selection as applied in the current experiments means well over 40% (∼80% of individuals with the current selection design) of flies each generation (those not at phenotypic extremes) are discarded, essentially ‘dead’ in the selective sense. Alternatively, Kasimatis et al. (2019) also simulated subsamples of 50 males and 50 females with antagonistic selection and found that sampling substantially increased variability in *F_ST_*values due to the fact that genomes under antagonistic selection tend to have increased numbers of intermediate frequency alleles. However, our simulations suggest this is not a likely explanation for the observed results (Figure S18-S25). Why this region in discordant treatment replicate E1 shows this distortion, and the contribution of intra-locus conflict, requires further exploration.

Given the strange peak in this one discordant replicate, we extracted all SNPs three standard deviations above mean *F_ST_* on chromosome 3L, that also were significant in the logistic regression (which better accounts for sampling variance). We filtered for SNPs that appear in both discordant replicates and do not occur in any concordant lineage to look for possible replicated mechanisms of divergence in phenotype. We found fewer candidate genes than in any other comparison (Supplemental file 1), but multiple genes present in the WNT pathway, including *frizzled2* (*fz2*). We also found candidate SNPs in the MAPK pathway. Interestingly, none of the male-female discordant treatments SNPs showed evidence for genetic differentiation between artificial selection treatments. Similar to other comparisons however, many candidate genes have mutational phenotypes listed on flybase as “abnormal size” so this could be suggestive again that response to selection is highly polygenic with alleles of small effect “inching” towards a phenotype rather than “sprinting”.

Finally, we explored genes found by (Ruzicka et al. 2019), identified as potentially under sexual conflict. These genes were considered in conflict based on sex-specific fitness (competitive mating success in males and competitive fecundity in females). An important caveat in our comparison is that although our lineages are from the same initial lab population (LH_m_), our treatment groups have gone through nearly 400 generations of strong selection from the ancestral population. It is also unknown how many generations of divergence occurred in the ancestral LH_m_ populations prior to both experiments. In our comparison between genes identified in Ruzicka et al. (2019) those identified in our study, only the gene Formin-like (*frl*) overlaps. *Frl* is upstream of our region of interest and does not have any previously identified body size or sex-specific phenotypes. We may not see overlap because Ruzicka et al.(2019) identified conflict alleles based on specific proxies of fitness, whereas ours may reflect a more indirect effect due to discordant selection on size. However, we do observe some evidence that adult sex ratios are male skewed in the discordant lineages, and possibly maintained even in crosses to the LH_m_ “ancestor”, as well as potential reductions in fecundity (Figure 7, 8, S34, S35).

### Discordant size selection likely has a polygenic basis, involving a number of genes that influence body size in a sex-specific manner

Given the number of generations of artificial selection, and the modest population sizes each generation, genetic drift will have an overwhelming impact on any genome scan performed with the evolved lineages. As such, any interpretation of these genome scans should be tempered by this impact. With that being said, the response is broadly consistent with a polygenic genetic basis, with thousands of SNPs showing evidence of genetic differentiation across the genome. In 378 generations of selection and more than 125 generations after clear evidence of phenotypic reversal of SSD, the opportunity for a mutation (or ancestrally segregating, relatively rare variants) of large phenotypic effect occurring was possible. However, we find no clear evidence for alleles of large effect that have swept to fixation in parallel across both lineages of discordant selection. The chance of identifying alleles of large effect is attenuated by the impact of drift making many SNPs fix randomly in each lineage. Possibly constrained by genetic correlation, the response to discordant selection appears to be a slow and steady crawl towards a response rather than large leaps forward. The lineage specific increase in discordant replicate 1 that does not appear in replicate 2 could imply a novel mutation that is under conflict in this replicate. This elevated *F_ST_* is shown in a large region of chromosome 3L that includes hundreds of SNPs however, so to identify a novel mutation responsible requires further investigation. This slow crawl towards a response is supported by the fact that Stewart and Rice (2018) saw a negligible phenotypic response to discordant selection during at least the first 100 generations, with the more substantial responses occurring after this. Large effect alleles with sex-specific impacts on size have been identified in functional studies, however the viability cost in the opposite sex may make these large effect alleles prohibitive (Rideout et al. 2015; Millington et al. 2021a). Although it is tempting to speculate that these small effect alleles are a likely way for SSD to evolve in natural populations, we must also caveat that it may be the case that these alleles of such small effect may be weeded out by viability or fecundity selection in the wild, which are much weaker in this tightly controlled artificial selection experiment. In fact, Testa and Dworkin (2016) demonstrated that mutations with a sex-specific response are rare in two distinct genetic backgrounds. Carreira et al. (2009) however, found that many genes associated with size change reduced SD when a P-element was inserted. This highlights the complexity of the genetic architecture of SSD.

Although Testa and Dworkin (2016) found that mutations with sex-specific effect on size were rare, they did identify mutations in the EGFR pathway with sex-specific effects. In our sex-discordant lineages we found a number of genes with known sex-specific size effect (d*Myc, H, InR, RcA1, sun*). Follow up on these genes to determine the role they play in response to discordant selection is required. Interestingly, we also found a number of enriched GO terms for short term memory, circadian rhythm, and nervous system functions, which don’t have any clear link to body size, but natural populations varying in body size also found to have enrichments for GO terms associated with memory & learning, as well as clock genes (Fabian et al. 2012). This could be further supporting that response to selection is very polygenic and involves a number of, likely highly pleiotropic genes. Other GO terms that were enriched were related to gustatory responses such as sweet taste and response to nutrients which may suggest that a response in how larvae are responding to their environment (eating more) may play a role in size changes, but this requires more follow up.

We also compared out genes of interest to the previously published gene list from Ruzicka et al. (2019) mentioned above. In our comparison we still find modest overlap (11 named genes and 8 unnamed genes). Two genes in particular stand out however, discs large 1 (*dlg1*) which has a known increased size phenotype as well as courtship and oogenesis phenotypes, and *smrter* (*smr*) which has known decreased size phenotypes as well as a sex-limited reduced fecundity phenotype in females. Although the caveats of this comparison mentioned above still stand, it is interesting that we find genes under potential conflict coming up in these two studies both using LHm. Optimistically, this could suggest genes under conflict are maintained in the population and are being captured in both studies exploring sexual conflict, but more work must be done to confirm this. Both of these genes are located on the X chromosome which has been suggested to be enriched in conflict alleles (Rice 1984).

### Concordant selection lineages show a response in growth pathways, but do not align with previous analyses

The concordant selection lineages (small and large) showed SNPs of interest in multiple growth-related pathways. Interestingly, we found no clear overlap with analyses done nearly 300 generations prior, by Turner et al. (2011). This result could be due to a few possibilities: 1) inconsistencies with bioinformatics tools and pipeline choices, 2) improvements in sequencing and software refining our more recent search, 3) potentially most biologically interesting, this could be due to multiple ‘soft’ selective sweeps generating allelic turn-over phenomena in the genes under active selection for size. To circumvent the first possibility, we re-analyzed the F100 data used by Turner et al. (2011) with our pipeline, adjusted for coverage and quality. Importantly (and not surprisingly given when the experiment was done), the sequence coverage per sample is very modest (coverage ∼25x) in comparison to our current study (∼200x). Using our pipeline, the F100 data had no genes with known size phenotypes overlap with our large genes of interest (Supplemental File 5). In the gene list overlap with our small treatment genes of interest, we retrieved Mnt (*mnt*), which has a known increased body size phenotype, we also identified saxophone (*sax*), which has an abnormal size phenotype, and potentially interestingly we recovered Tousled-like kinase (*tlk*), which has a known decreased body size phenotype (Supplemental file 6). The fact that we recover overlapping size related genes in the small treatment, but not the large treatment is likely due to the stronger selection in this treatment.

Although our candidate genes differ from those listed by Turner et al. (2011), both analyses found genes in related general pathways such as ecdysone signalling and the EGFR pathway. This could be evidence that the standing genetic variation present in the starting population (LH_M_) could be fueling the clearly polygenic response and new mutation or stochastic changes in allele frequency are continuously altering which genes are used to response to the selection over such a large time frame. The pathways both we and Turner et al. (2011) identify have been previously implicated as being under selection in naturally occurring clines in *D. melanogaster*, and therefore may provide interesting insight into natural variation in body size. We identified a substantial number of genes after conservative filtering (126 in Large and 101 in Small) that appeared to be under selection in our concordant selection lineages. A number of these genes have known size phenotypes, however since such a large number of genes in the genome impact body size, it is difficult to say if these are directly responsible for response to selection or are mere coincidence (Carreira et al. 2009; Turner et al. 2011). Additional experimental work will be necessary to distinguish between these possibilities.

Our results are both an important exploration of genomic conflict using artificial selection, as well as a follow up on one of the longest (in generations) artificial selection experiment in animals. Although with such strong and persistent selection, our results may not mirror natural settings, it does demonstrate one possibility of how the genome may respond to discordant selection. We also manage to demonstrate a likely differentially segregating region of the genome in a discordantly selected lineage, possibly a distinct autosomal region of sexual conflict in the genome. Although these works require further experimentation to narrow down and validate alleles of interest, we suggest that we have demonstrated at least one route for the genome to respond to discordant selection and sexual conflict.

## Author contributions

Study Conceptualization and funding: ID

Study Design: ID

Artificial Selection and rearing: ADS

Dissections: JK

DNA extraction: TA

Image Analysis: JK, ID

Analysis: ID, TA, KP

Manuscript drafting: TA, ID

Manuscript editing: TA, ADS, ID

Manuscript revisions: TA, ID

## Conflict of Interest statement

The authors declare no conflict of interests.

Upon acceptance all data and scripts will be made available via github as well as either Figshare or DRYAD. All raw sequence data will be deposited in NCBI.

## Acknowledgement and Funding

This work was funded by an NSERC Discovery grant to ID, and an Ontario Graduate Scholarship to TA.

---

**Table S1:**
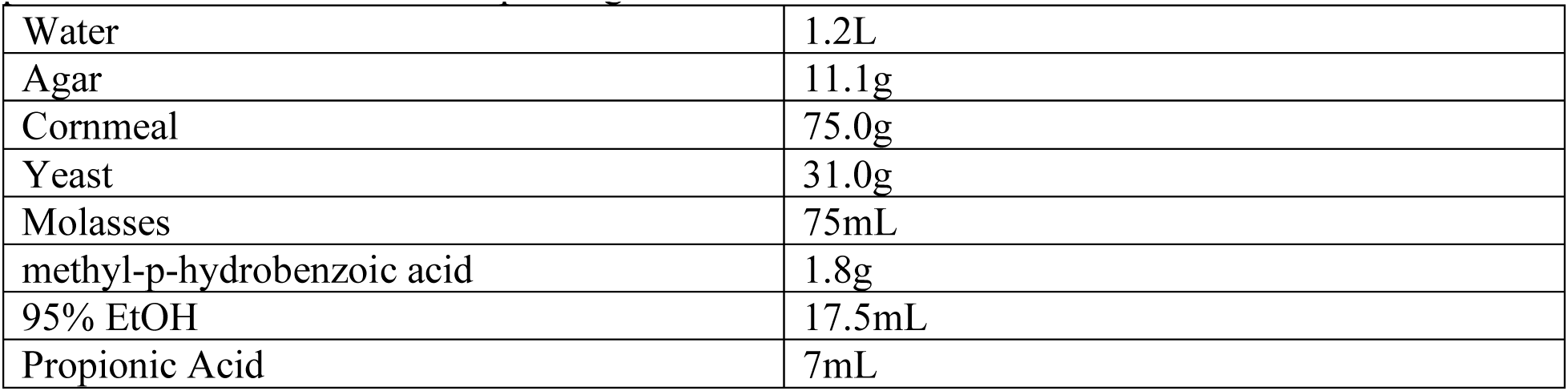
Food recipe for artificial selection lineages. All ingredients except cornmeal, yeast, and preservatives are brought to a boil for 10 minutes, yeast/cornmeal and a small amount of reserved water are added and boiled for another 10 minutes, food is cooled to 60°C then preservatives are added before pouring ∼10mL in to vials.

**Figure S1:**
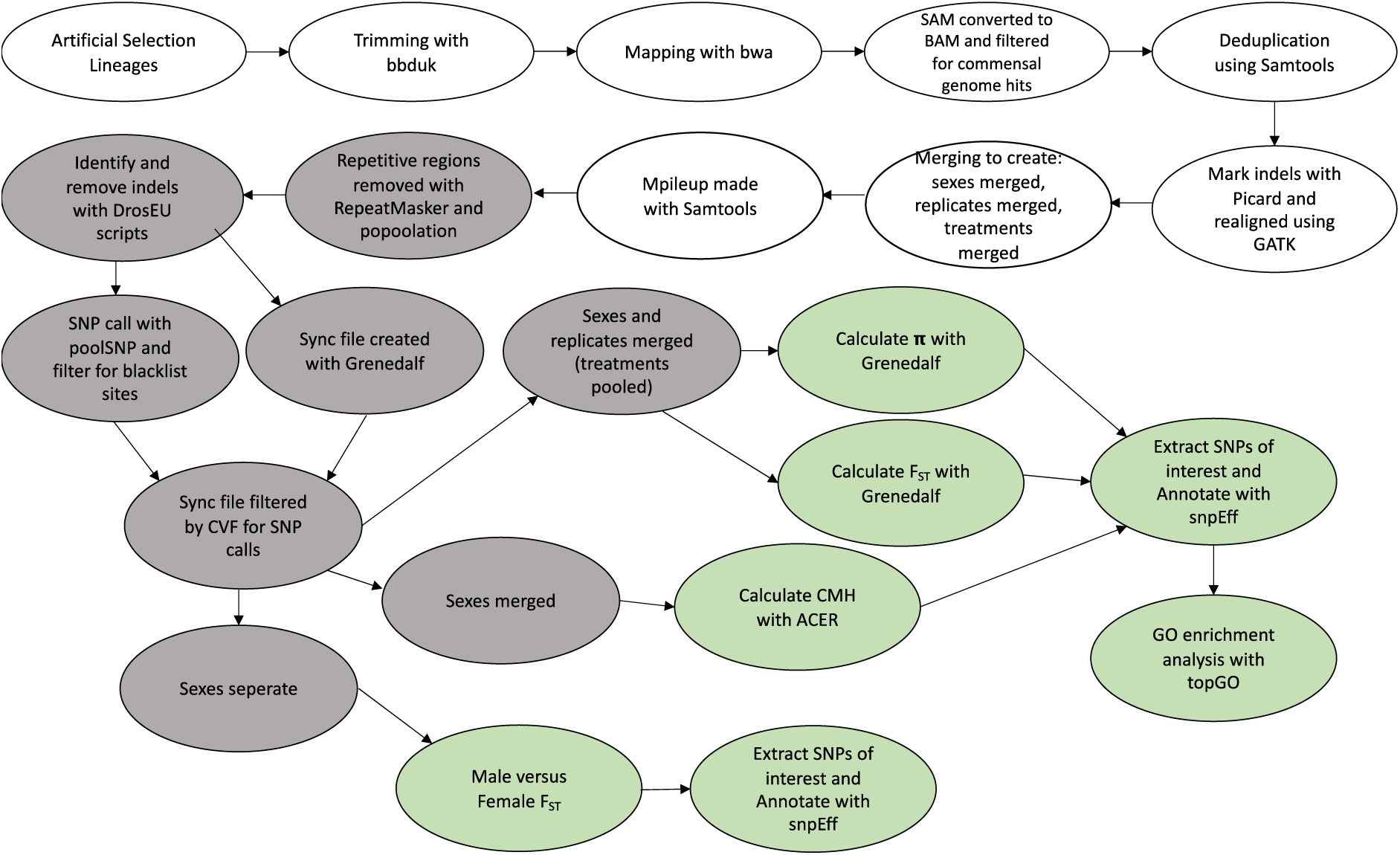
Flow chart showing steps and tools involved in the bioinformatic pipeline. Version numbers, parameter settings and more detail available in text. White: bam files, grey: intermediate files, green: output results files.

**Table S2:**
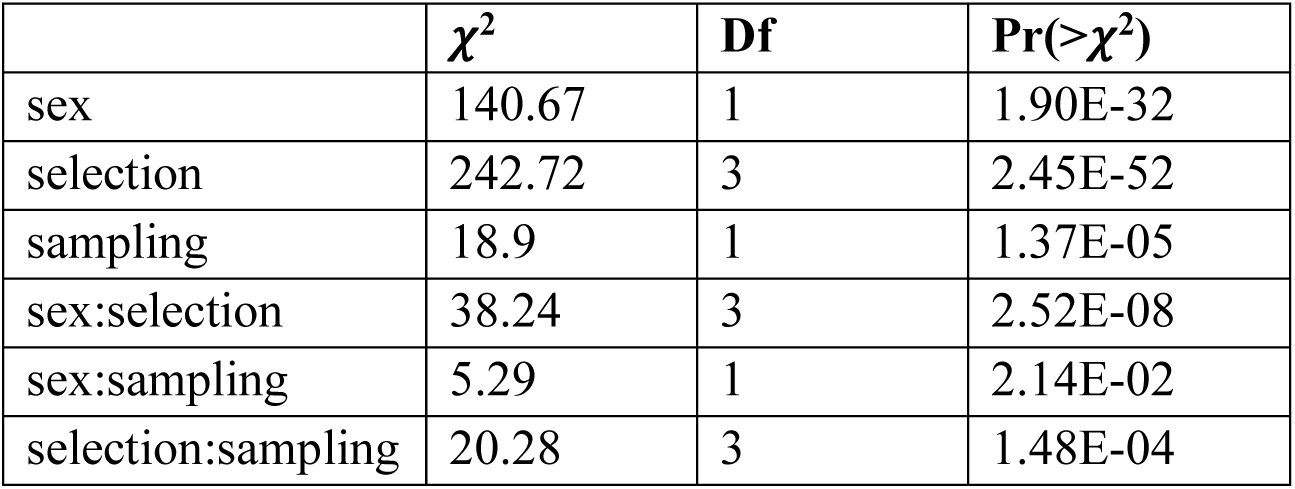
model ANOVA for thorax.

**Table S3:**
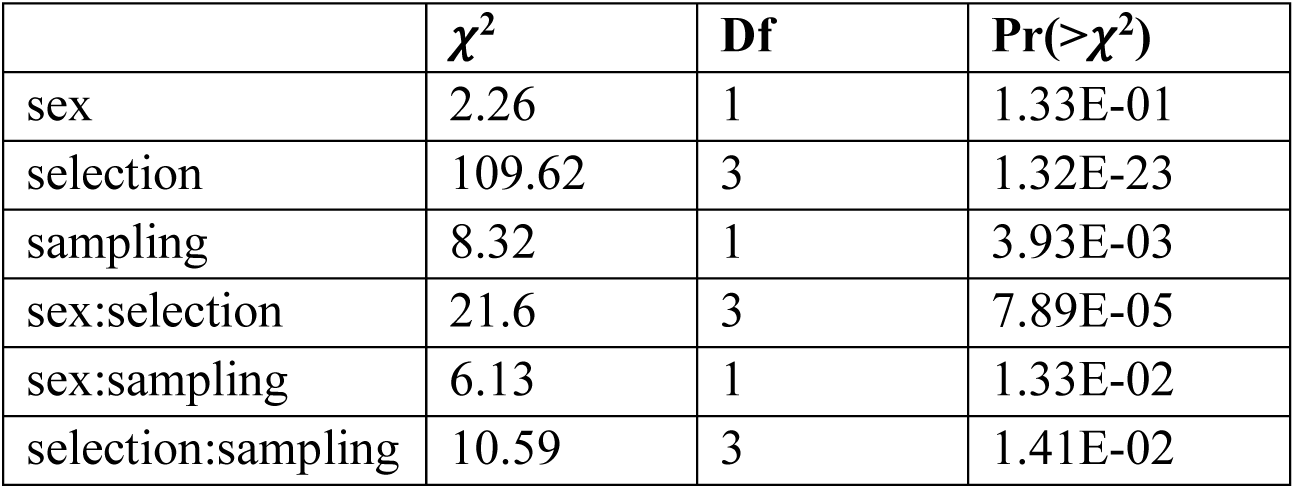
model ANOVA for femur.

**Table S4:**
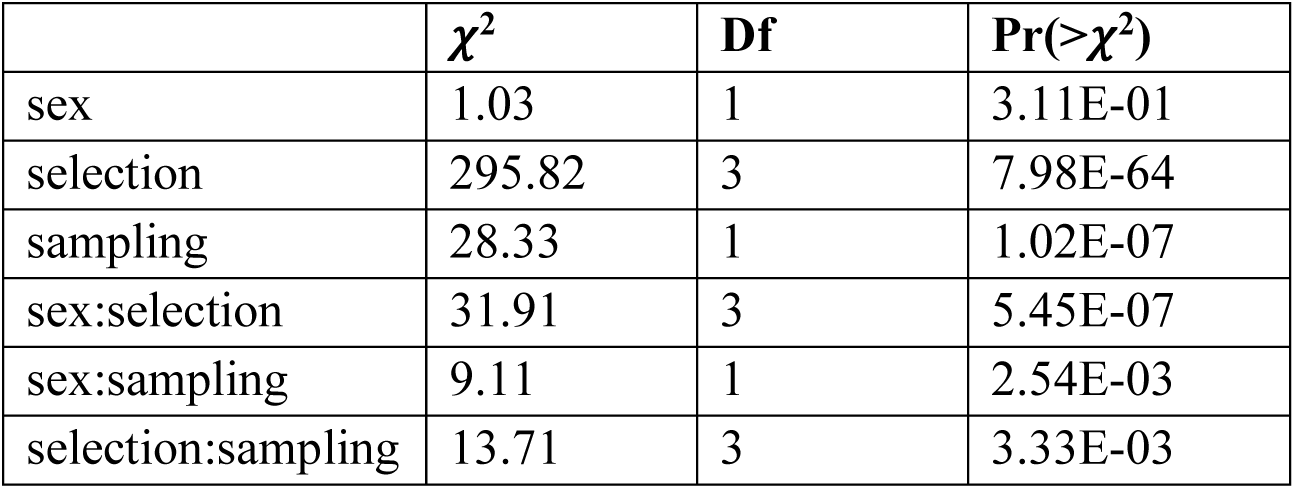
model ANOVA for tibia.

**Table S5:**
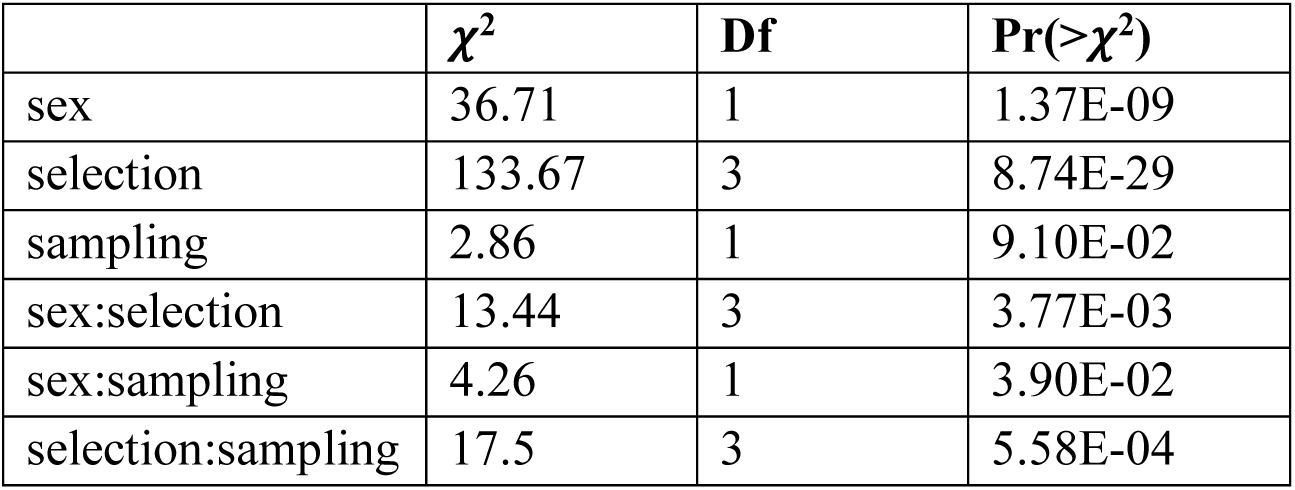
model ANOVA for tarsus.

**Figure S2:**
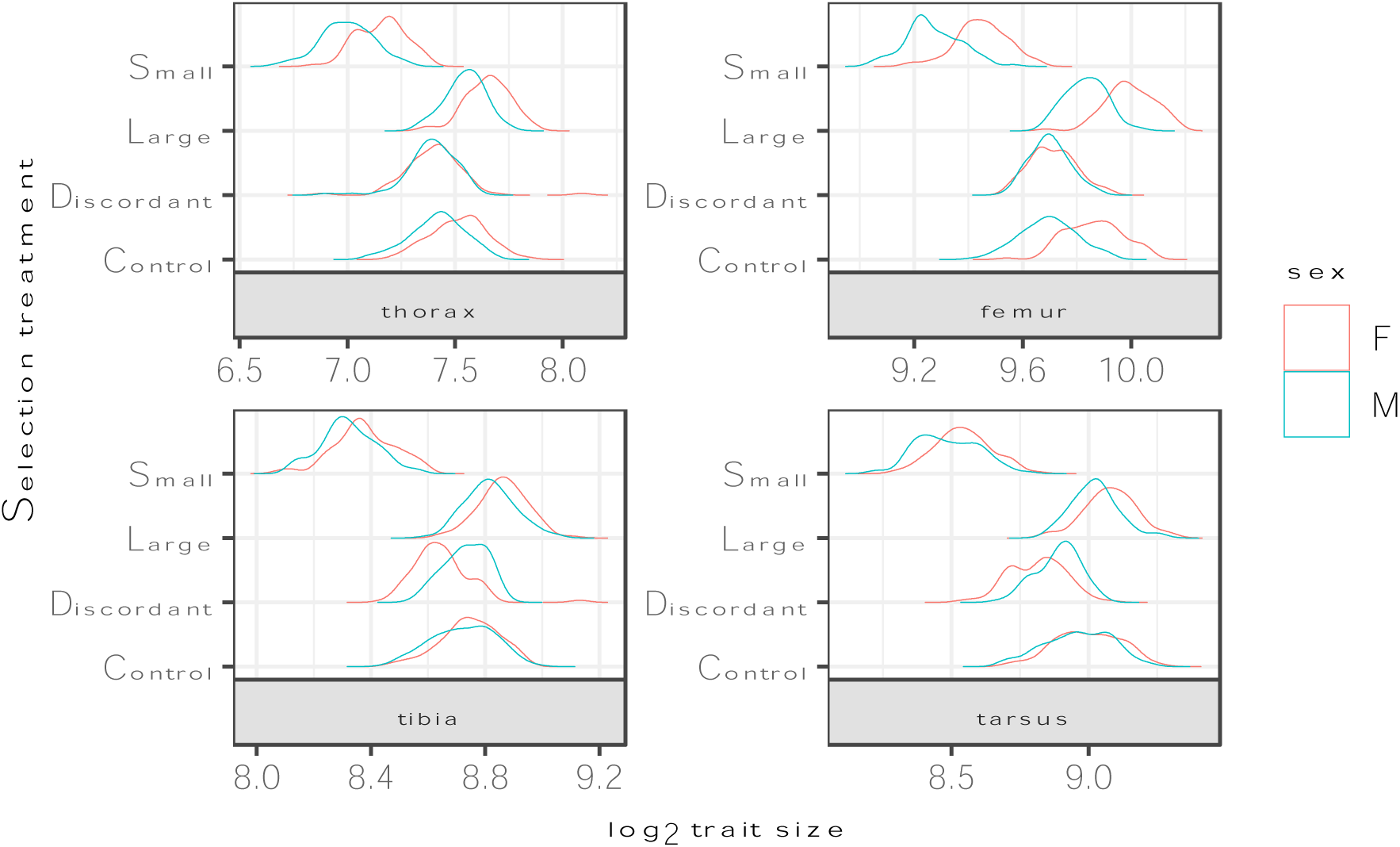
Density plots of raw measures (*log_2_* transformed) for traits and sexes

**Figure S3:**
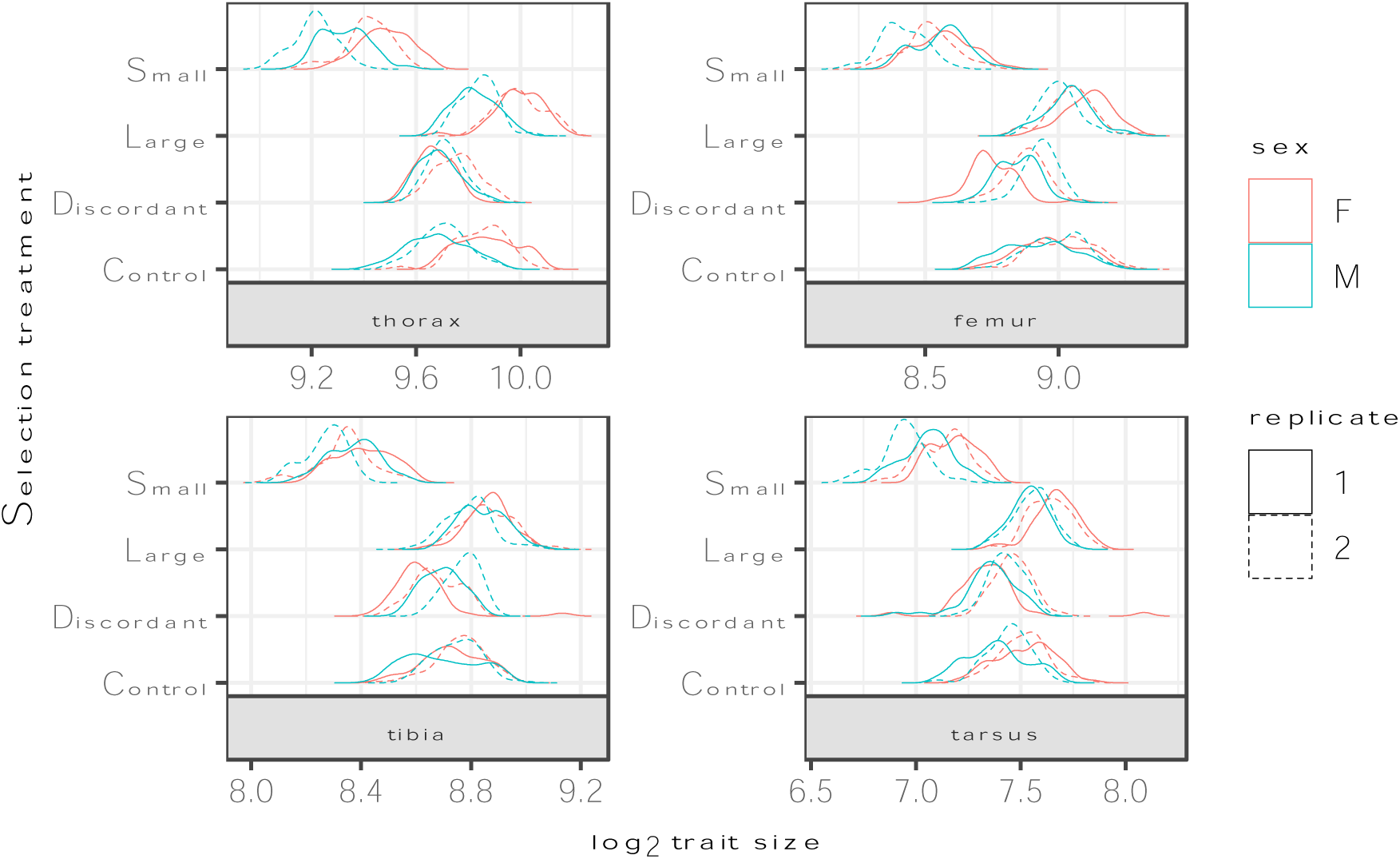
Density plots of raw measures (*log_2_* transformed) for traits and sexes with replicates separated to show replicate heterogeneity

**Figure S4:**
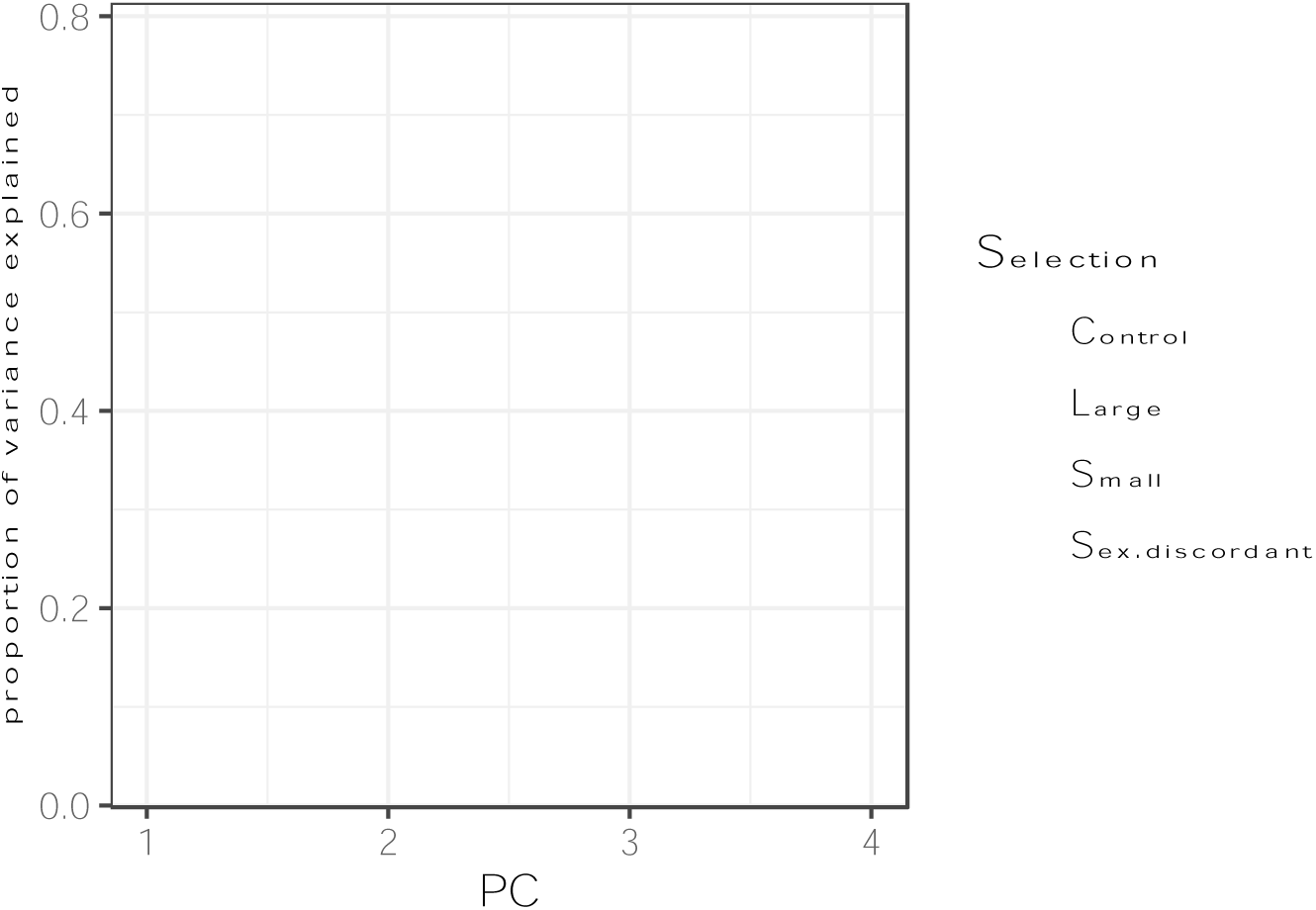
scree plot of proportion of variation explained by each associated eigenvector.

**Figure S5:**
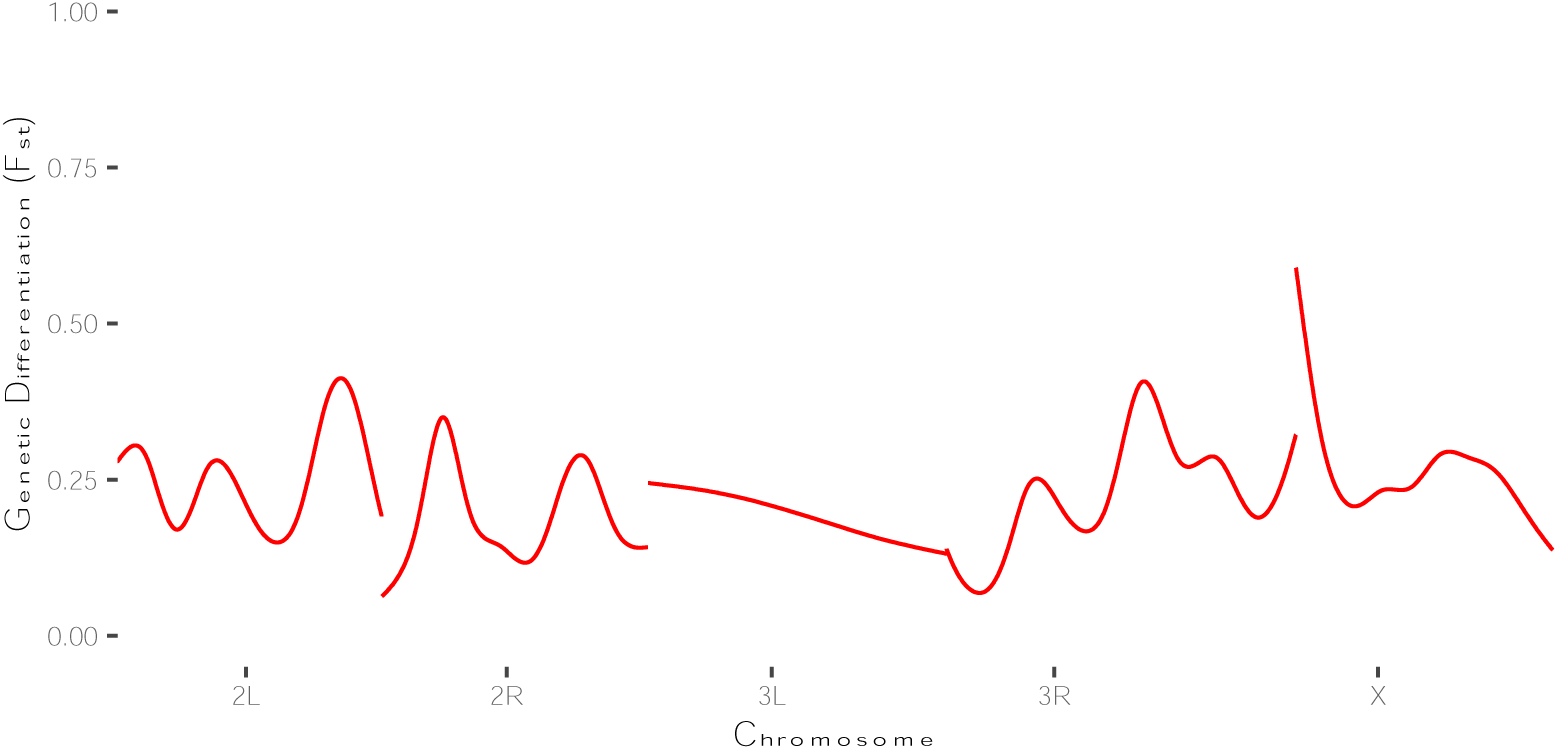
discordant vs. large 10000bp windows

**Figure S6:**
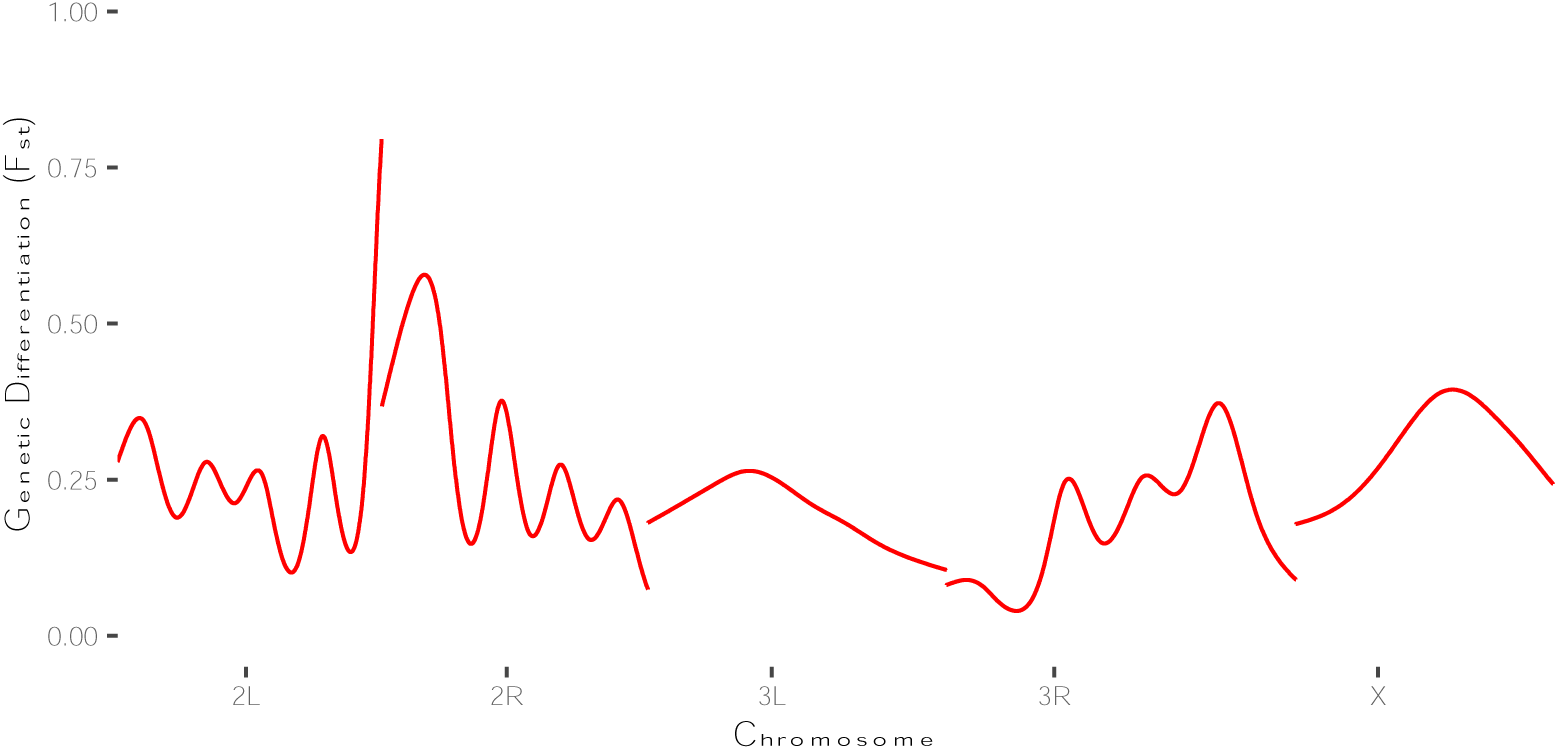
discordant vs. small 10000bp windows

**Figure S7:**
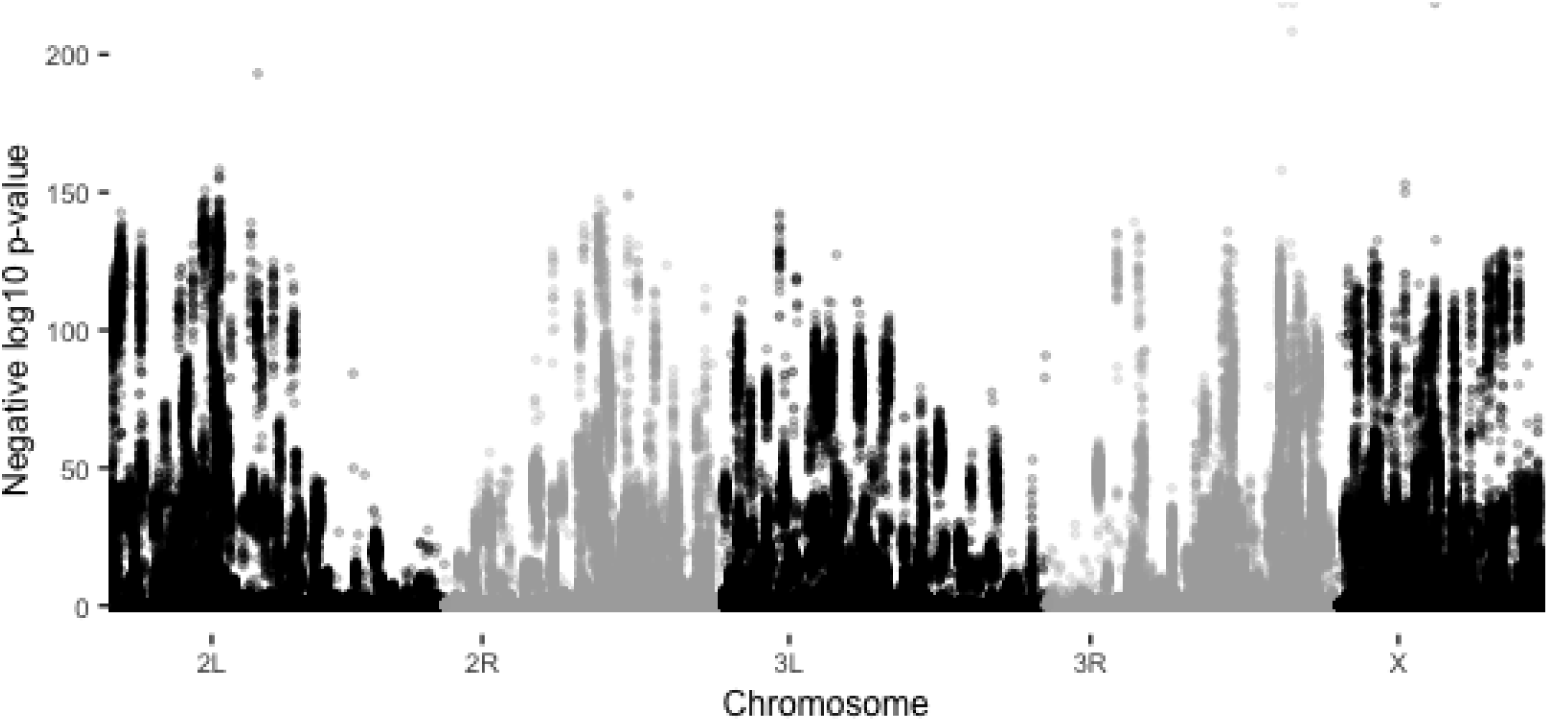
CMH p-values for Large vs. Small treatments

**Figure S8:**
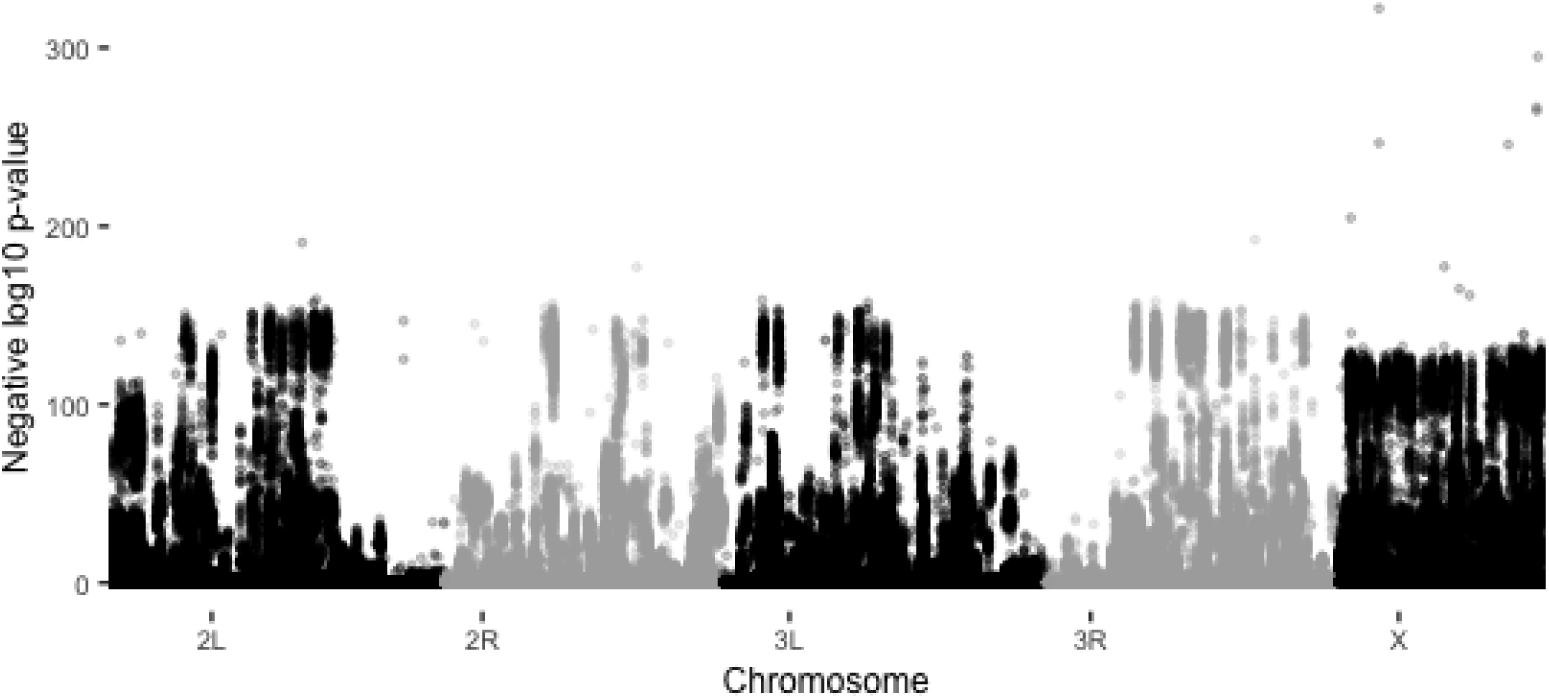
CMH p-values for discordant vs. control treatments

**Figure S9:**
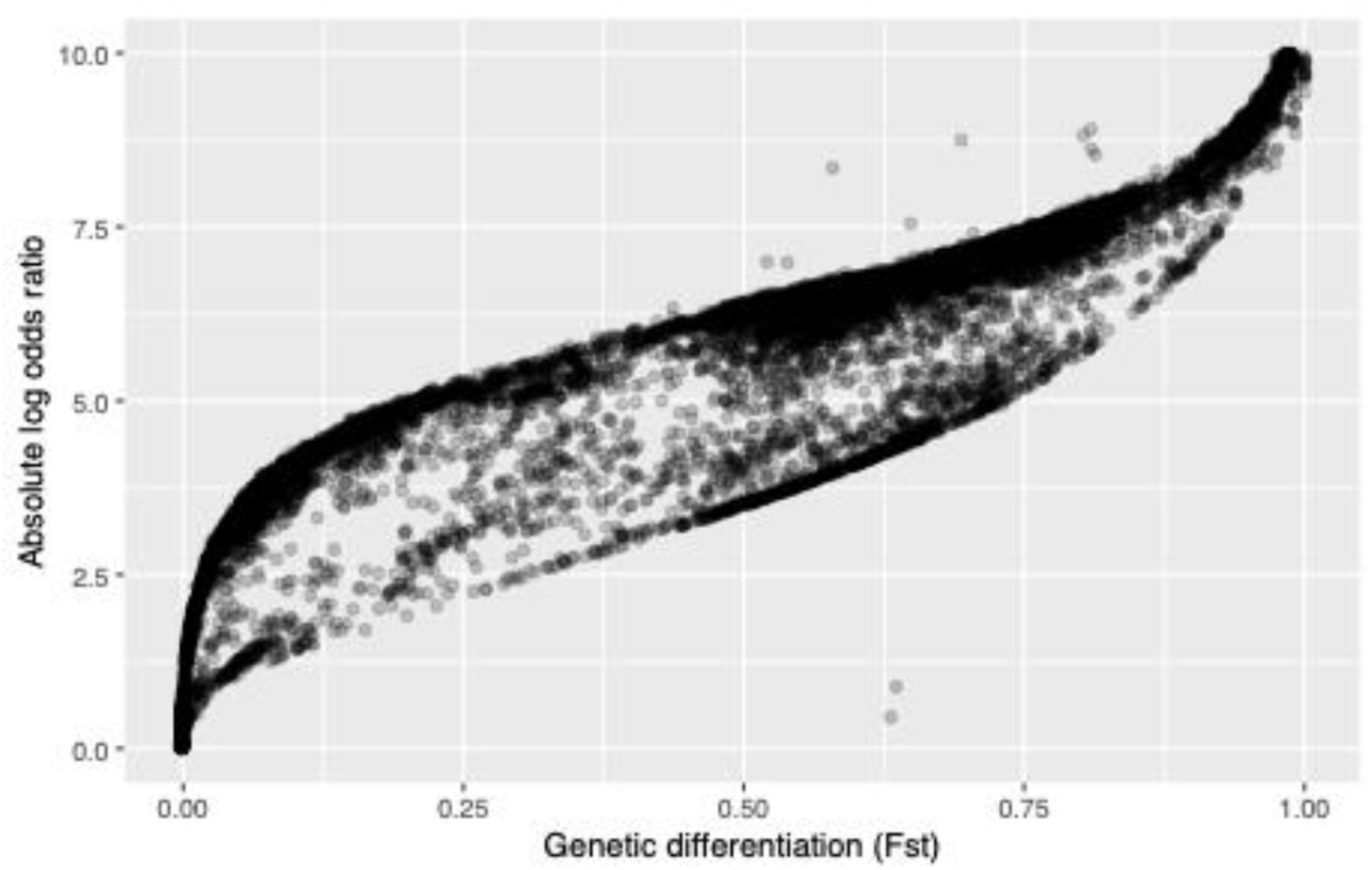
Correlation between F_ST_ and the log odds from our model estimates for the Discordant vs. control comparisons

**Figure S10:**
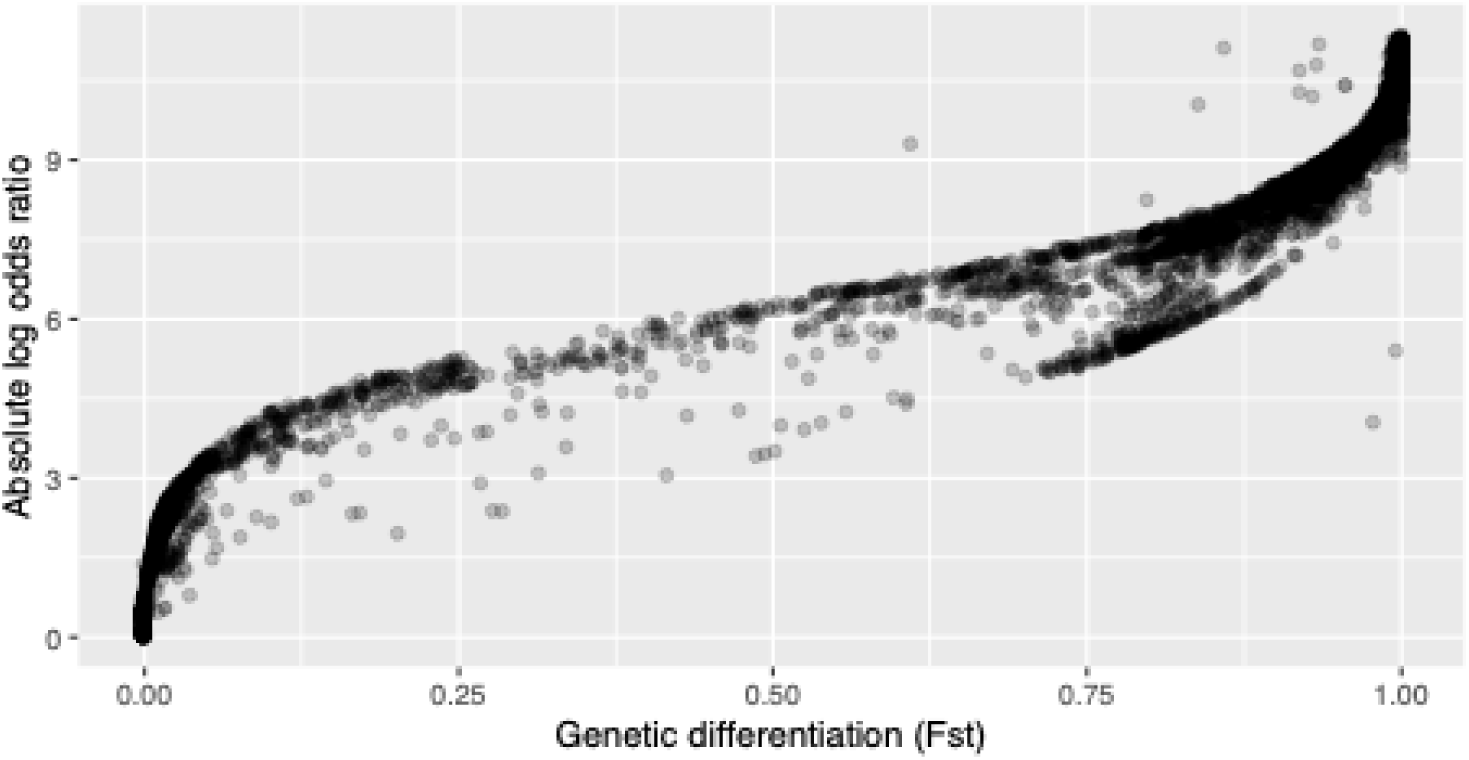
Correlation between Large vs. Small F_ST_ and the log odds from our model estimates for the SNPs found to be associated with the Large treatment

**Figure S11:**
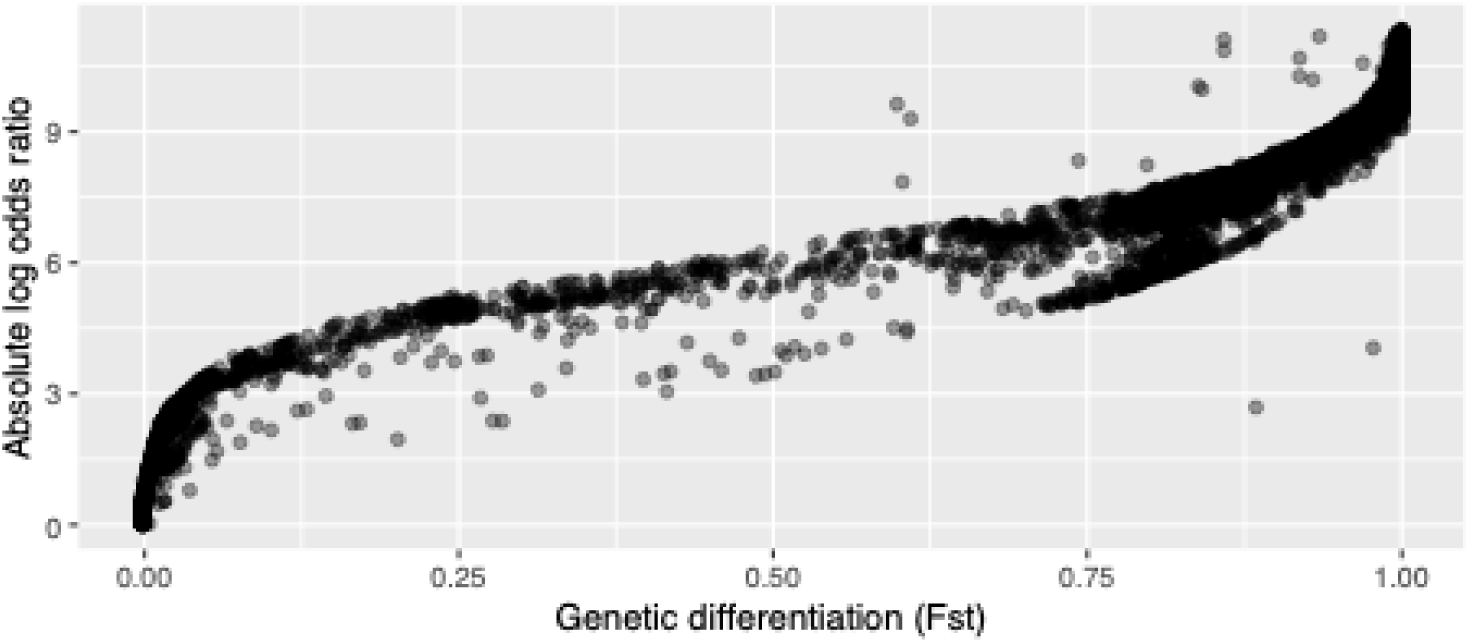
Correlation between Large vs. Small F_ST_ and the log odds from our model estimates for the SNPs found to be associated with the Small treatment

**Figure S12:**
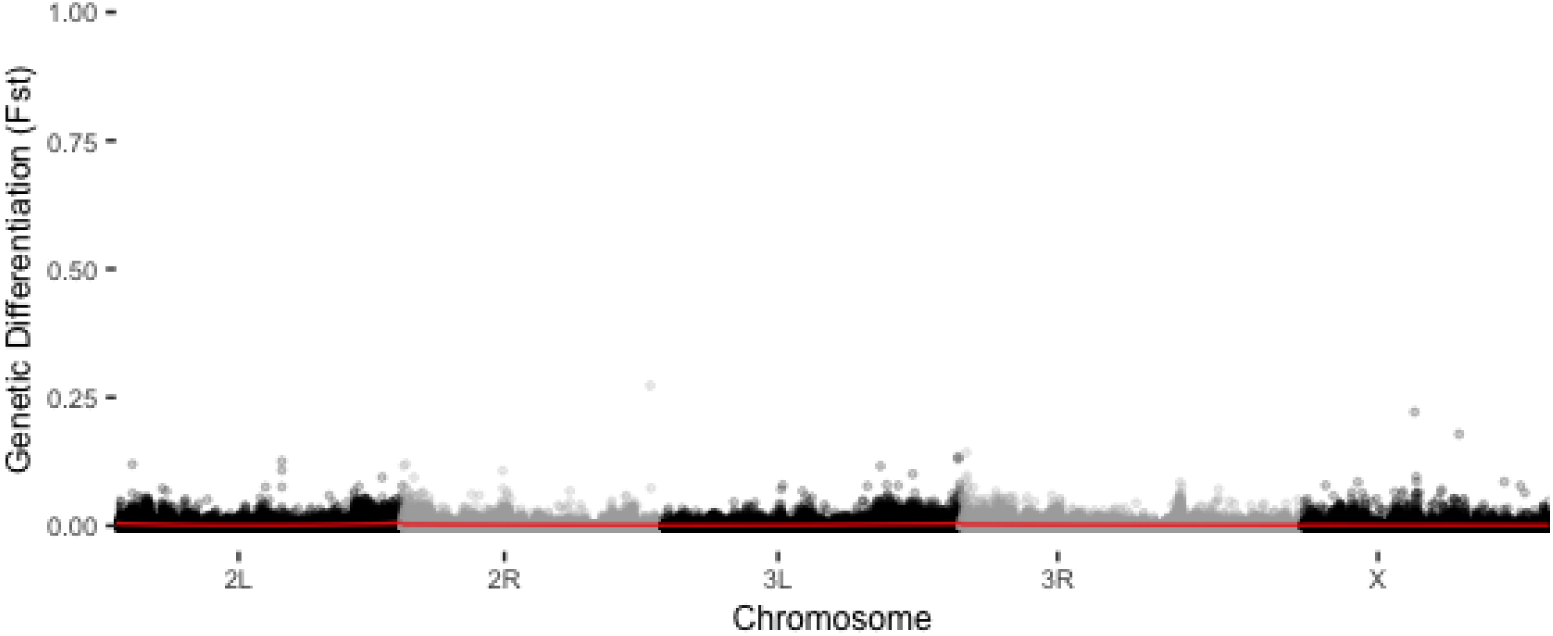
control replicate 2 males vs. females 1000bp

**Figure S13:**
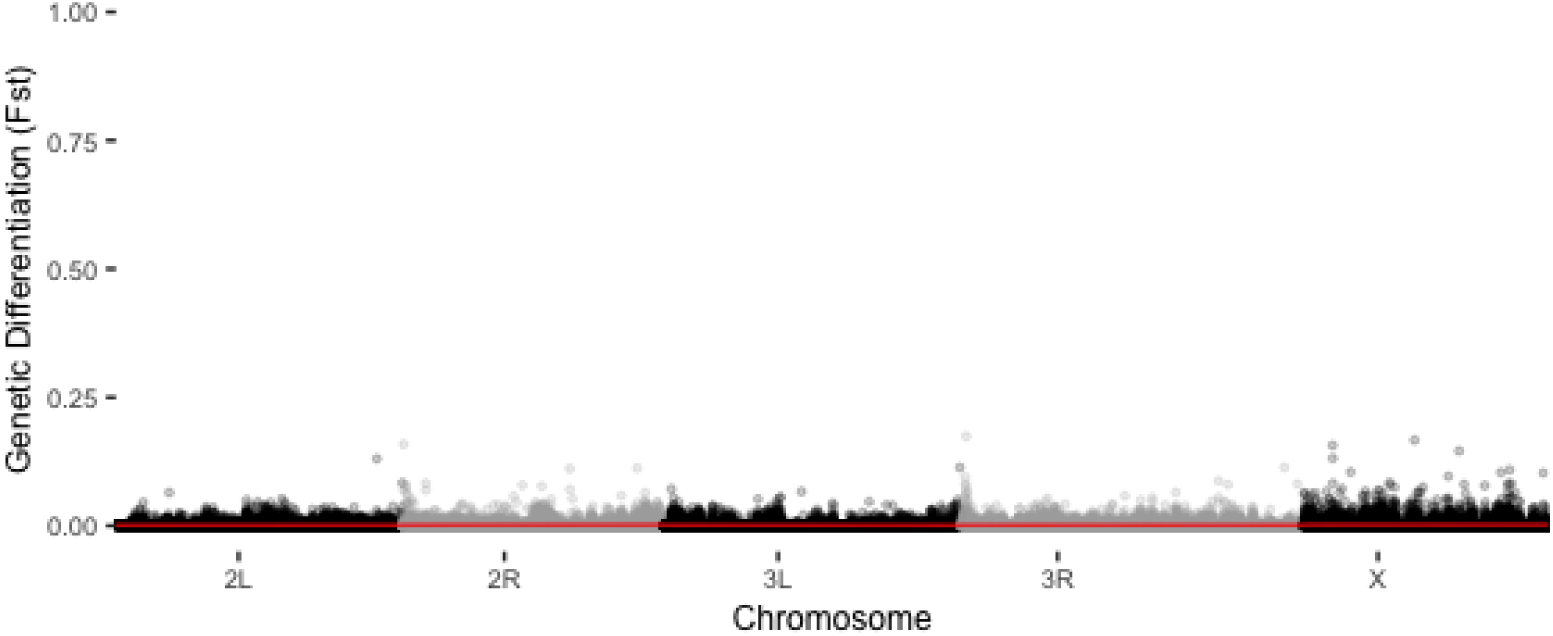
large replicate 1 males vs. females 1000bp

**Figure S14:**
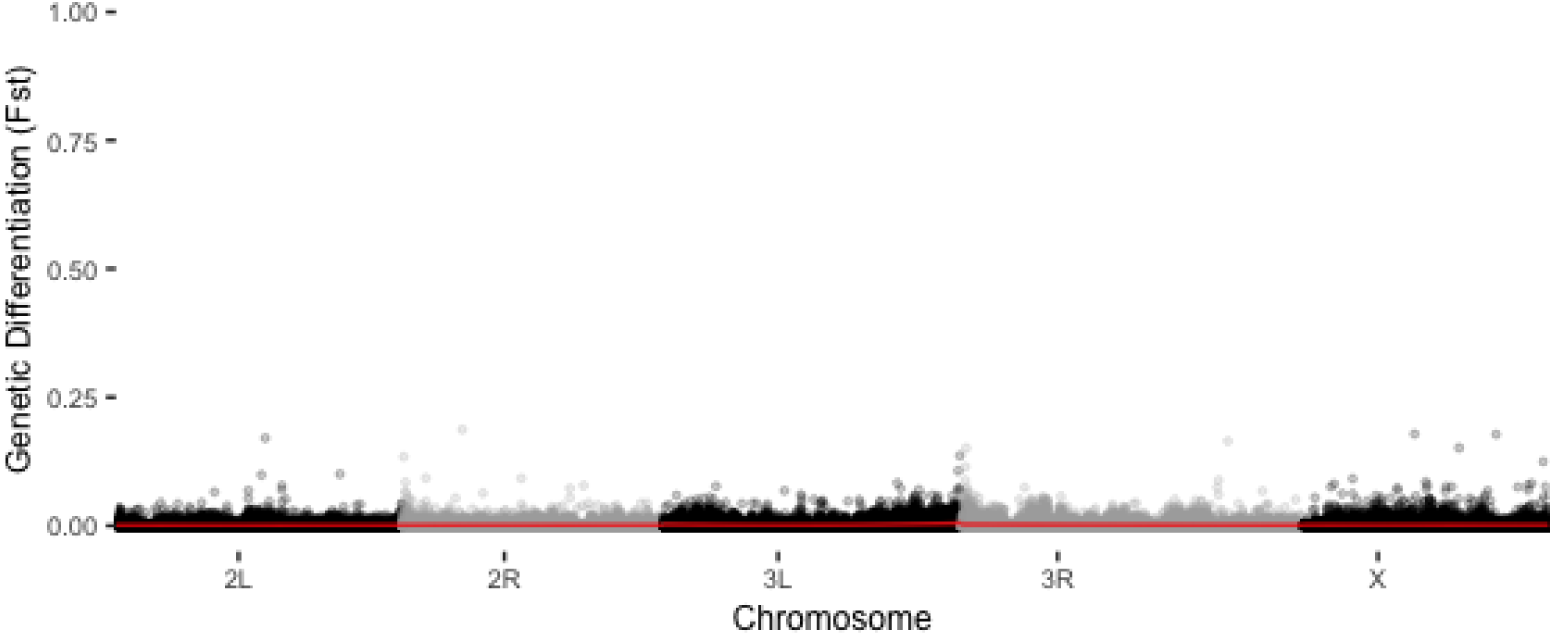
large replicate 2 males vs. females 1000bp

**Figure S15:**
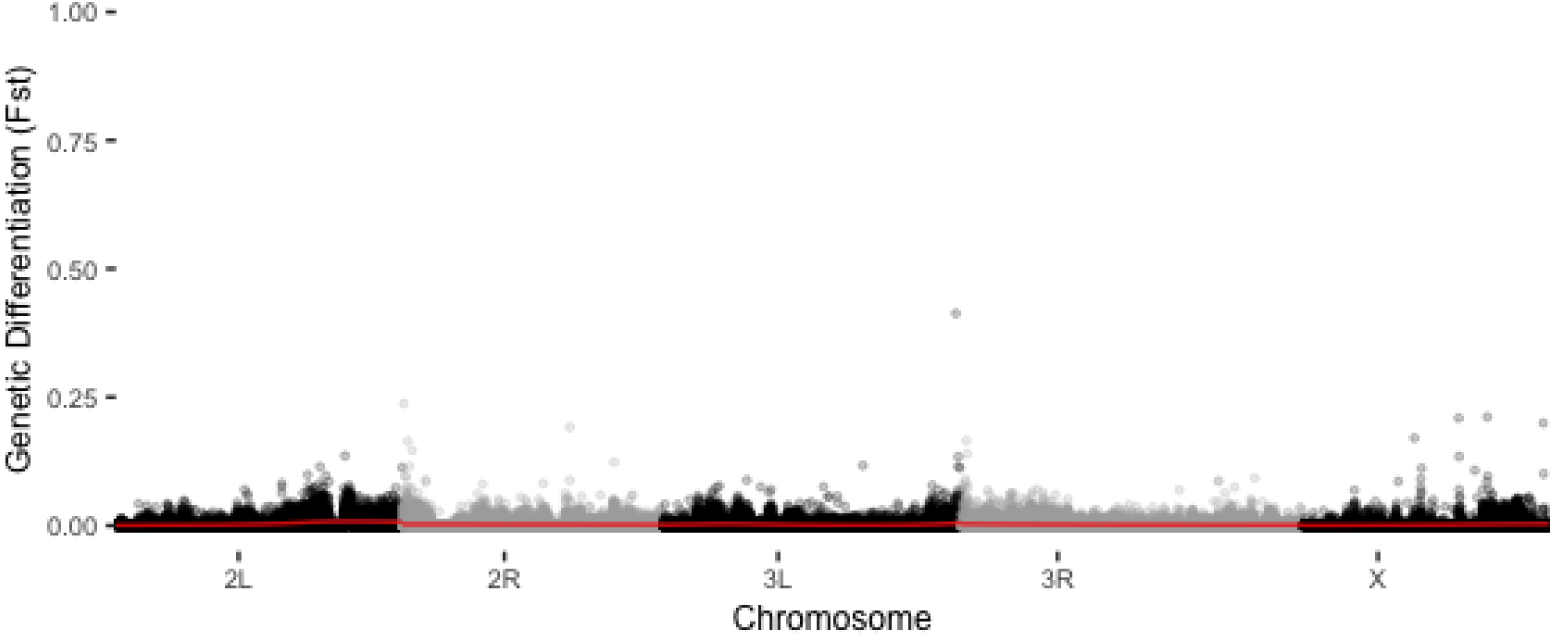
small replicate 1 males vs. females 1000bp

**Figure S16:**
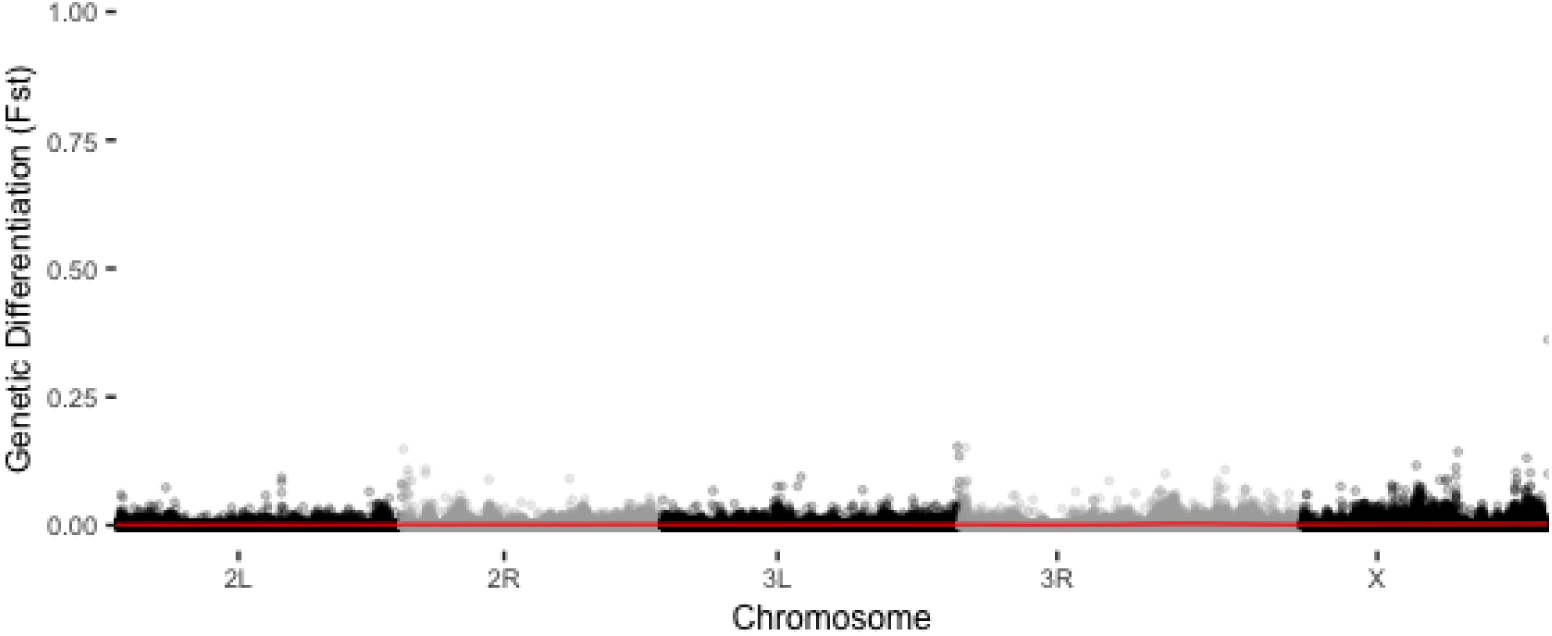
small replicate 2 males vs. females 1000bp

**Figure S17:**
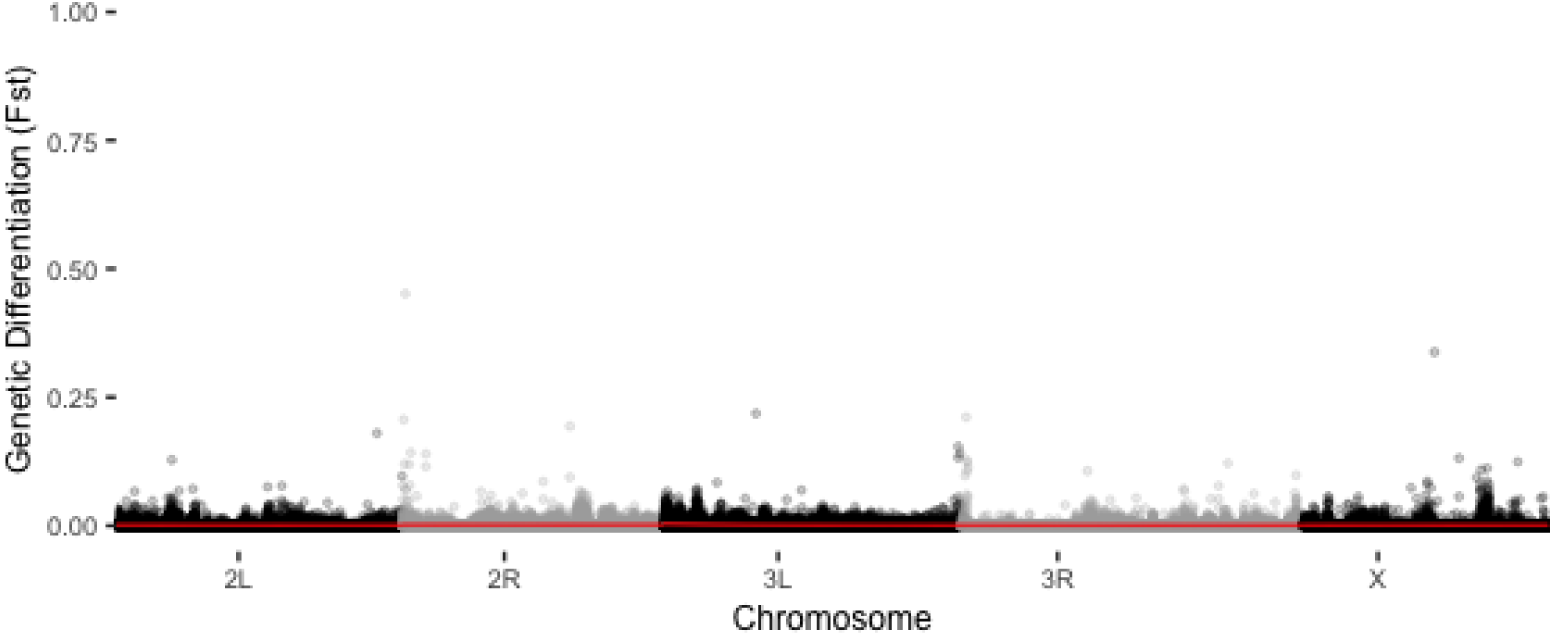
discordant replicate 2 males vs. females 1000bp

**Figure S18:**
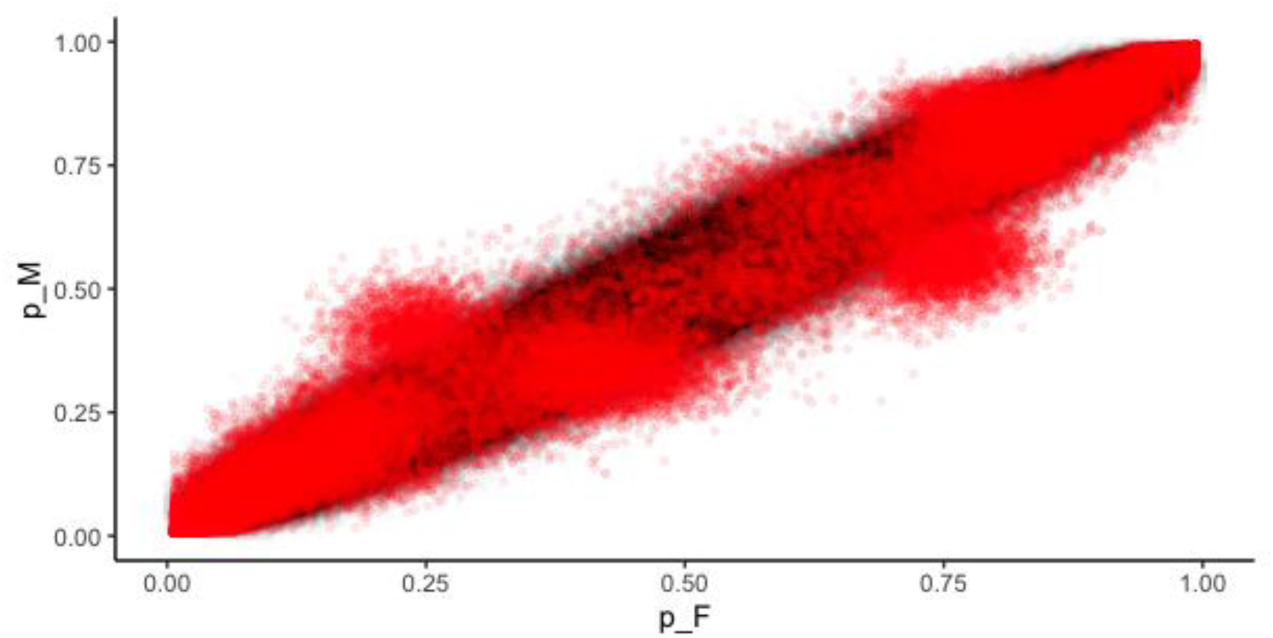
E1 male versus female minor allele frequency (computed as ALT/(REF+ALT)) in red plotted on top of 1000000 simulated variants with similar coverage and allele frequency distributions of our data.

**Figure S19:**
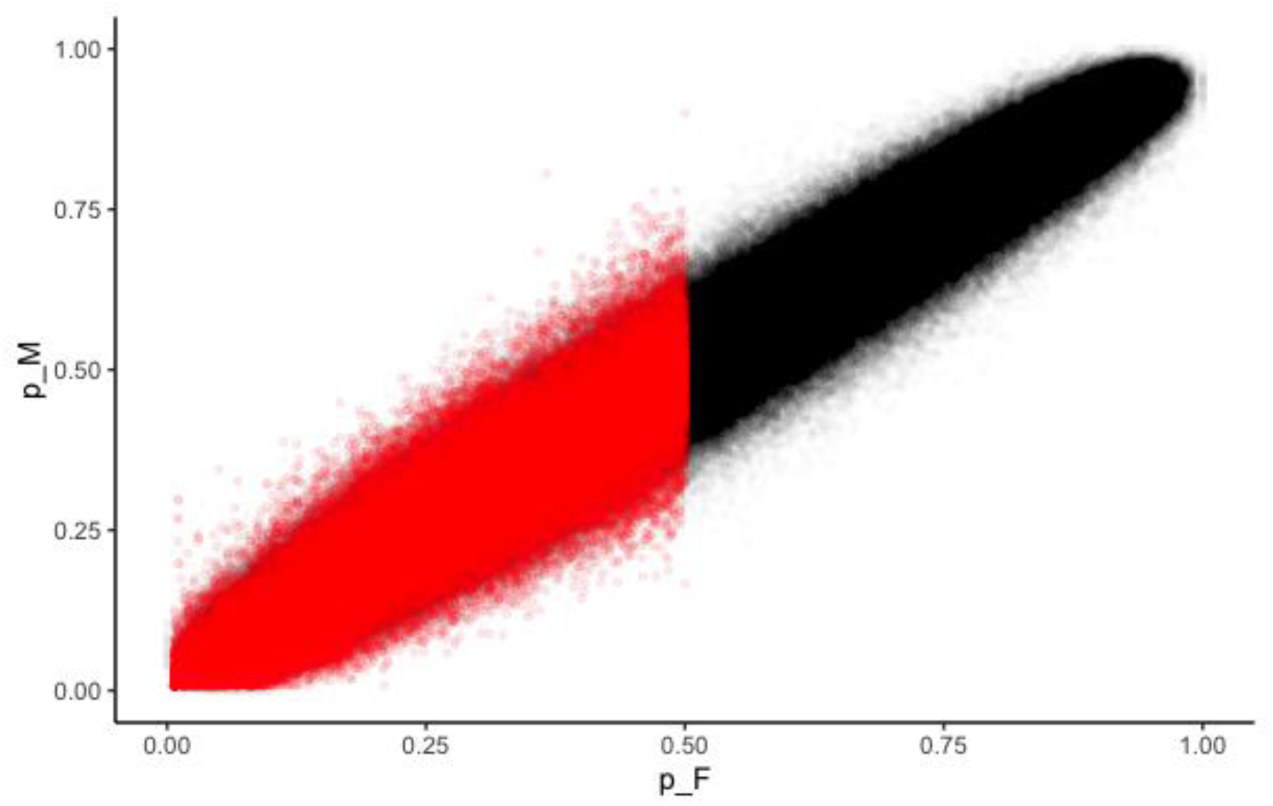
C1 male versus female minor allele frequency (computed as ALT/(REF+ALT)) in red plotted on top of 1000000 samples binomially sampled from a simulated pool with a similar coverage and distribution of our data. ***Please note***, the reference allele frequency for all treatments are set to the major allele of C1 females (this sample) so no alternate alleles are expected to reach >0.5 frequency.

**Figure S20:**
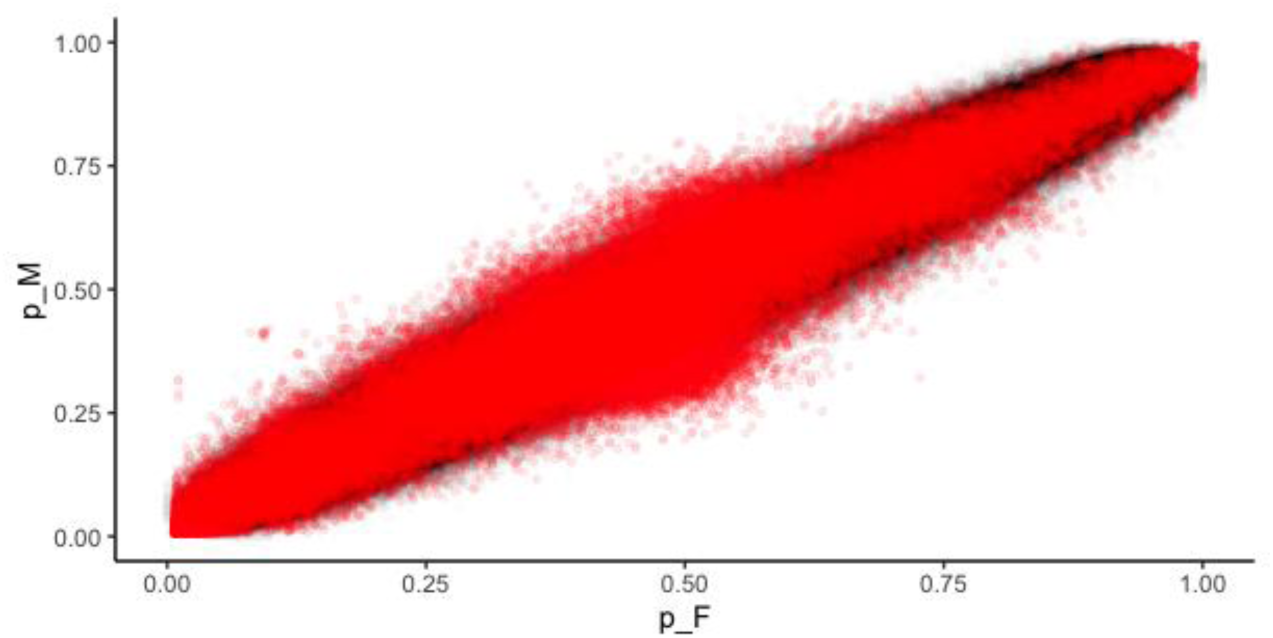
C2 male versus female minor allele frequency (computed as ALT/(REF+ALT)) in red plotted on top of 1000000 samples binomially sampled from a simulated pool with a similar coverage and distribution of our data.

**Figure S21:**
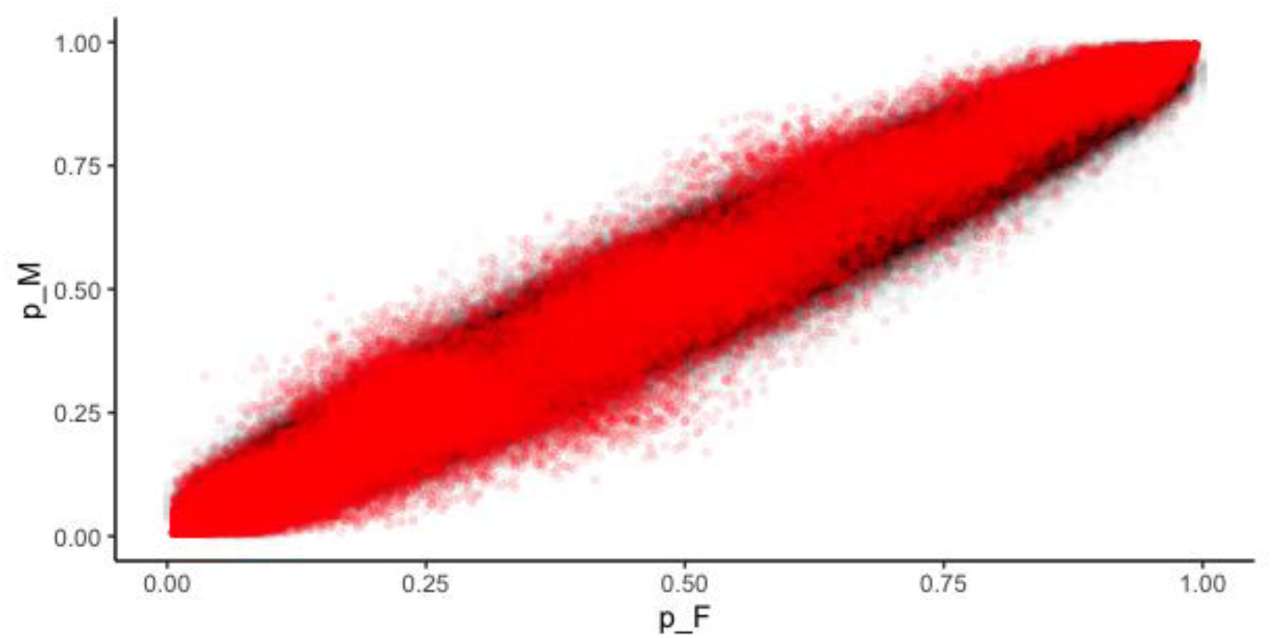
E2 male versus female minor allele frequency (computed as ALT/(REF+ALT)) in red plotted on top of 1000000 samples binomially sampled from a simulated pool with a similar coverage and distribution of our data.

**Figure S22:**
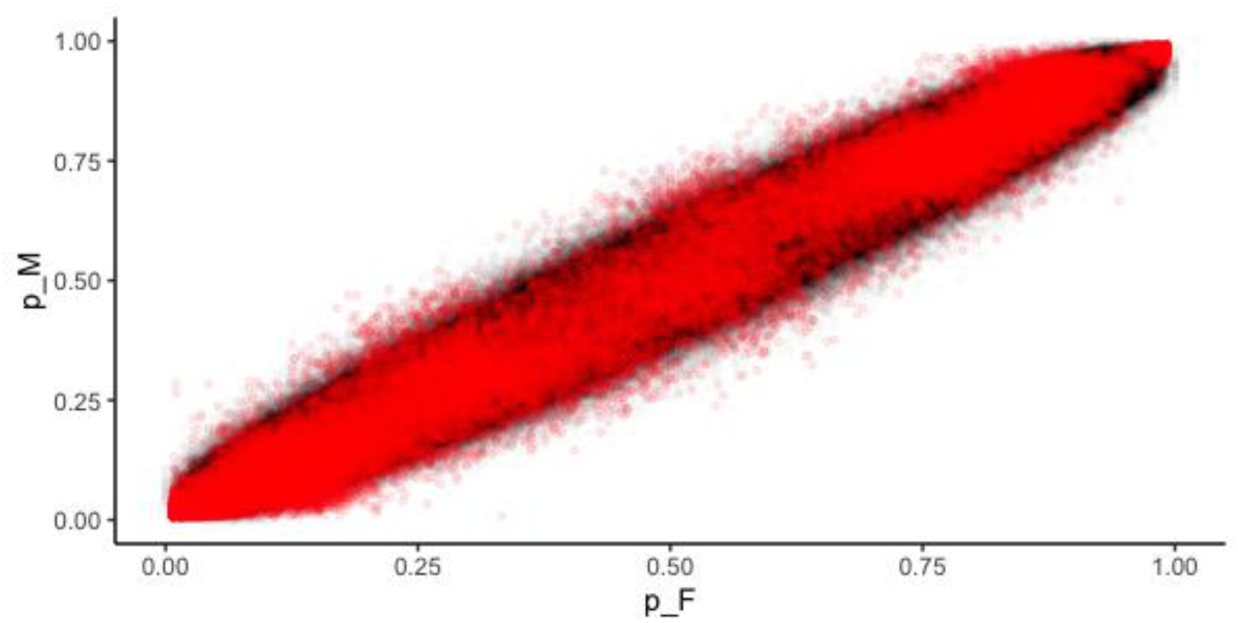
L1 male versus female minor allele frequency (computed as ALT/(REF+ALT)) in red plotted on top of 1000000 samples binomially sampled from a simulated pool with a similar coverage and distribution of our data.

**Figure S23:**
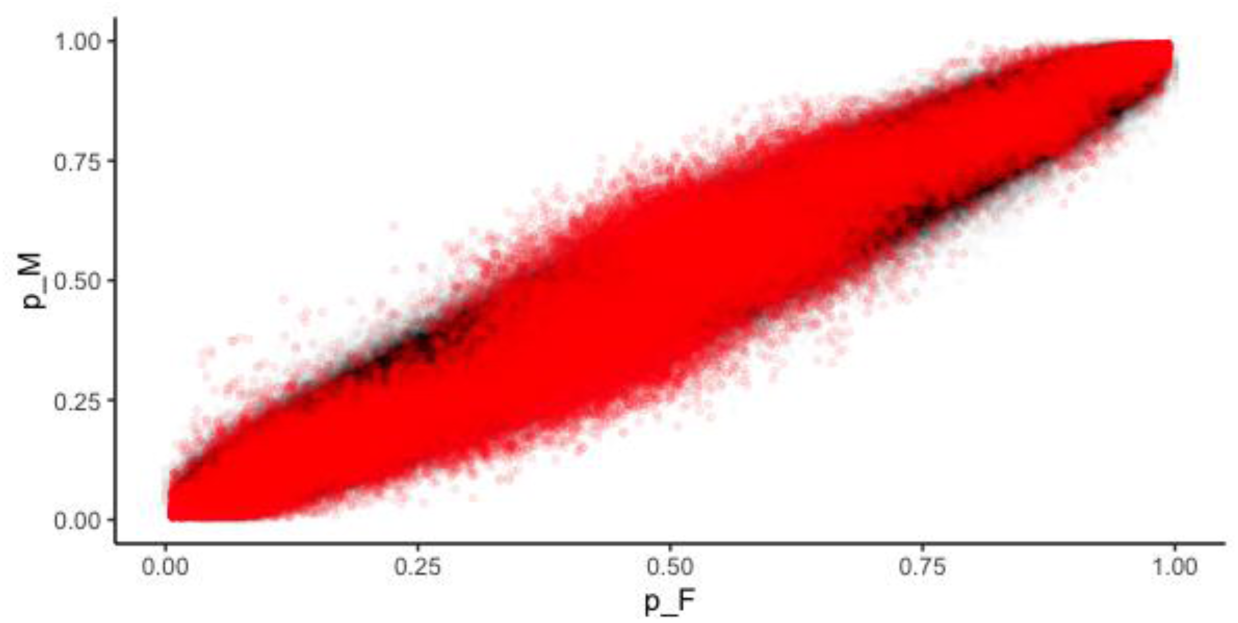
L2 male versus female minor allele frequency (computed as ALT/(REF+ALT)) in red plotted on top of 1000000 samples binomially sampled from a simulated pool with a similar coverage and distribution of our data.

**Figure S24:**
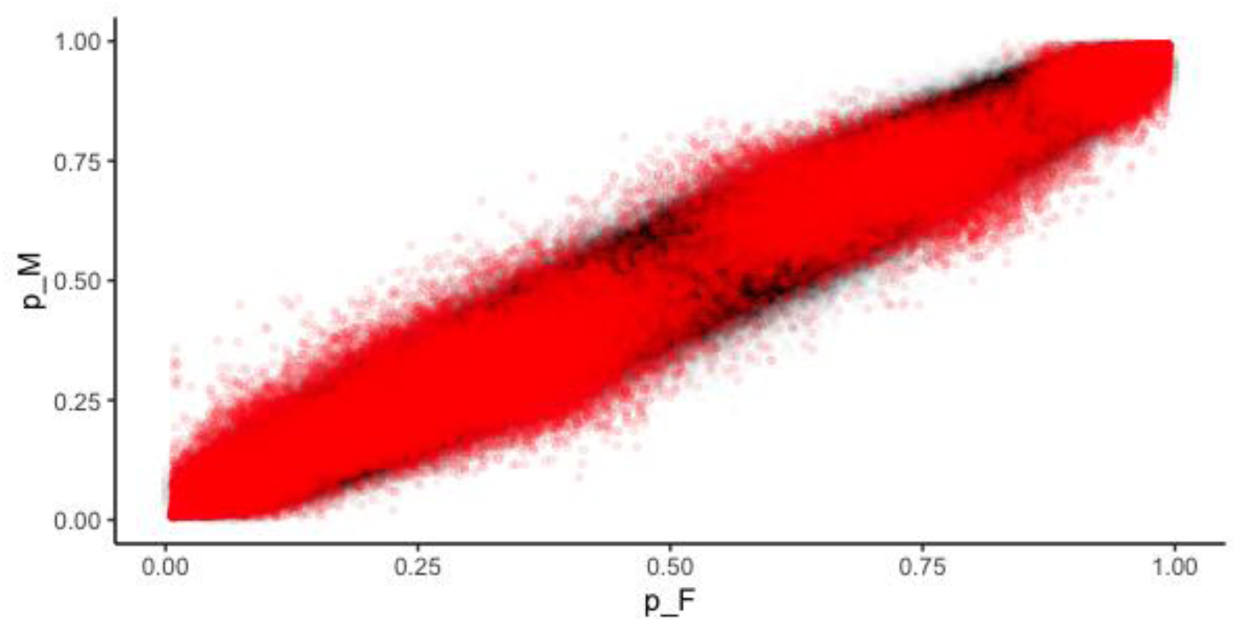
S1 male versus female minor allele frequency (computed as ALT/(REF+ALT)) in red plotted on top of 1000000 samples binomially sampled from a simulated pool with a similar coverage and distribution of our data.

**Figure S25:**
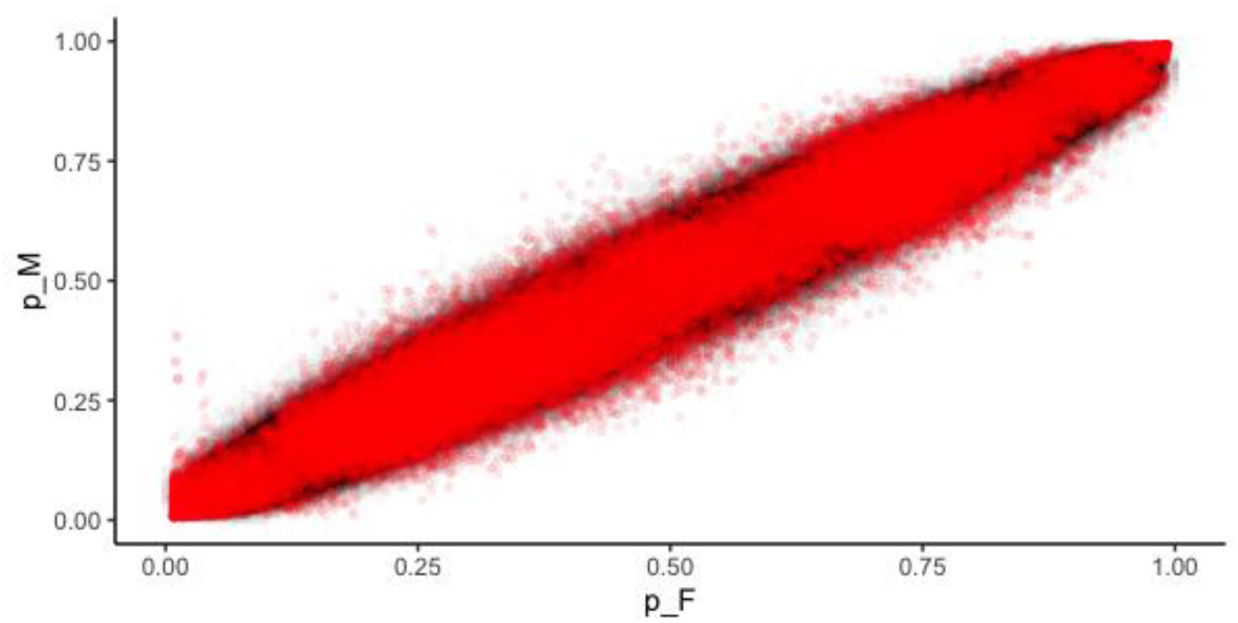
S2 male versus female minor allele frequency (computed as ALT/(REF+ALT)) in red plotted on top of 1000000 samples binomially sampled from a simulated pool with a similar coverage and distribution of our data.

**Figure S26:**
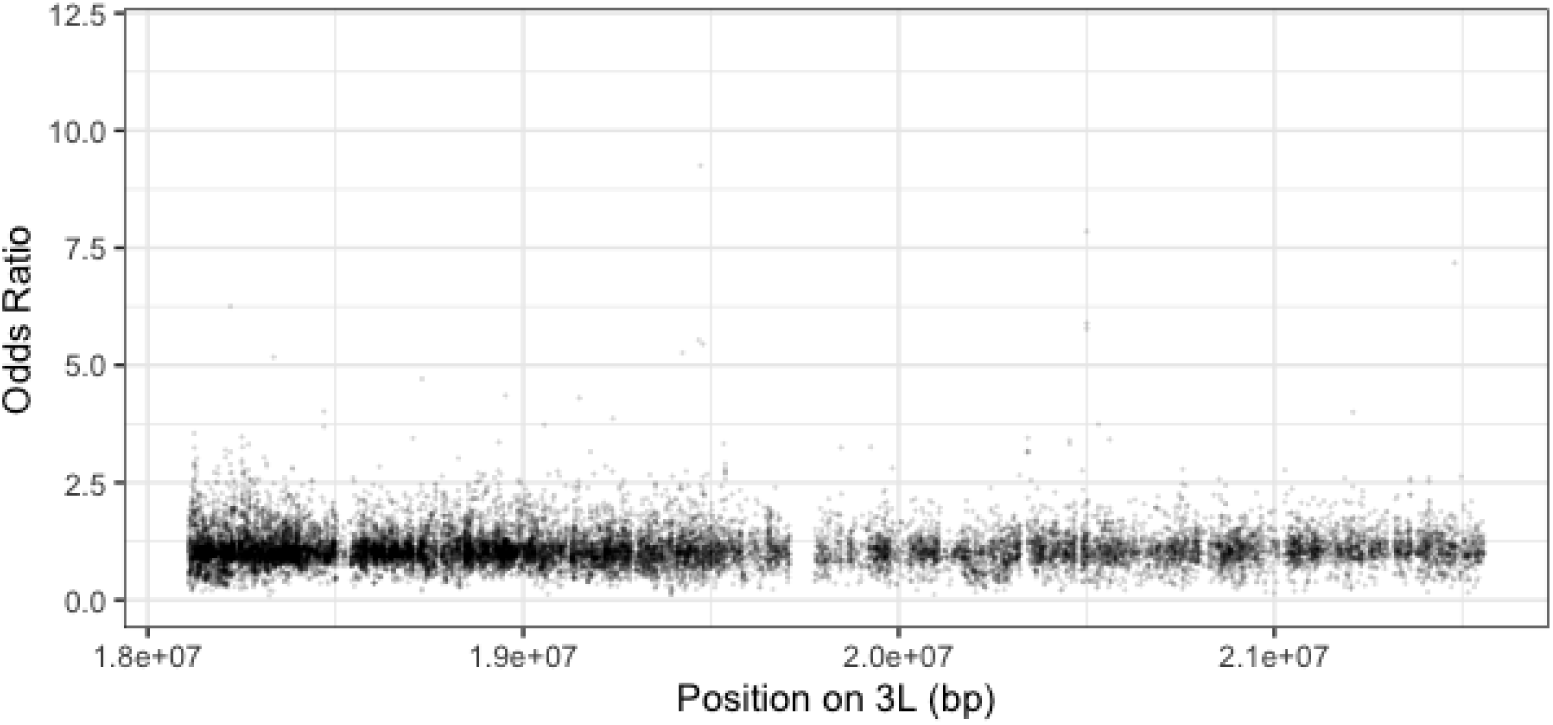
C1 modelled odds ratios by position on chromosome 3L within the elevated region on E1. Positions with a p-value < 0.0001 highlighted red with upper and lower 95% confidence intervals shown.

**Figure S27:**
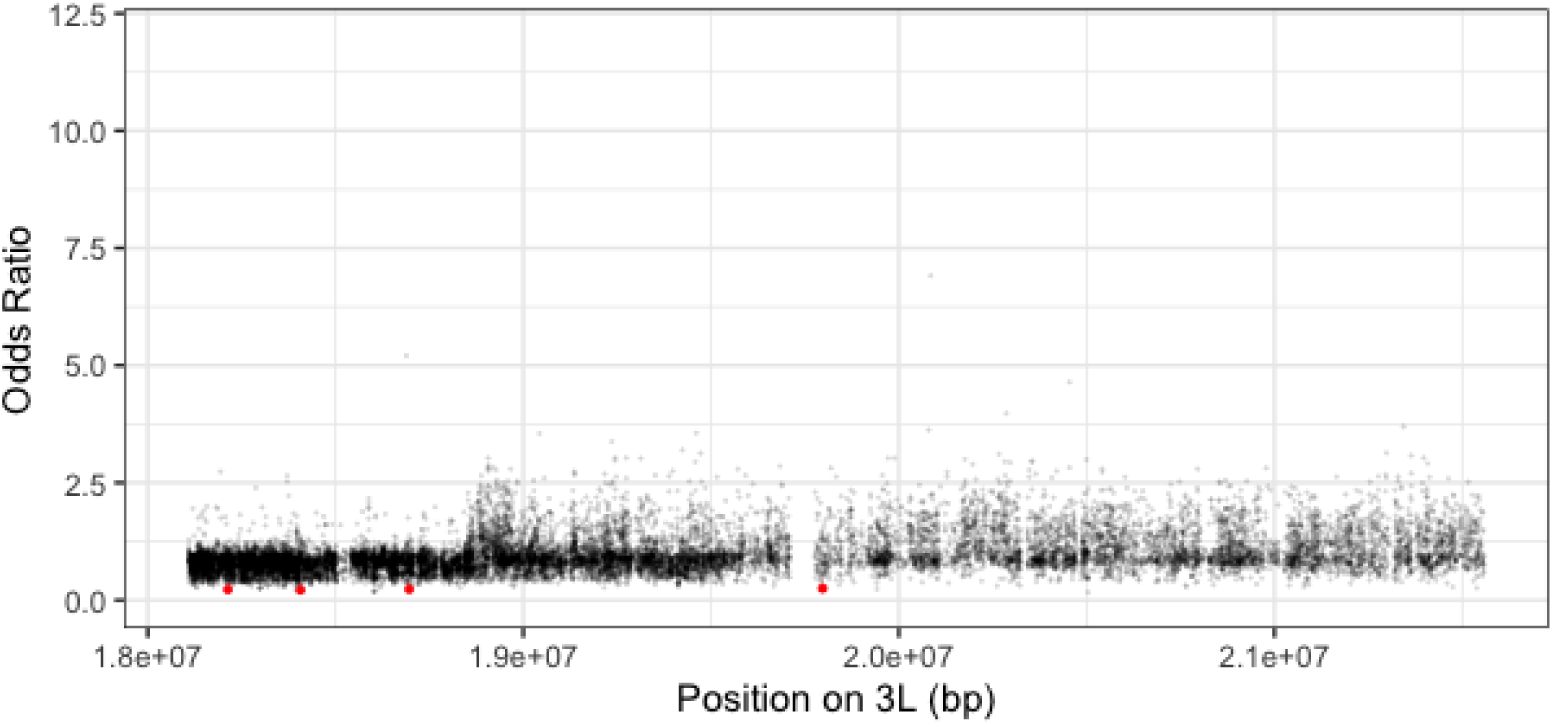
C2 modelled odds ratios by position on chromosome 3L within the elevated region on E1. Positions with a p-value < 0.0001 highlighted red with upper and lower 95% confidence intervals shown.

**Figure S28:**
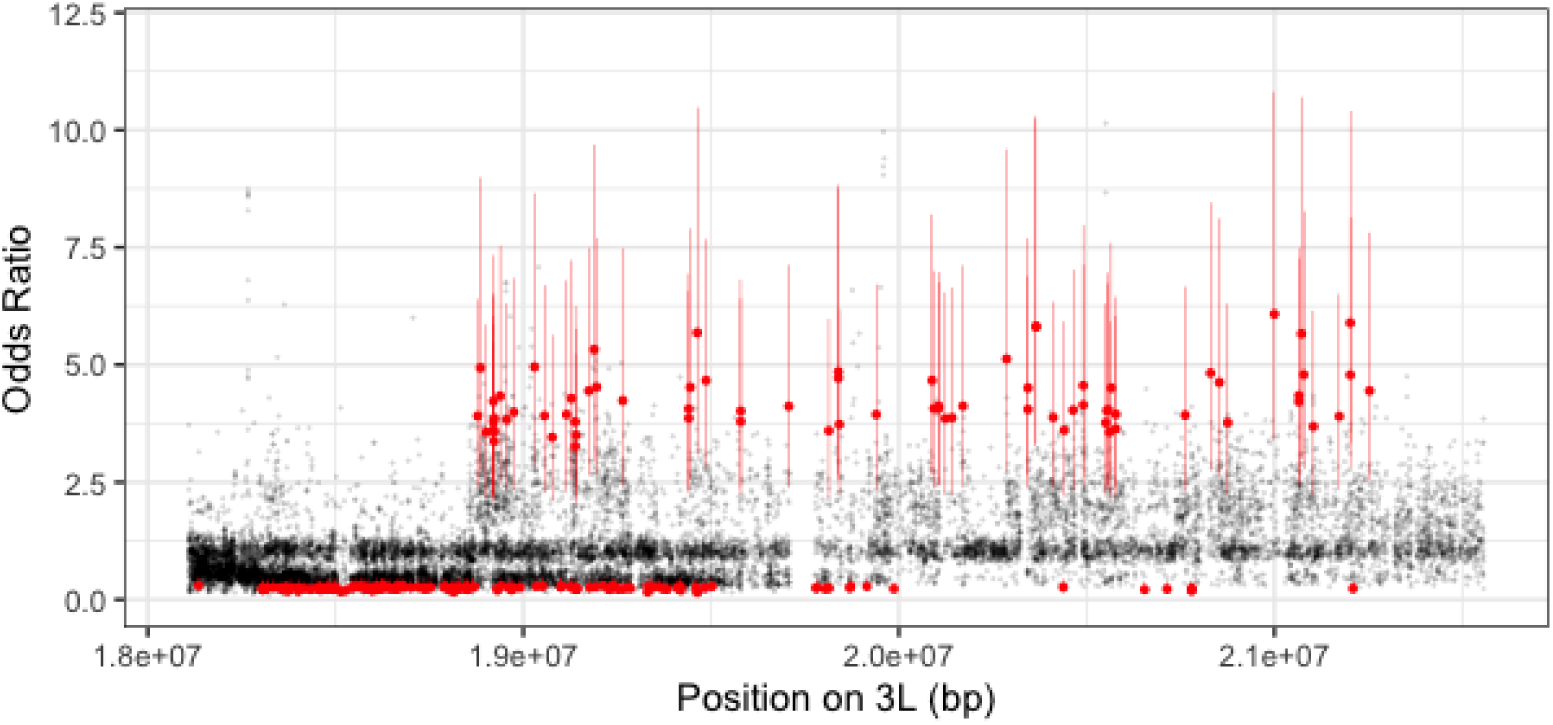
E1 modelled odds ratios by position on chromosome 3L within the elevated region on E1. Positions with a p-value < 0.0001 highlighted red with upper and lower 95% confidence intervals shown.

**Figure S29:**
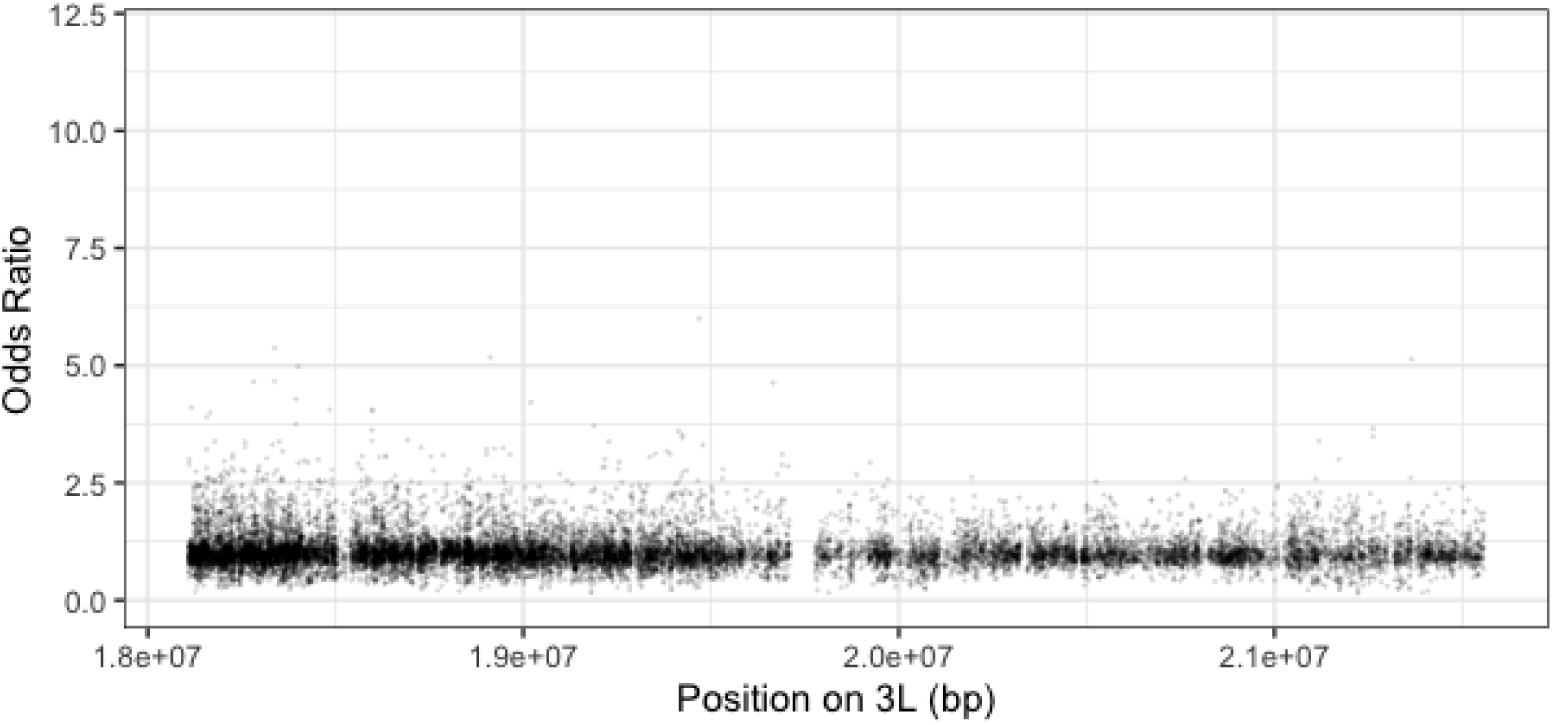
E2 modelled odds ratios by position on chromosome 3L within the elevated region on E1. Positions with a p-value < 0.0001 highlighted red with upper and lower 95% confidence intervals shown.

**Figure S30:**
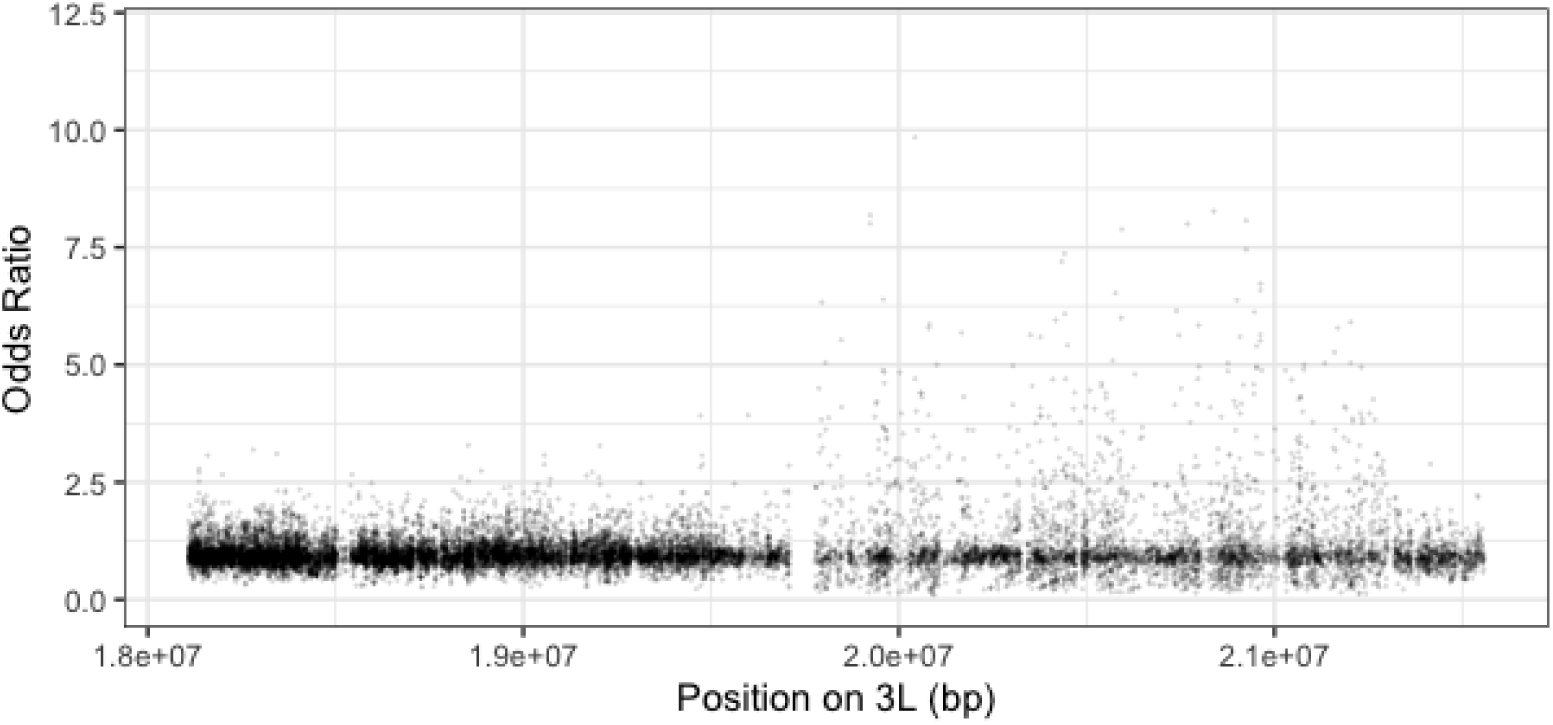
L1 modelled odds ratios by position on chromosome 3L within the elevated region on E1. Positions with a p-value < 0.0001 highlighted red with upper and lower 95% confidence intervals shown.

**Figure S31:**
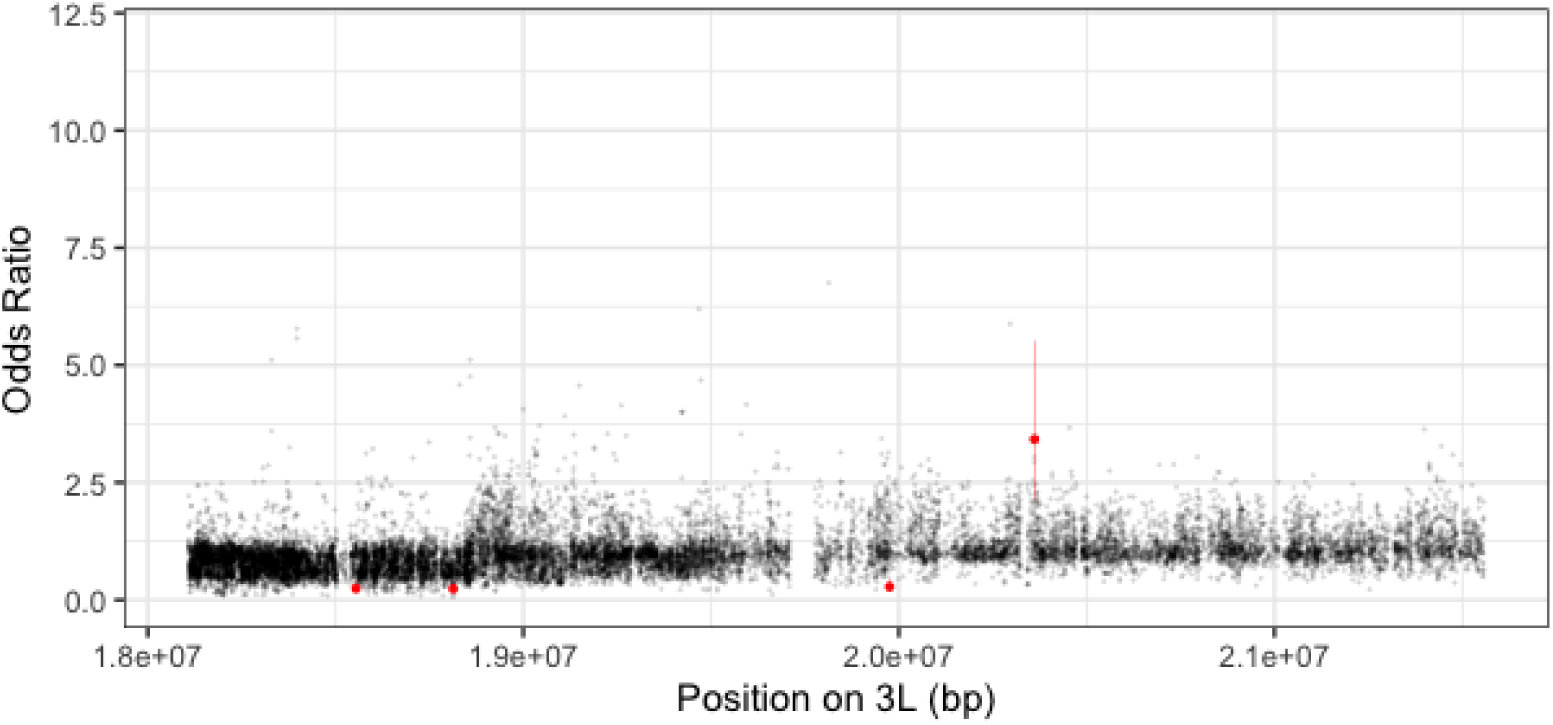
L2 modelled odds ratios by position on chromosome 3L within the elevated region on E1. Positions with a p-value < 0.0001 highlighted red with upper and lower 95% confidence intervals shown.

**Figure S32:**
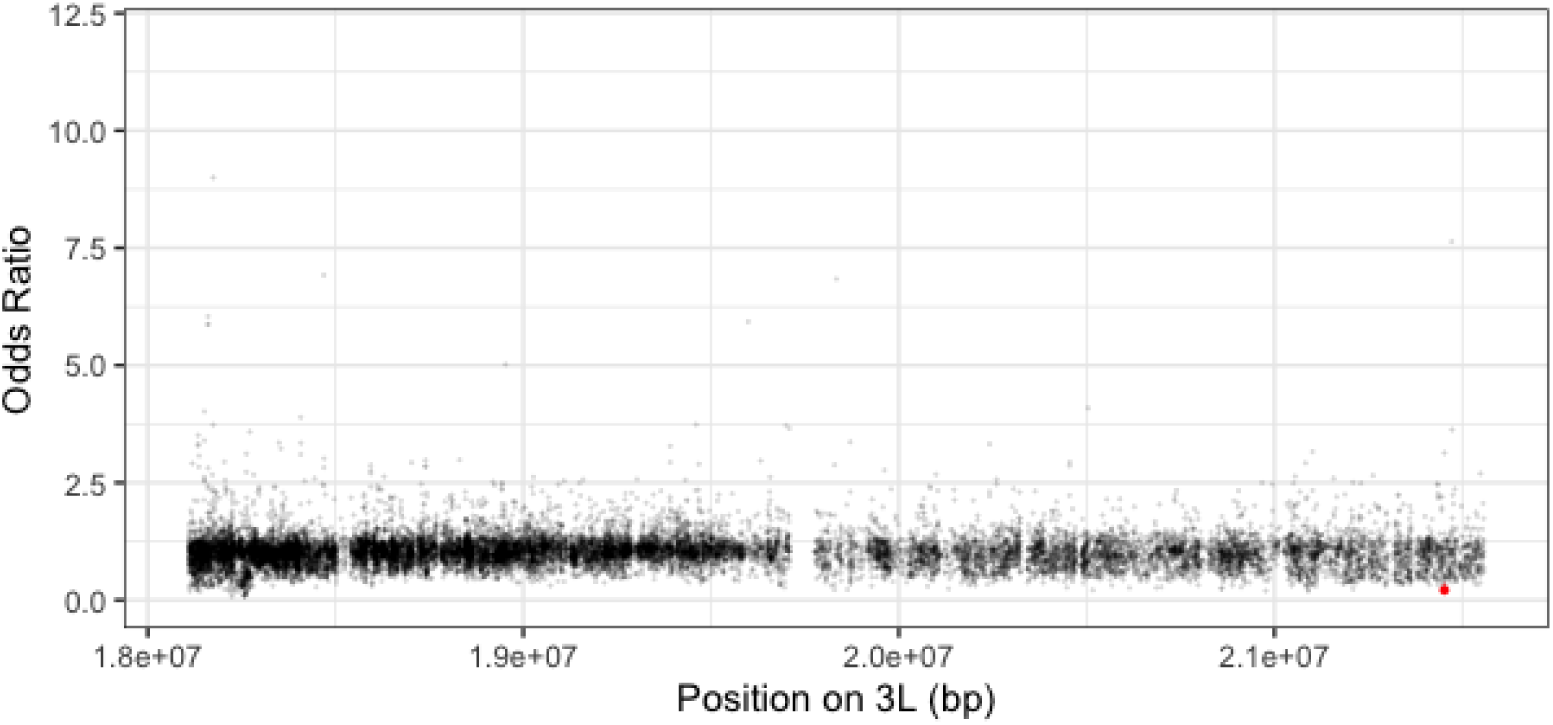
S1 modelled odds ratios by position on chromosome 3L within the elevated region on E1. Positions with a p-value < 0.0001 highlighted red with upper and lower 95% confidence intervals shown.

**Figure S33:**
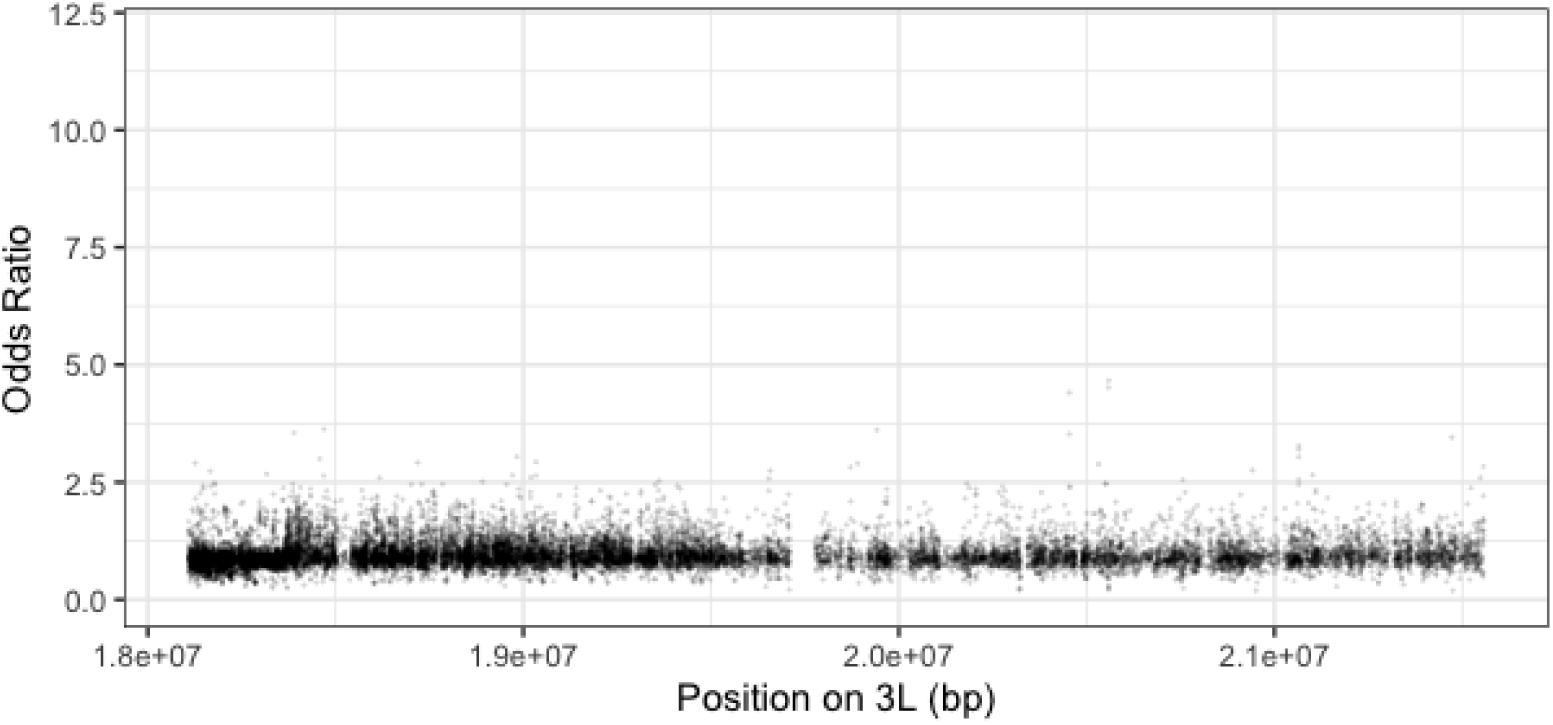
S2 modelled odds ratios by position on chromosome 3L within the elevated region on E1. Positions with a p-value < 0.0001 highlighted red with upper and lower 95% confidence intervals shown.

**Figure S34:**
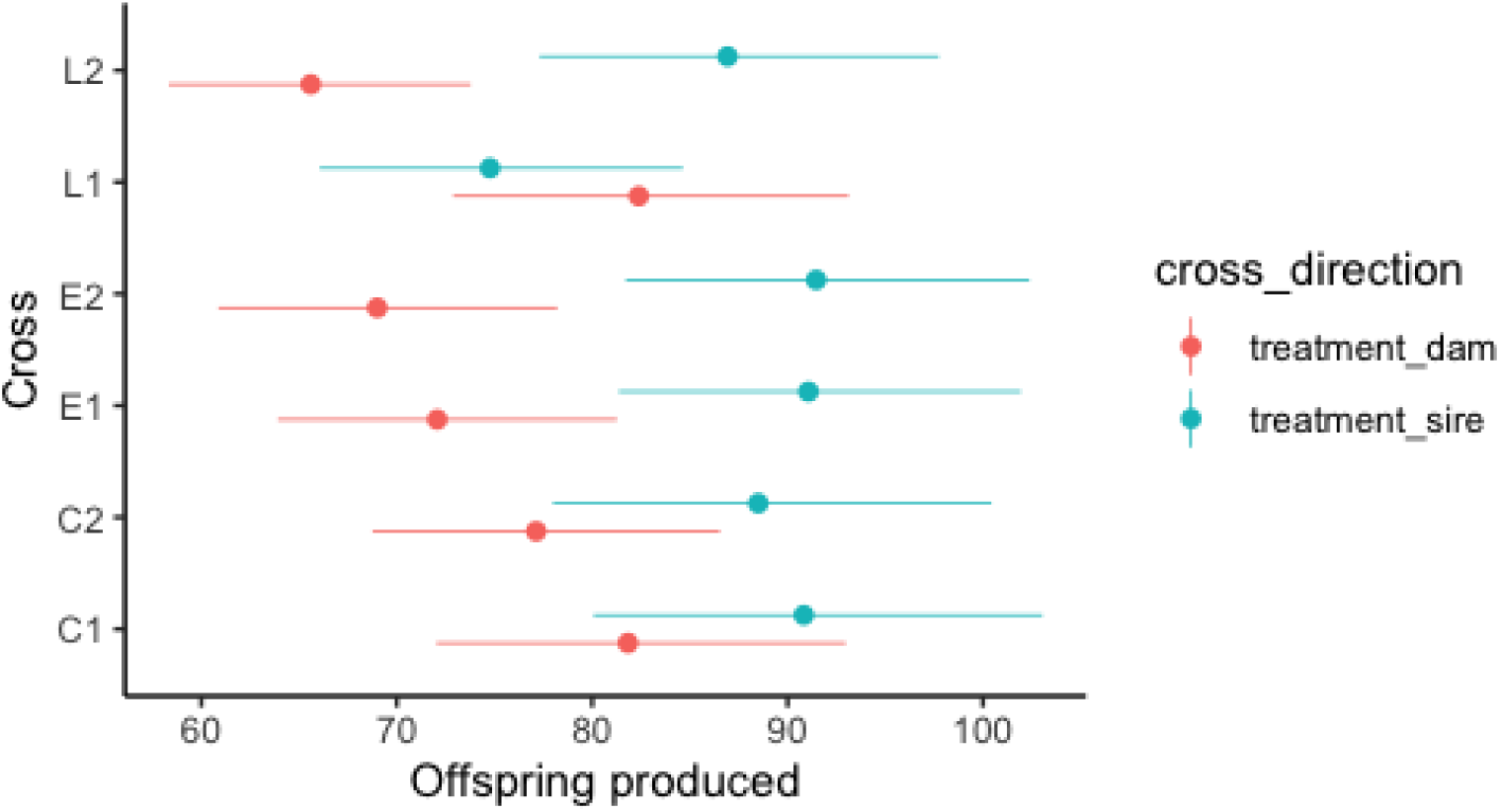
Fecundity measure for F1 of our sex ratio cross based on F0 treatment

**Figure S35:**
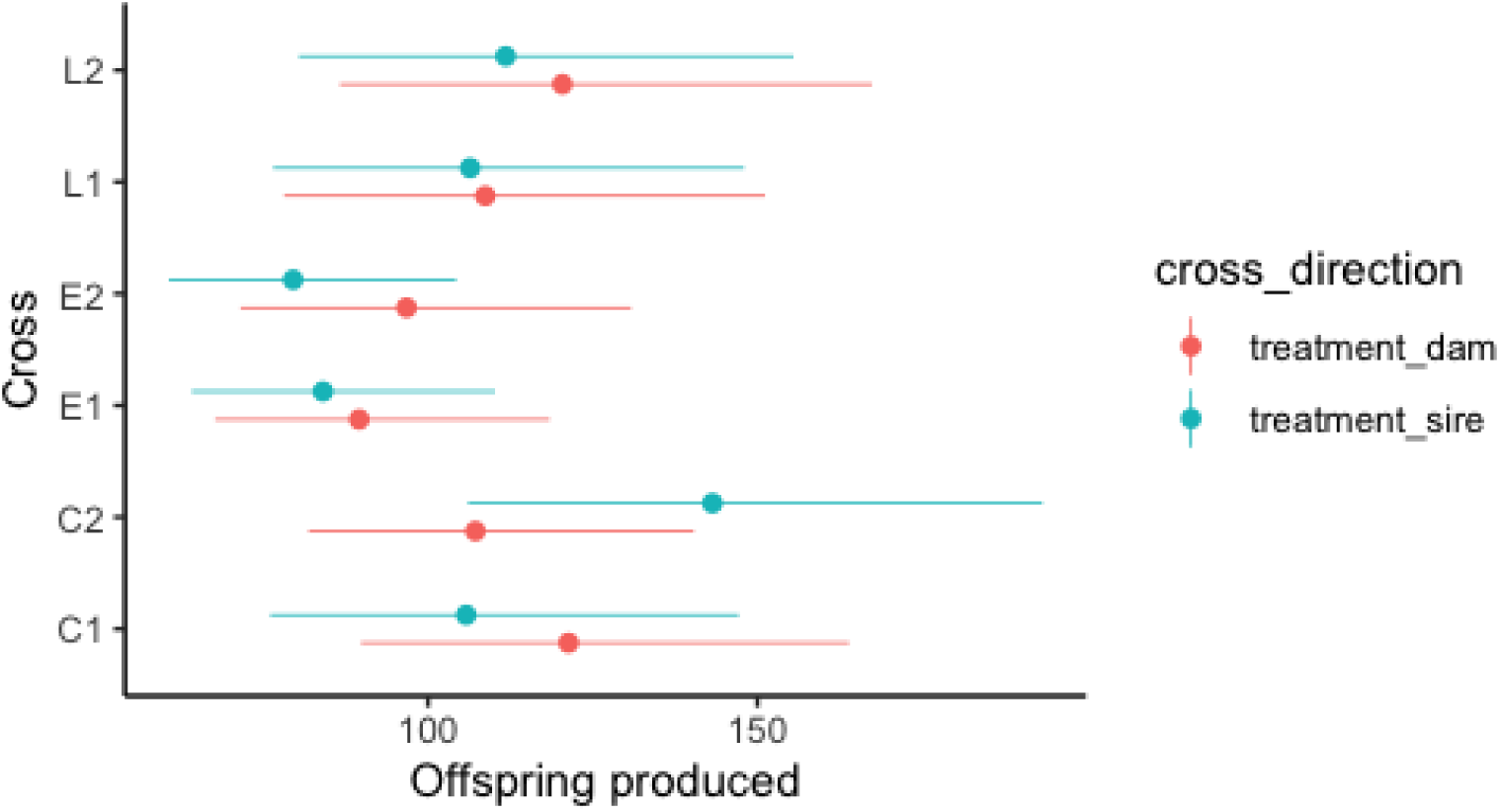
Fecundity measure for F2 of our sex ratio cross based on F0 treatment.

**Table S6:**
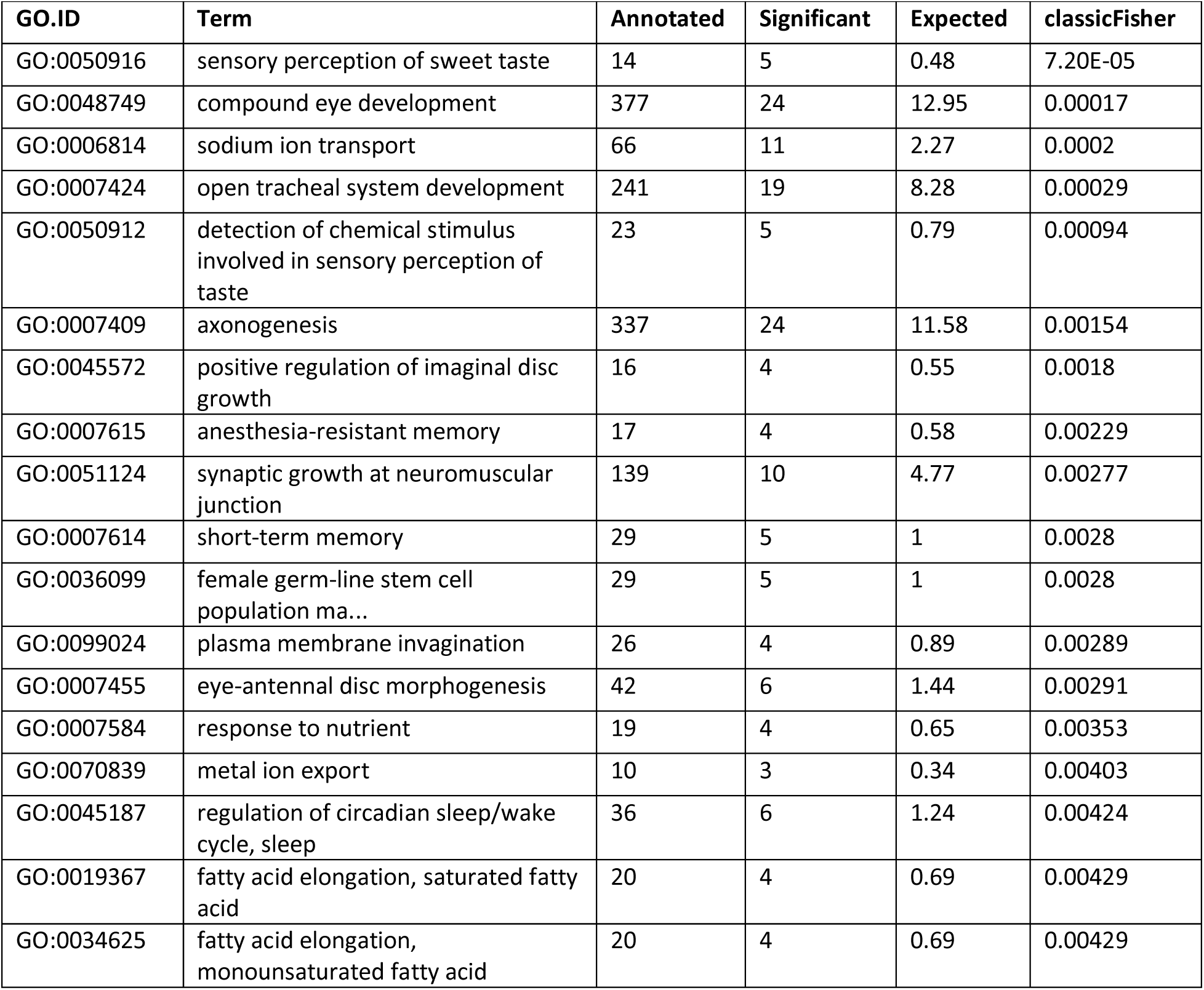
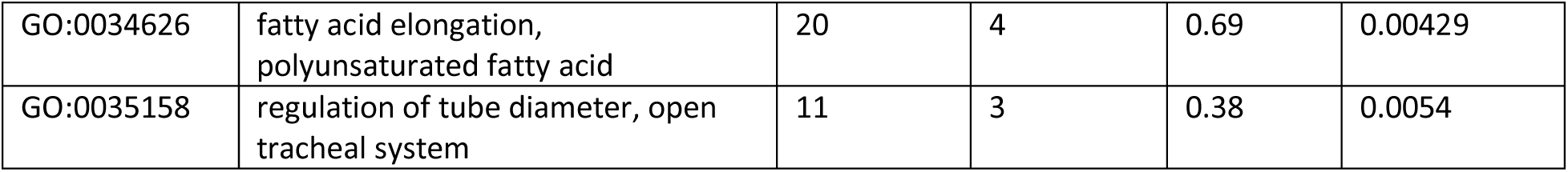
Discordant -Top 20 biological process GO terms.

**Table S7:**
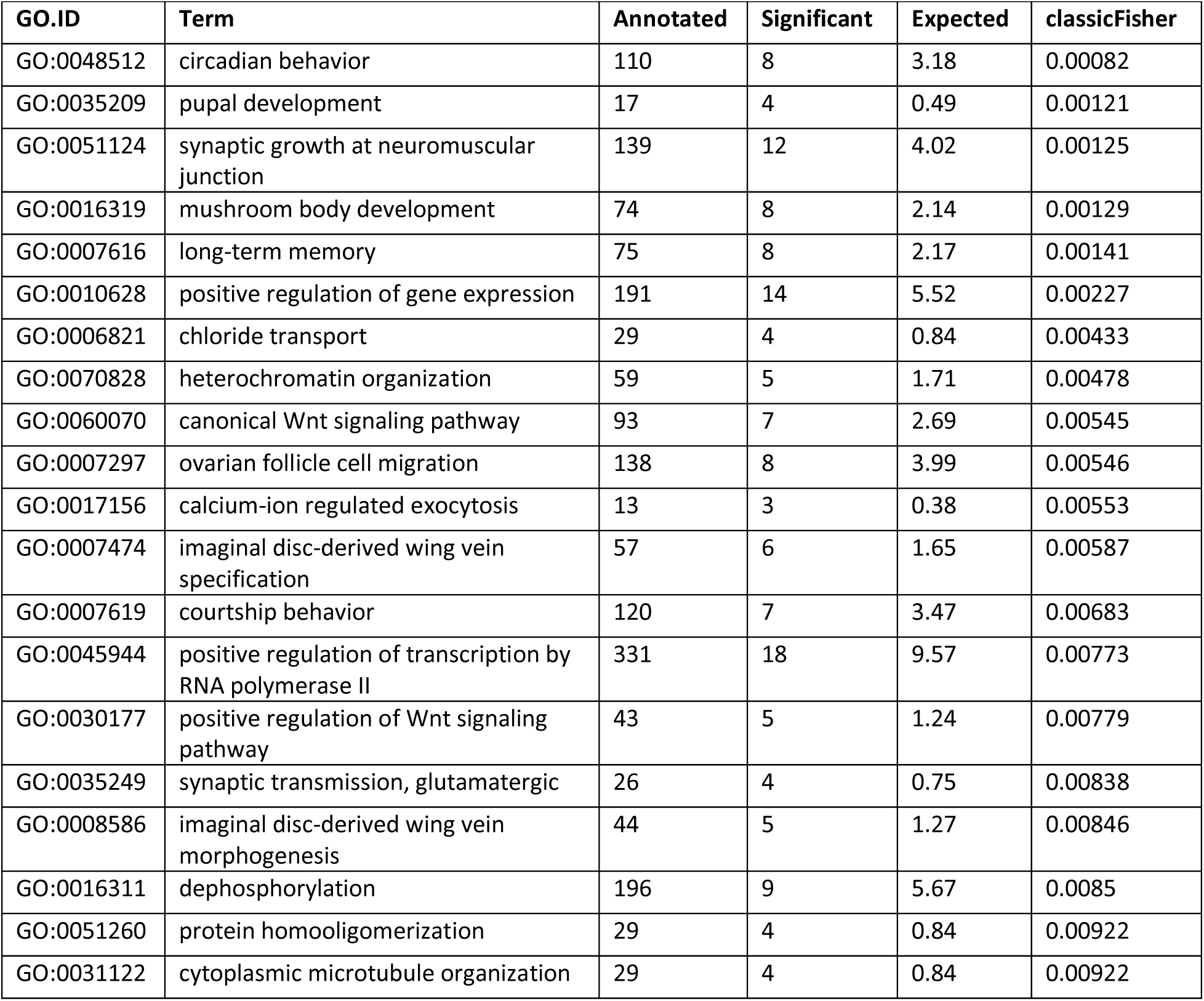
Large - Top 20 biological process GO terms.

**Table S8:**
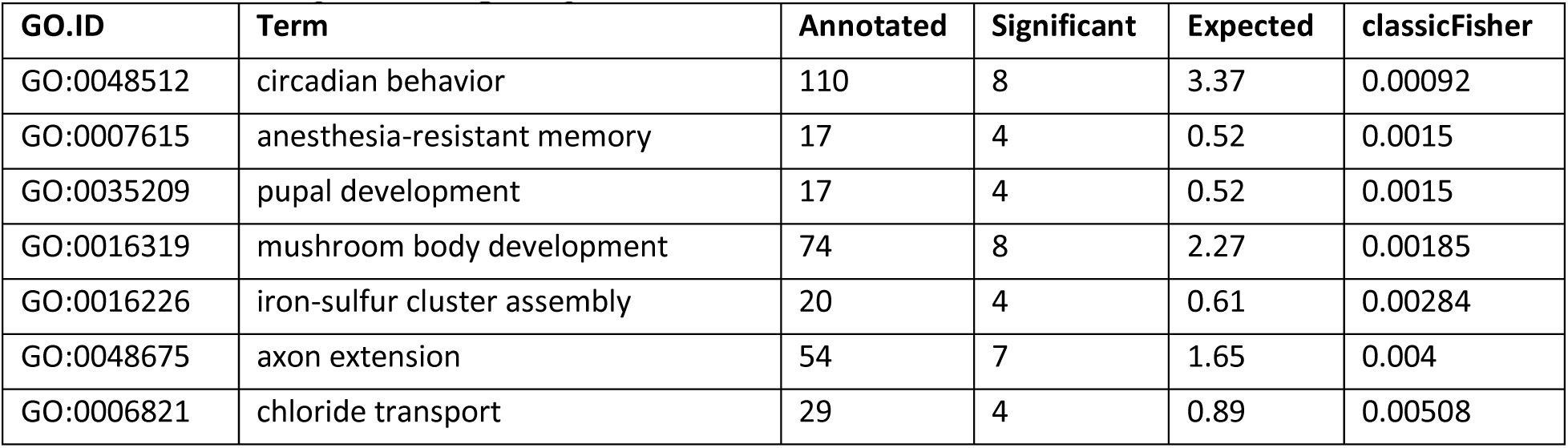
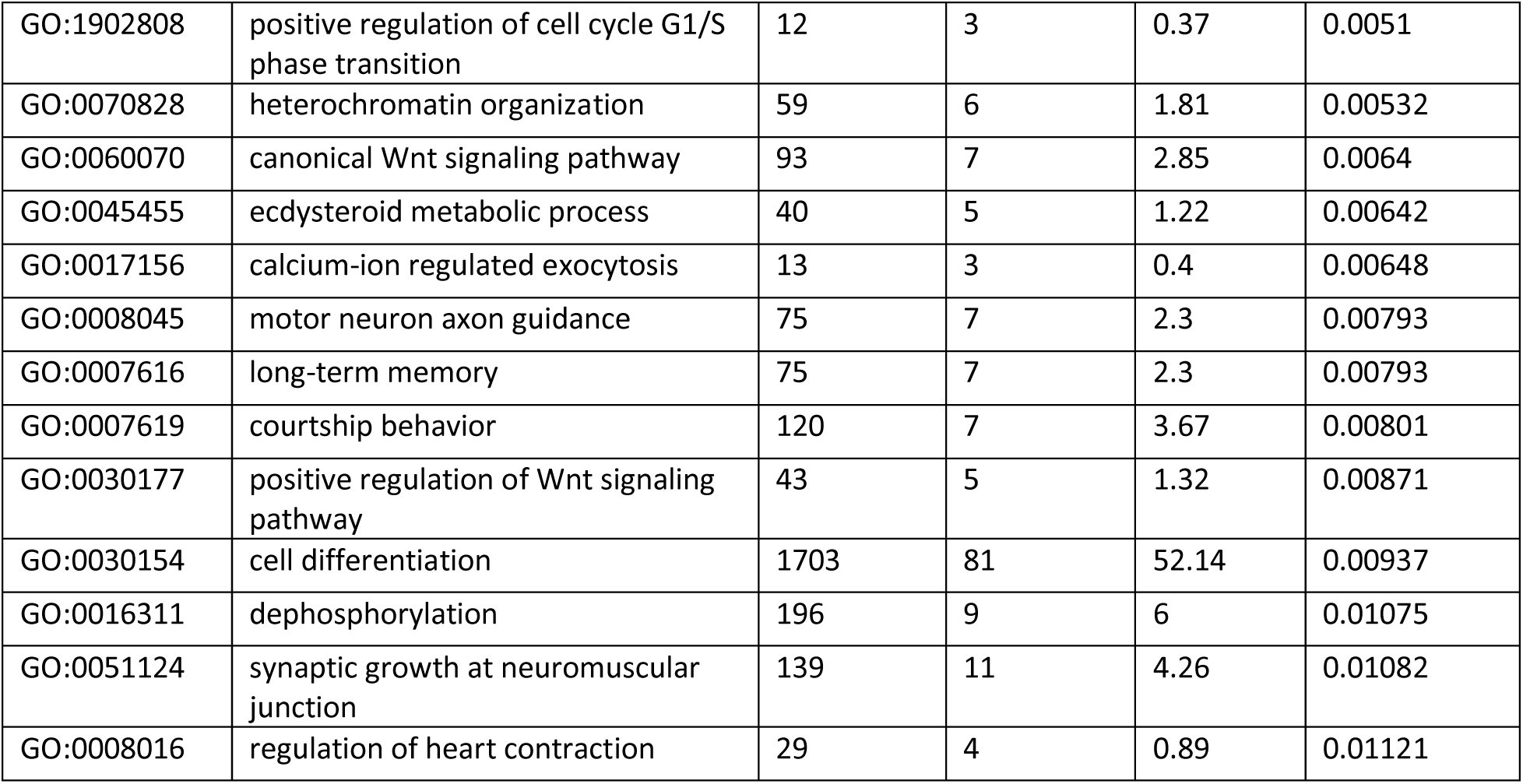
Small Top 20 biological process GO terms.

**Table S9:**
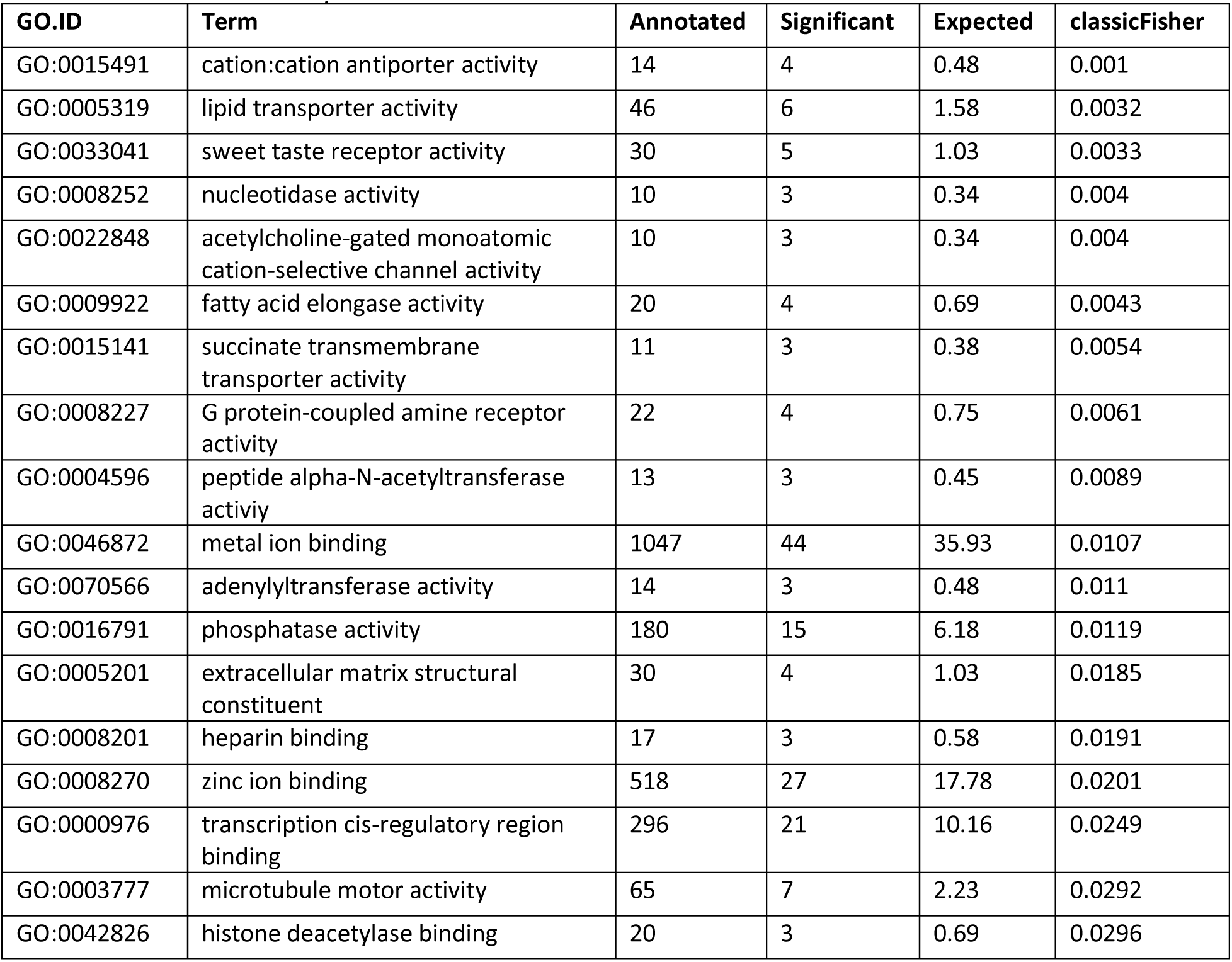
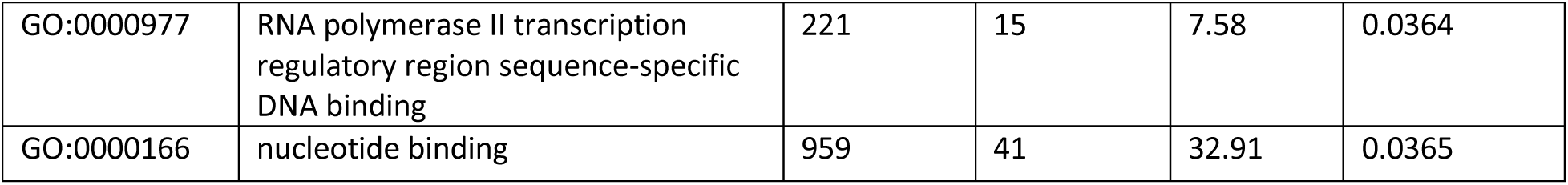
discordant - Top 20 molecular function GO terms.

**Table S10:**
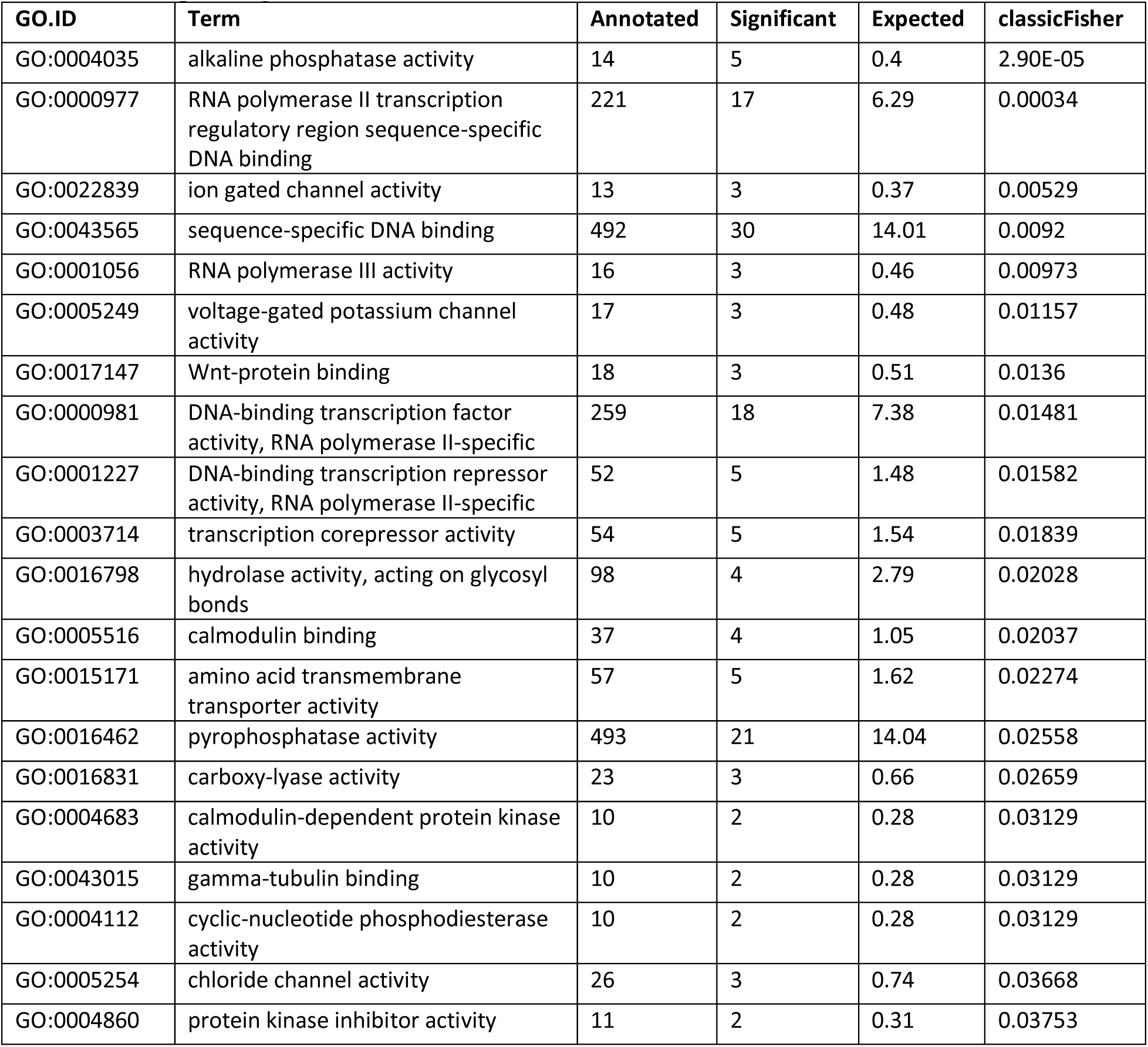
Large - Top 20 molecular function GO terms.

**Table S11:**
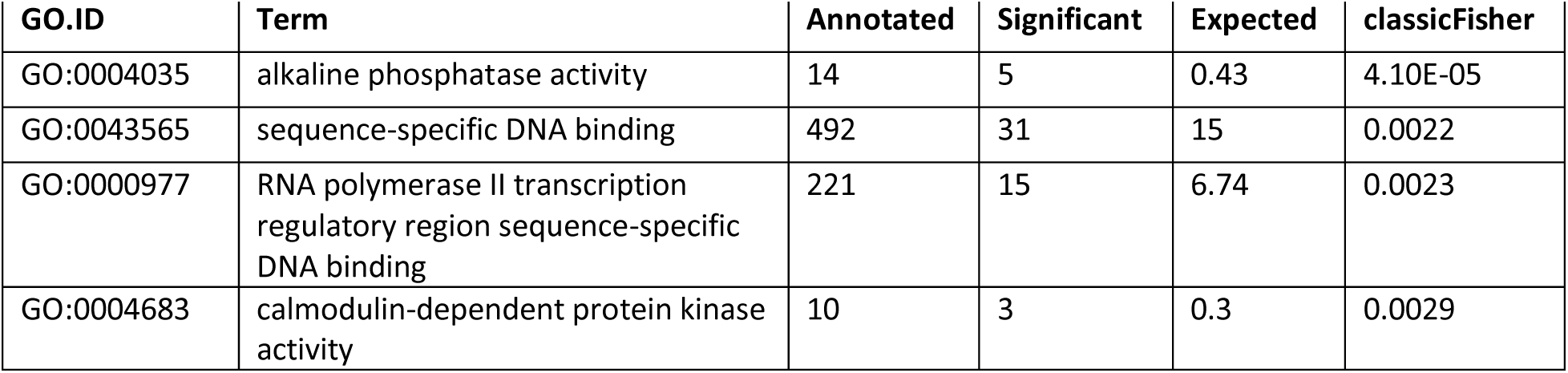
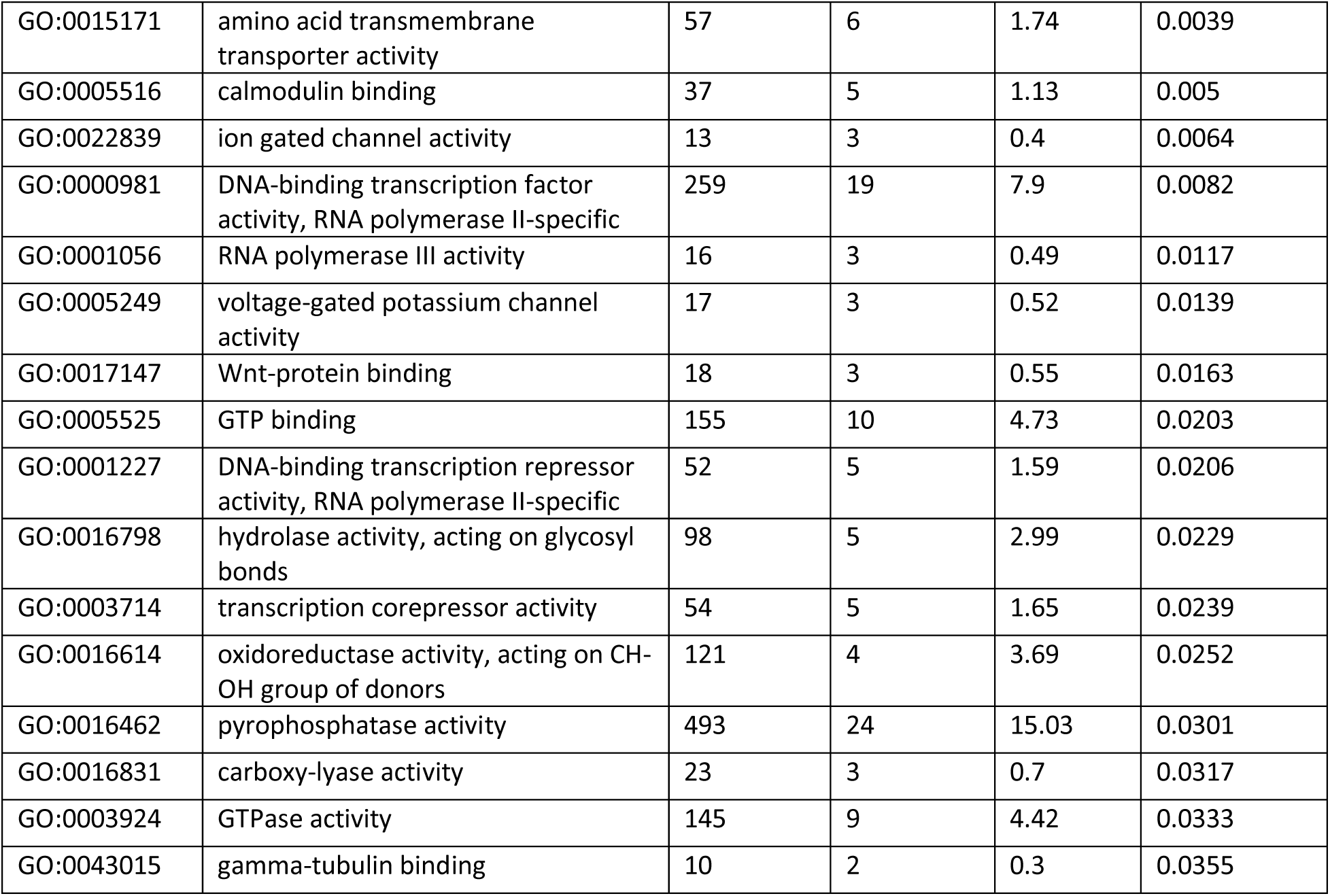
Small - Top 20 molecular function GO terms.

## Notes

### Competing Interest Statement

The authors have declared no competing interest.

### Summary of Updates

This is a a revised version of the manuscript after receiving feedback from editors and reviewers. In particular additional genomic analyses and comparisons have been added. Additionally an experiment examining adult sex ratios has been added.

