## SupplementalFiguresTables for "Sexually discordant selection is associated with trait specific morphological changes and a complex genomic response"

Table S1: Food recipe for artificial selection lineages. All ingredients except cornmeal, yeast, and preservatives are brought to a boil for 10 minutes, yeast/cornmeal and a small amount of reserved water are added and boiled for another 10 minutes, food is cooled to 60°C then preservatives are added before pouring ~10mL in to vials.

| | |
|---|---|
| Water | 1.2L |
| Agar | 11.1g |
| Cornmeal | 75.0g |
| Yeast | 31.0g |
| Molasses | 75mL |
| methyl-p-hydrobenzoic acid | 1.8g |
| 95% EtOH | 17.5mL |
| Propionic Acid | 7mL |

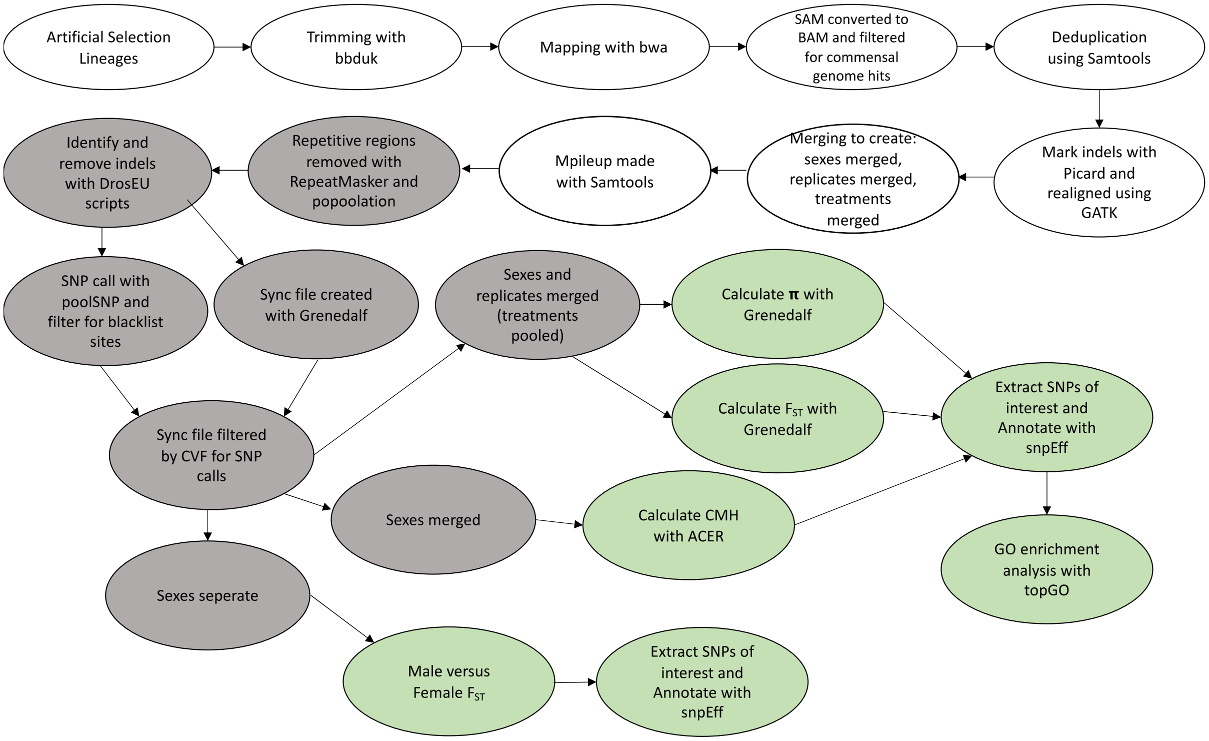


Figure S1: Flow chart showing steps and tools involved in the bioinformatic pipeline. Version numbers, parameter settings and more detail available in text. White: bam files, grey: intermediate files, green: output results files.

Table S2: model ANOVA for thorax

| | $\chi^2$ | **Df** | **Pr(>$\chi^2$)** |
|---|---|---|---|
| sex | 140.67 | 1 | 1.90E-32 |
| selection | 242.72 | 3 | 2.45E-52 |
| sampling | 18.9 | 1 | 1.37E-05 |
| sex:selection | 38.24 | 3 | 2.52E-08 |
| sex:sampling | 5.29 | 1 | 2.14E-02 |
| selection:sampling | 20.28 | 3 | 1.48E-04 |

Table S3: model ANOVA for femur

| | $\chi^2$ | **Df** | **Pr(>$\chi^2$)** |
|---|---|---|---|
| sex | 2.26 | 1 | 1.33E-01 |
| selection | 109.62 | 3 | 1.32E-23 |
| sampling | 8.32 | 1 | 3.93E-03 |
| sex:selection | 21.6 | 3 | 7.89E-05 |
| sex:sampling | 6.13 | 1 | 1.33E-02 |
| selection:sampling | 10.59 | 3 | 1.41E-02 |

Table S4: model ANOVA for tibia

| | $\chi^2$ | **Df** | **Pr(>$\chi^2$)** |
|---|---|---|---|
| sex | 1.03 | 1 | 3.11E-01 |
| selection | 295.82 | 3 | 7.98E-64 |
| sampling | 28.33 | 1 | 1.02E-07 |
| sex:selection | 31.91 | 3 | 5.45E-07 |
| sex:sampling | 9.11 | 1 | 2.54E-03 |
| selection:sampling | 13.71 | 3 | 3.33E-03 |

Table S5: model ANOVA for tarsus

| | $\chi^2$ | **Df** | **Pr(>$\chi^2$)** |
|---|---|---|---|
| sex | 36.71 | 1 | 1.37E-09 |
| selection | 133.67 | 3 | 8.74E-29 |
| sampling | 2.86 | 1 | 9.10E-02 |
| sex:selection | 13.44 | 3 | 3.77E-03 |
| sex:sampling | 4.26 | 1 | 3.90E-02 |
| selection:sampling | 17.5 | 3 | 5.58E-04 |

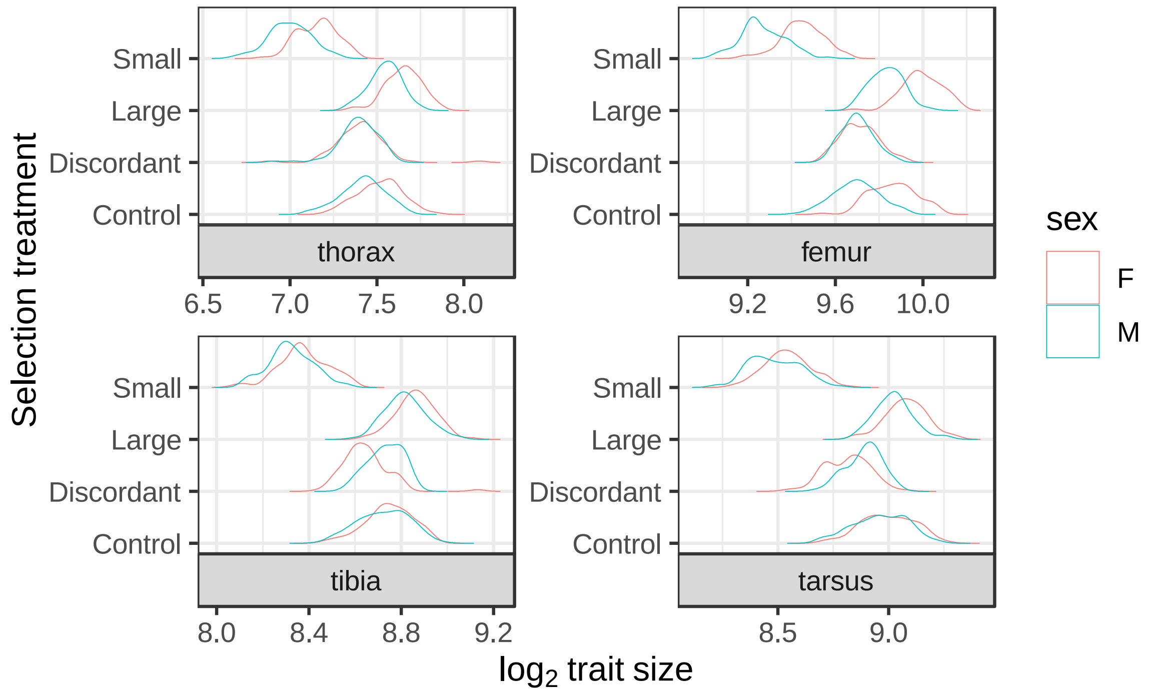


Figure S2: Density plots of raw measures ($log_2$ transformed) for traits and sexes

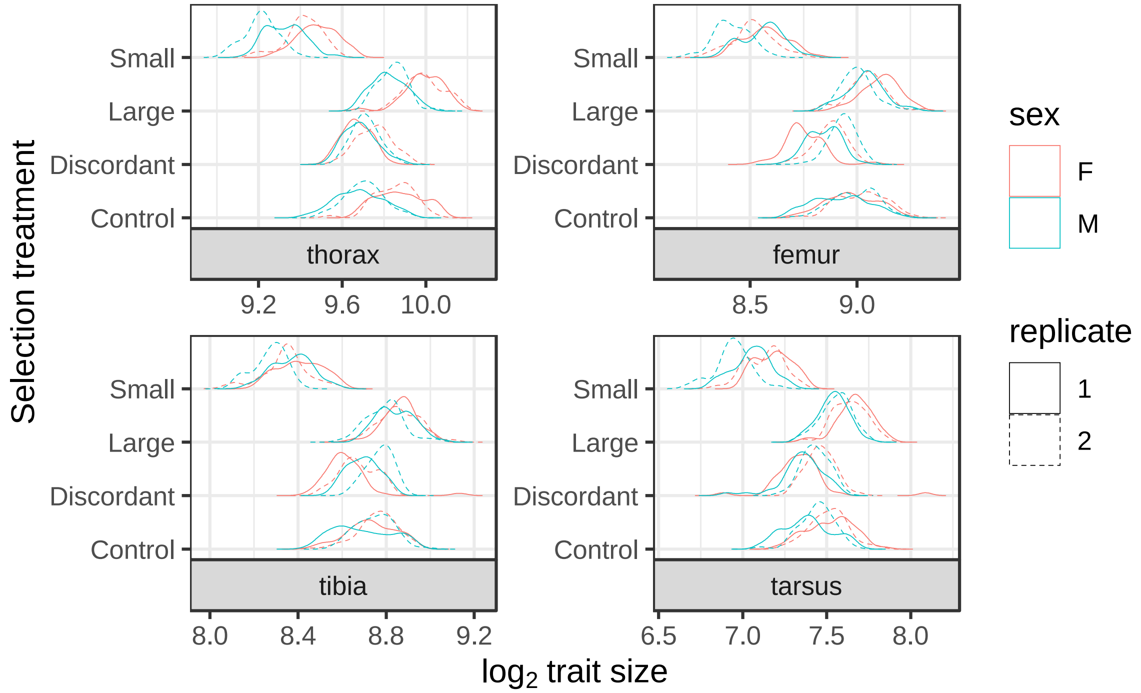


Figure S3: Density plots of raw measures ($log_2$ transformed) for traits and sexes with replicates separated to show replicate heterogeneity

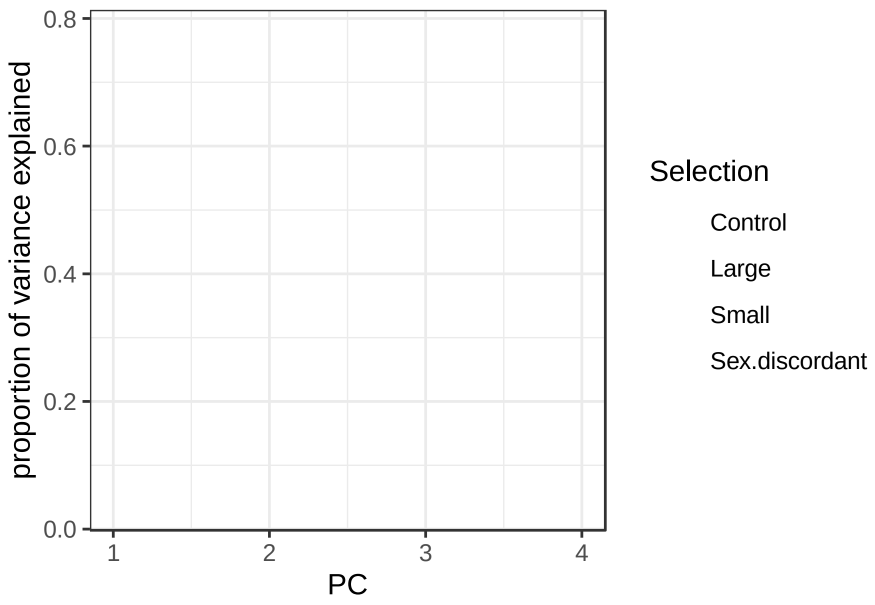


Figure S4: scree plot of proportion of variation explained by each associated eigenvector.

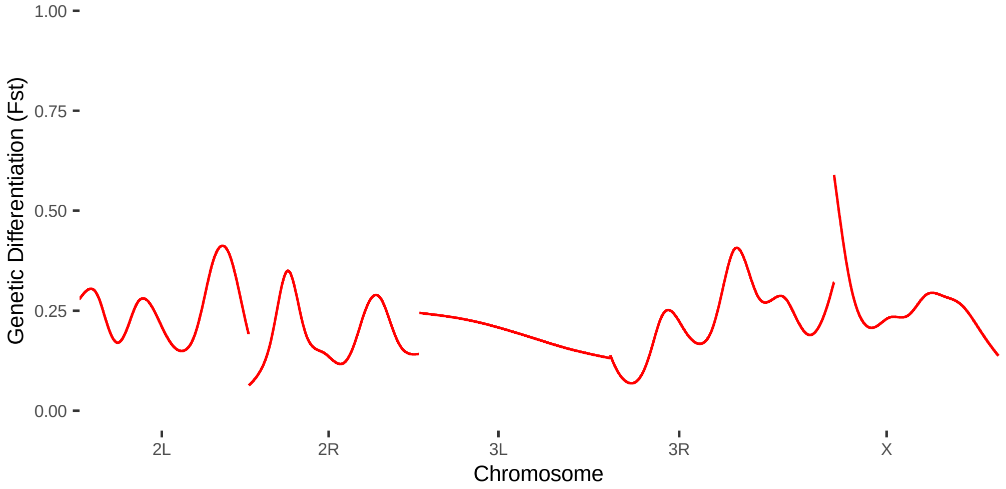


Figure S5: discordant vs. large 10000bp windows

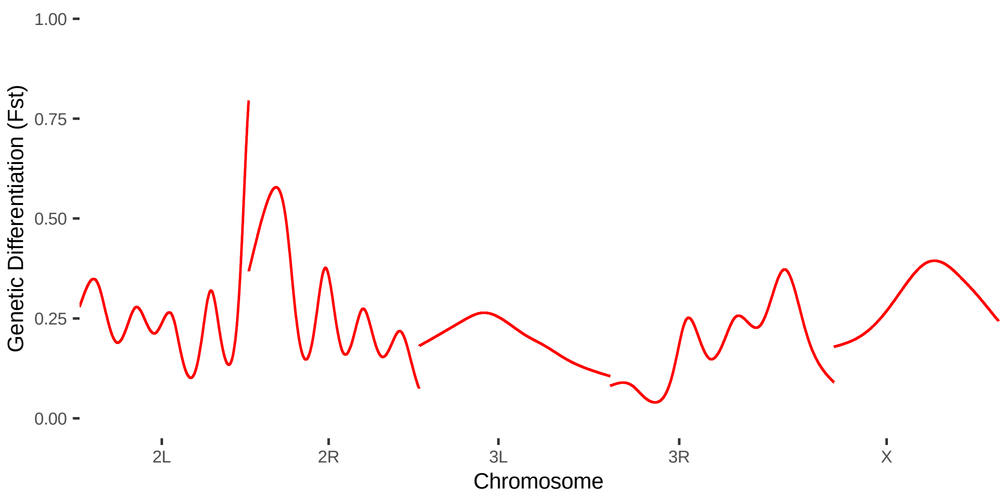


Figure S6: discordant vs. small 10000bp windows

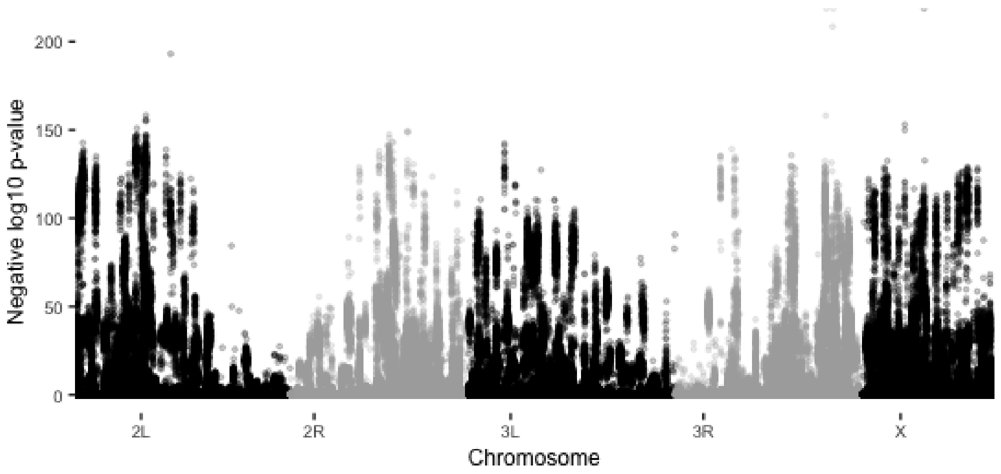


Figure S7: CMH p-values for Large vs. Small treatments

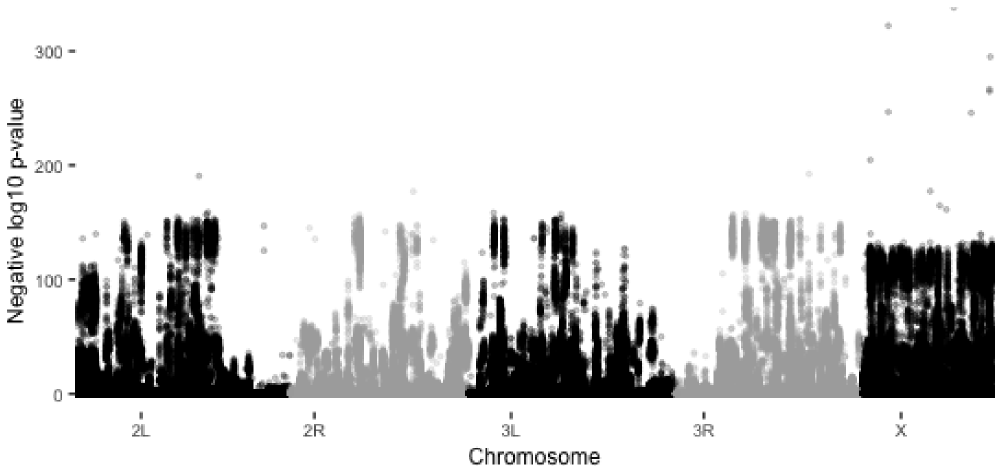


Figure S8: CMH p-values for discordant vs. control treatments

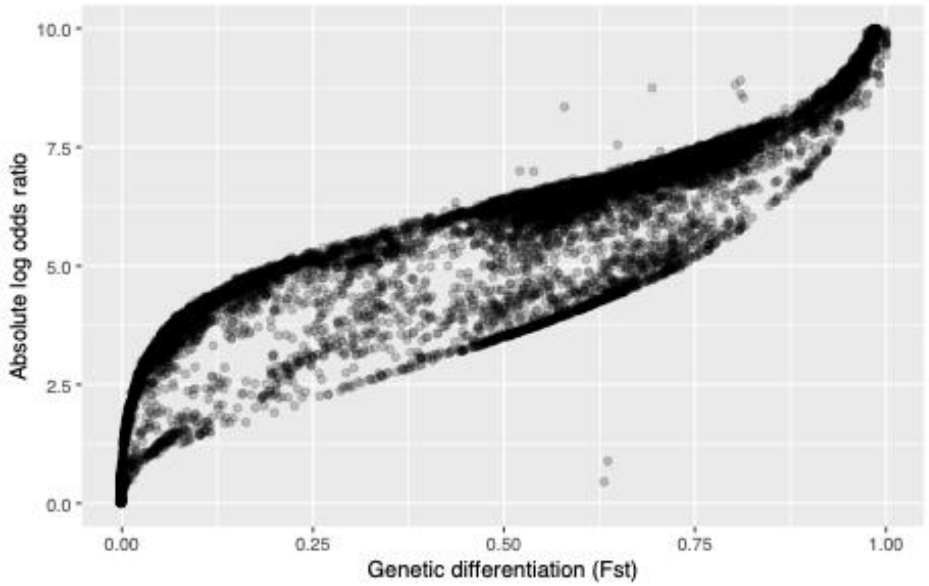


Figure S9: Correlation between $F_{ST}$ and the log odds from our model estimates for the Discordant vs. control comparisons

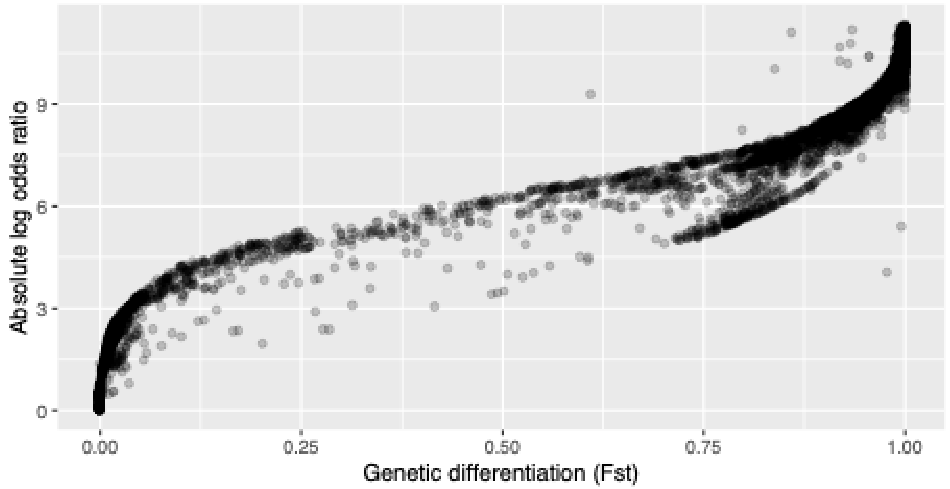


Figure S10: Correlation between Large vs. Small $F_{ST}$ and the log odds from our model estimates for the SNPs found to be associated with the Large treatment

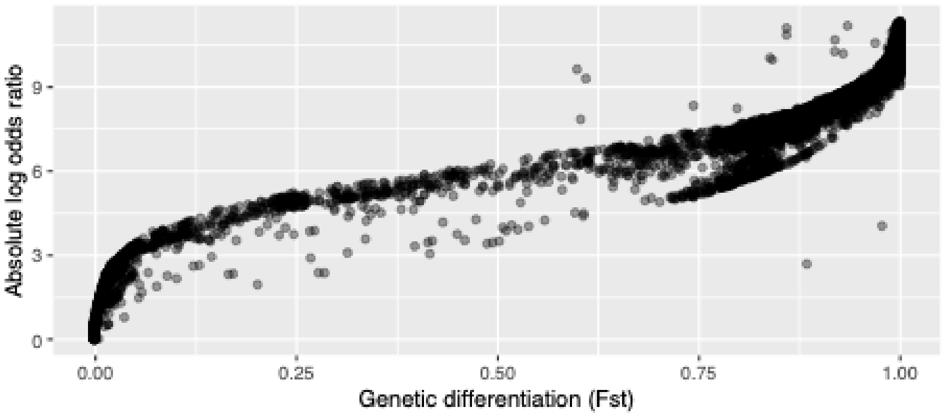


Figure S11: Correlation between Large vs. Small $F_{ST}$ and the log odds from our model estimates for the SNPs found to be associated with the Small treatment

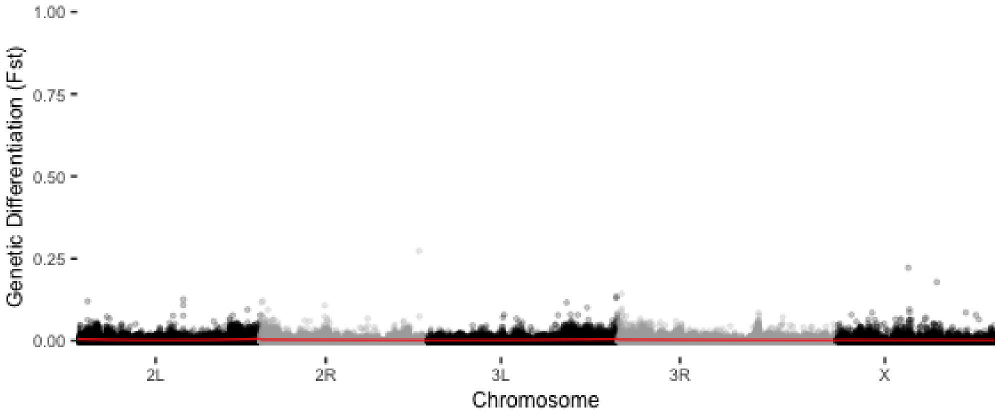


Figure S12: control replicate 2 males vs. females 1000bp

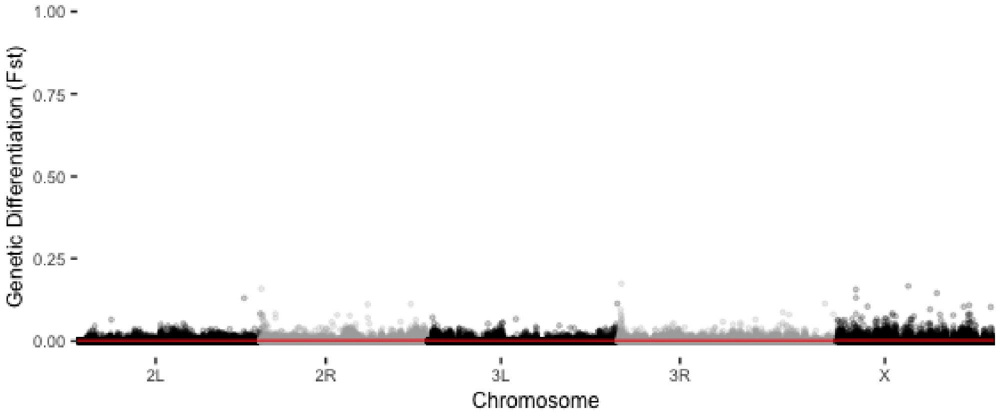


Figure S13: large replicate 1 males vs. females 1000bp

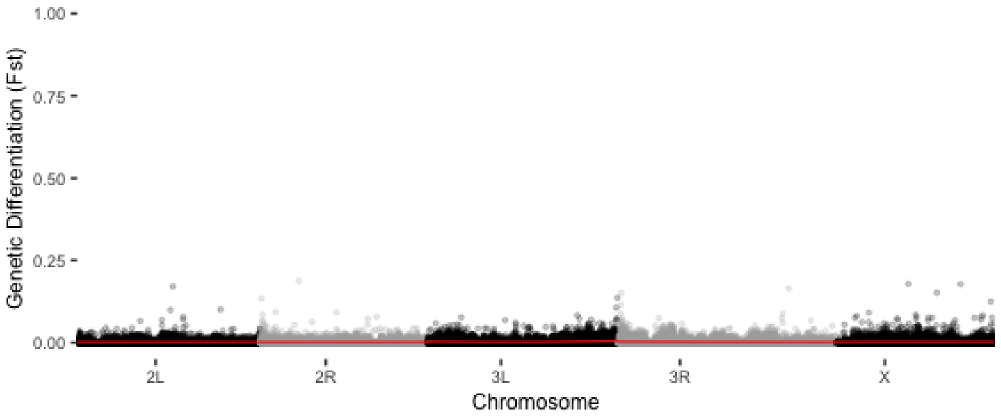

Figure S14: large replicate 2 males vs. females 1000bp

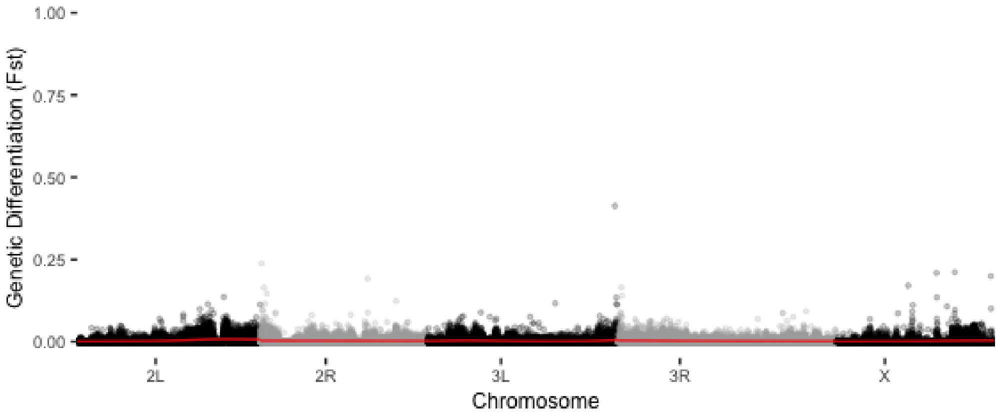

Figure S15: small replicate 1 males vs. females 1000bp

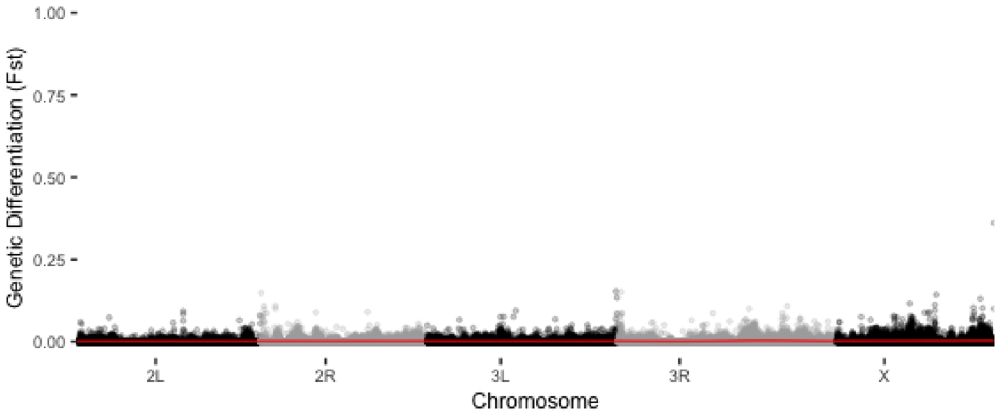

Figure S16: small replicate 2 males vs. females 1000bp

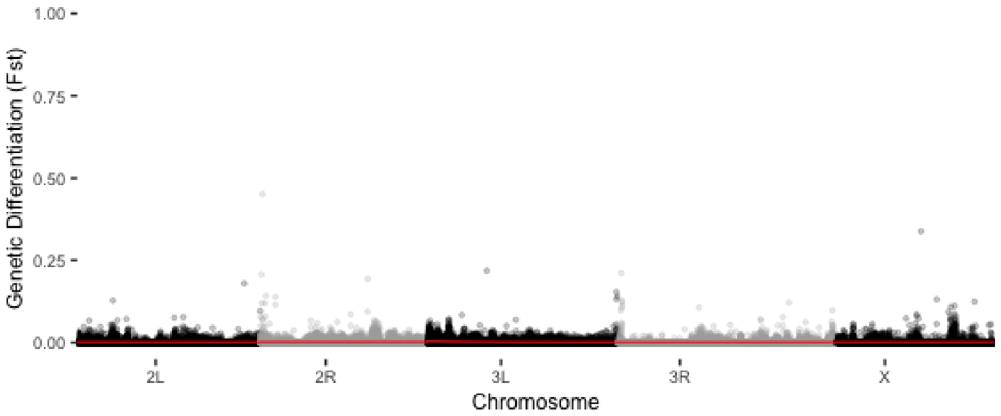


Figure S17: discordant replicate 2 males vs. females 1000bp

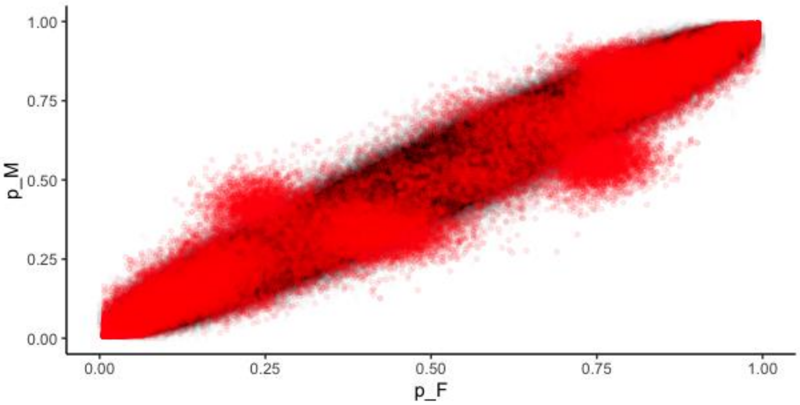


Figure S18: E1 male versus female minor allele frequency (computed as ALT/(REF+ALT)) in red plotted on top of 1000000 simulated variants with similar coverage and allele frequency distributions of our data.

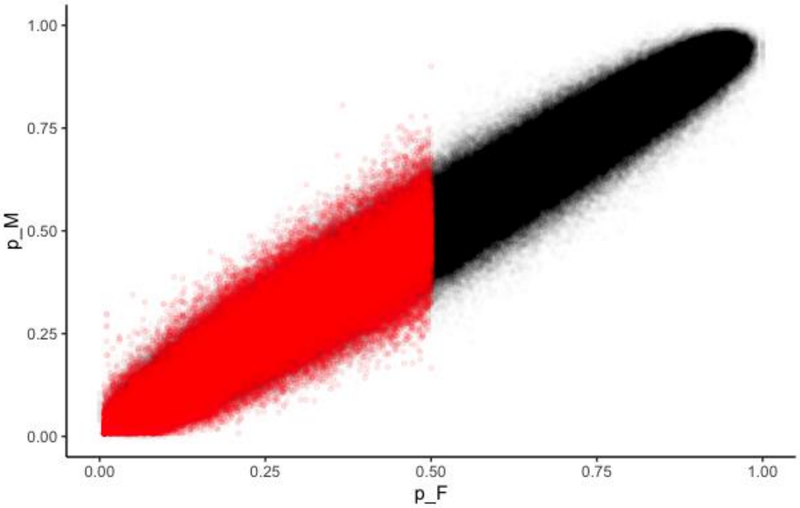


Figure S19: C1 male versus female minor allele frequency (computed as ALT/(REF+ALT)) in red plotted on top of 1000000 samples binomially sampled from a simulated pool with a similar coverage and distribution of our data. ***Please note***, the reference allele frequency for all treatments are set to the major allele of C1 females (this sample) so no alternate alleles are expected to reach >0.5 frequency.

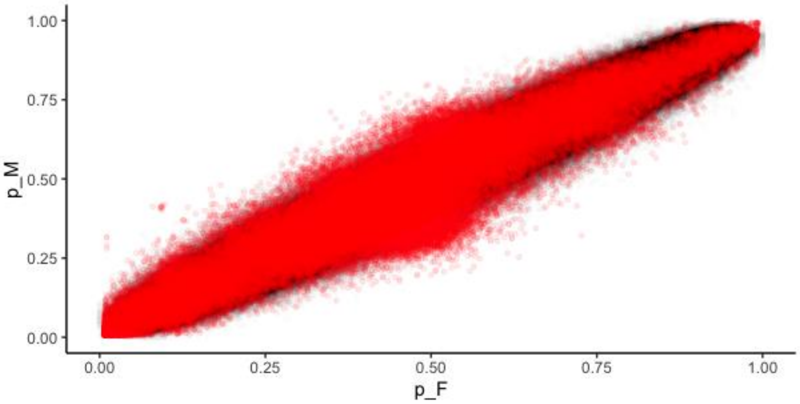


Figure S20: C2 male versus female minor allele frequency (computed as ALT/(REF+ALT)) in red plotted on top of 1000000 samples binomially sampled from a simulated pool with a similar coverage and distribution of our data.

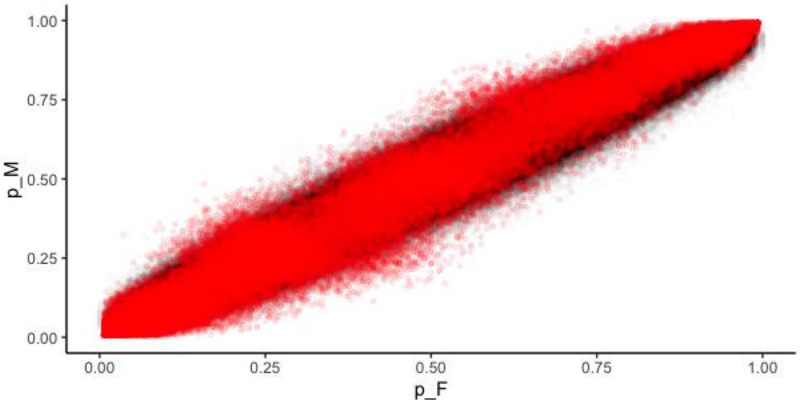


Figure S21: E2 male versus female minor allele frequency (computed as ALT/(REF+ALT)) in red plotted on top of 1000000 samples binomially sampled from a simulated pool with a similar coverage and distribution of our data.

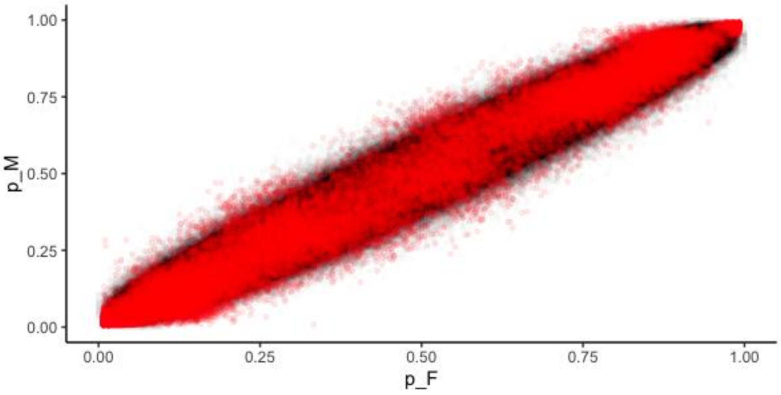

Figure S22: L1 male versus female minor allele frequency (computed as ALT/(REF+ALT)) in red plotted on top of 1000000 samples binomially sampled from a simulated pool with a similar coverage and distribution of our data.

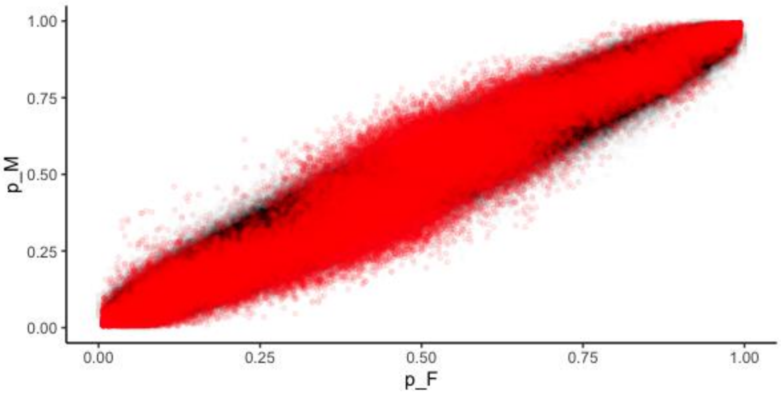


Figure S23: L2 male versus female minor allele frequency (computed as ALT/(REF+ALT)) in red plotted on top of 1000000 samples binomially sampled from a simulated pool with a similar coverage and distribution of our data.

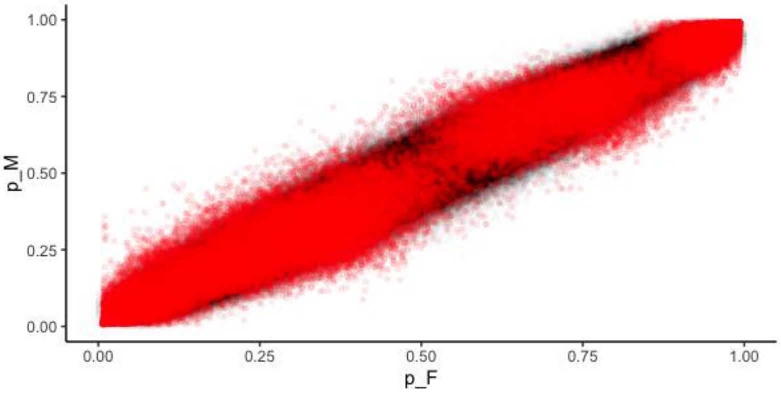


Figure S24: S1 male versus female minor allele frequency (computed as ALT/(REF+ALT)) in red plotted on top of 1000000 samples binomially sampled from a simulated pool with a similar coverage and distribution of our data.

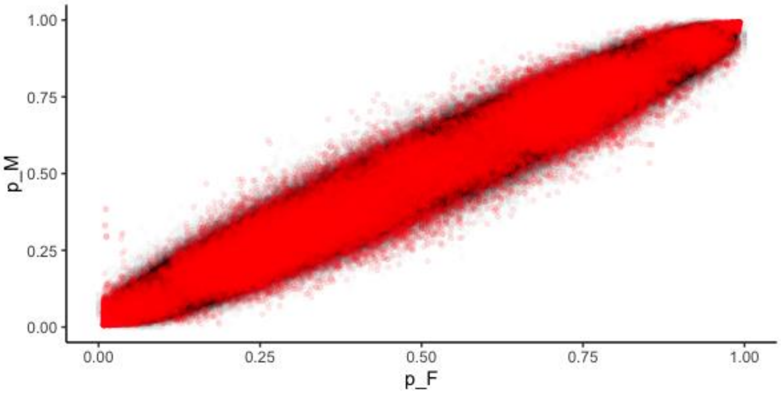

Figure S25: S2 male versus female minor allele frequency (computed as ALT/(REF+ALT)) in red plotted on top of 1000000 samples binomially sampled from a simulated pool with a similar coverage and distribution of our data.

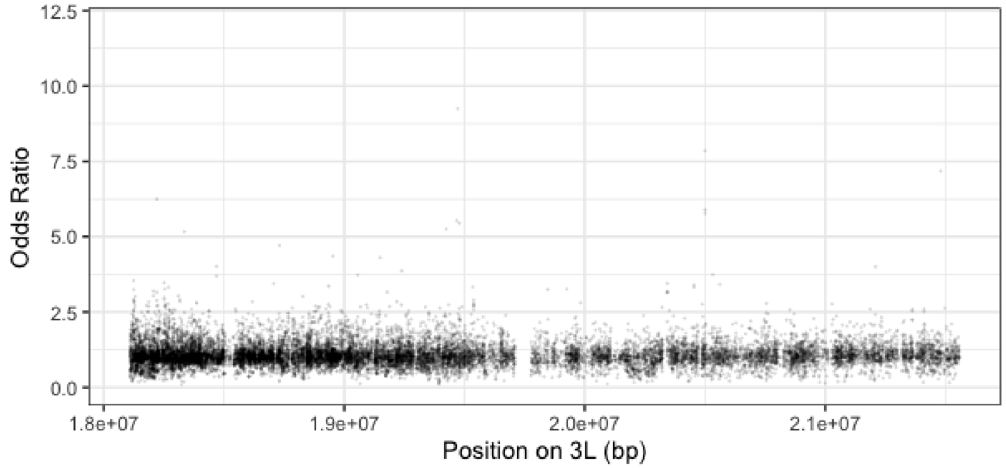


Figure S26: C1 modelled odds ratios by position on chromosome 3L within the elevated region on E1. Positions with a p-value < 0.0001 highlighted red with upper and lower 95% confidence intervals shown.

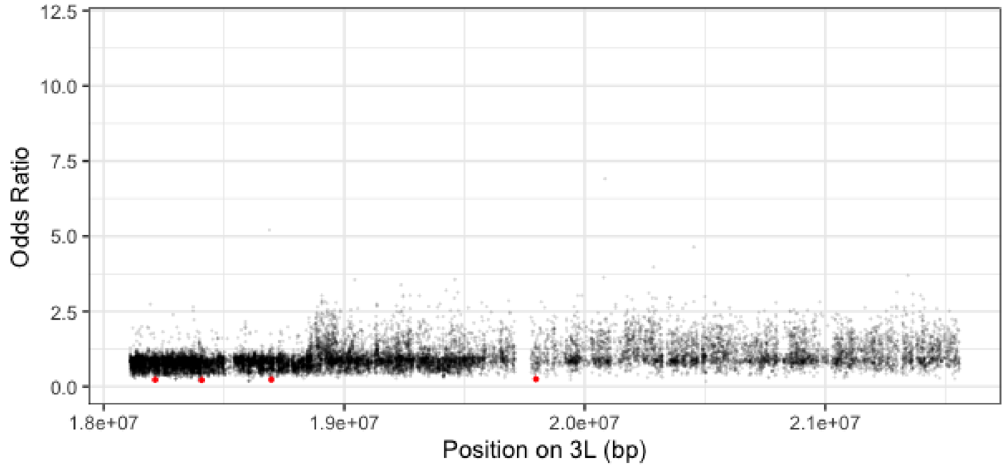


Figure S27: C2 modelled odds ratios by position on chromosome 3L within the elevated region on E1. Positions with a p-value < 0.0001 highlighted red with upper and lower 95% confidence intervals shown.

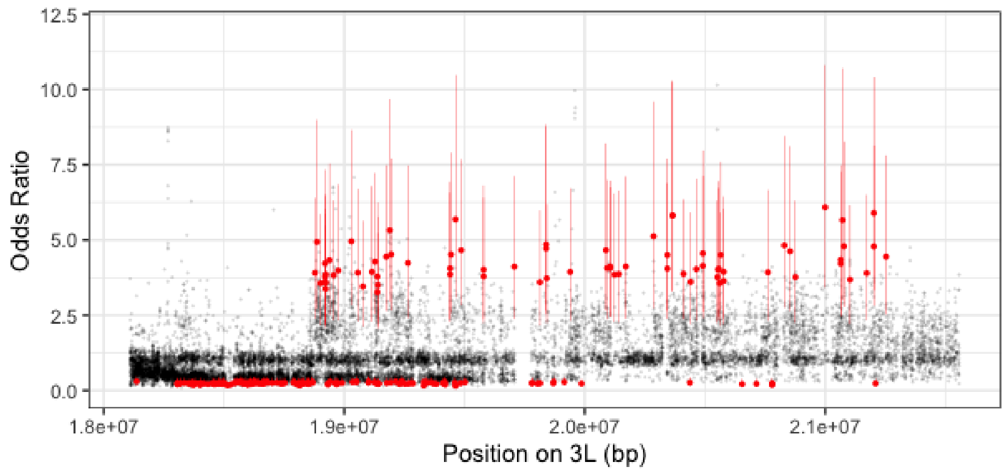


Figure S28: E1 modelled odds ratios by position on chromosome 3L within the elevated region on E1. Positions with a p-value < 0.0001 highlighted red with upper and lower 95% confidence intervals shown.

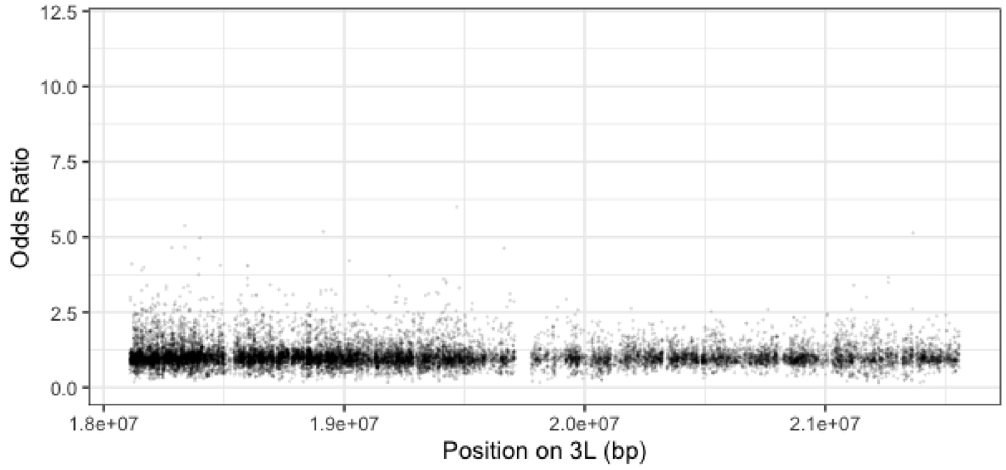


Figure S29: E2 modelled odds ratios by position on chromosome 3L within the elevated region on E1. Positions with a p-value < 0.0001 highlighted red with upper and lower 95% confidence intervals shown.

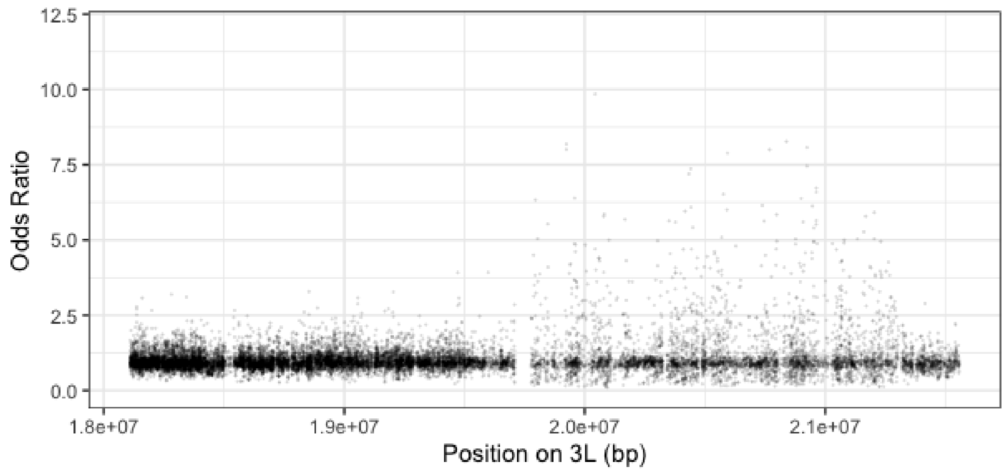


Figure S30: L1 modelled odds ratios by position on chromosome 3L within the elevated region on E1. Positions with a p-value < 0.0001 highlighted red with upper and lower 95% confidence intervals shown.

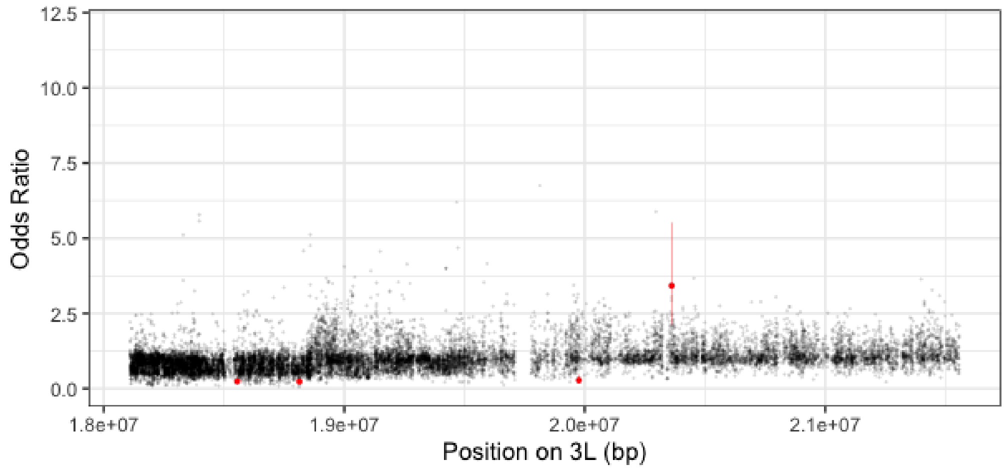


Figure S31: L2 modelled odds ratios by position on chromosome 3L within the elevated region on E1. Positions with a p-value < 0.0001 highlighted red with upper and lower 95% confidence intervals shown.

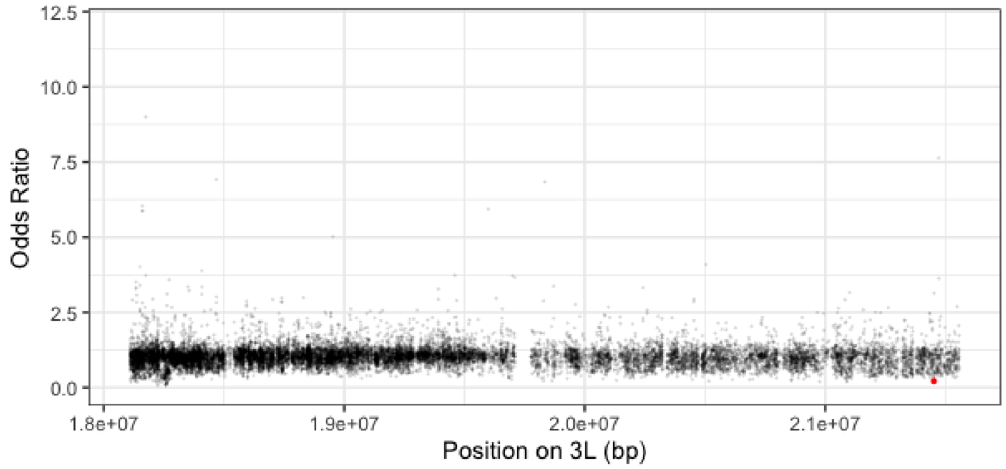

Figure S32: S1 modelled odds ratios by position on chromosome 3L within the elevated region on E1. Positions with a p-value < 0.0001 highlighted red with upper and lower 95% confidence intervals shown.

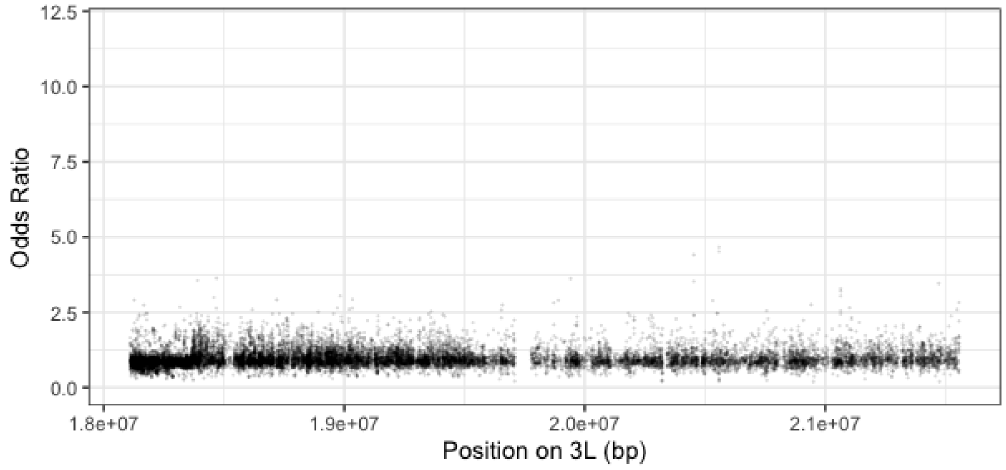


Figure S33: S2 modelled odds ratios by position on chromosome 3L within the elevated region on E1. Positions with a p-value < 0.0001 highlighted red with upper and lower 95% confidence intervals shown.

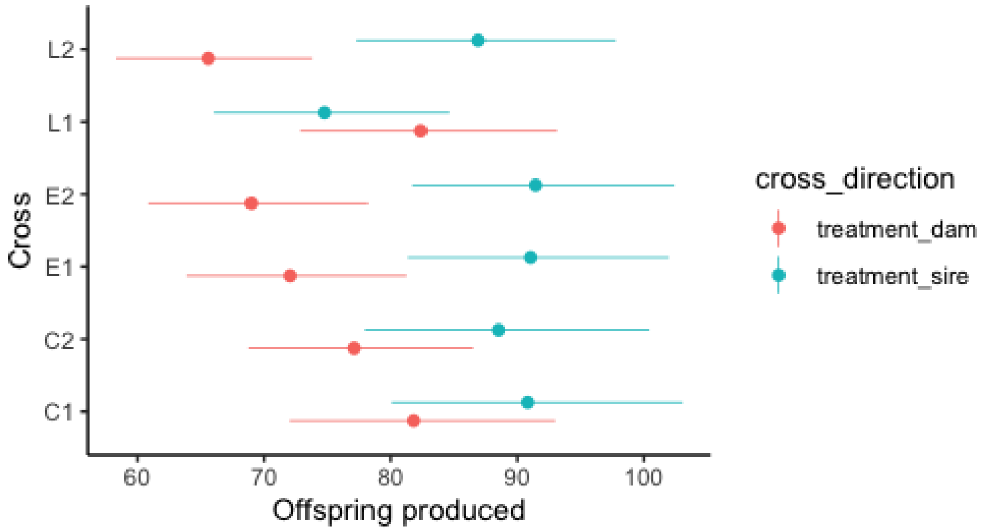


Figure S34: Fecundity measure for F1 of our sex ratio cross based on F0 treatment

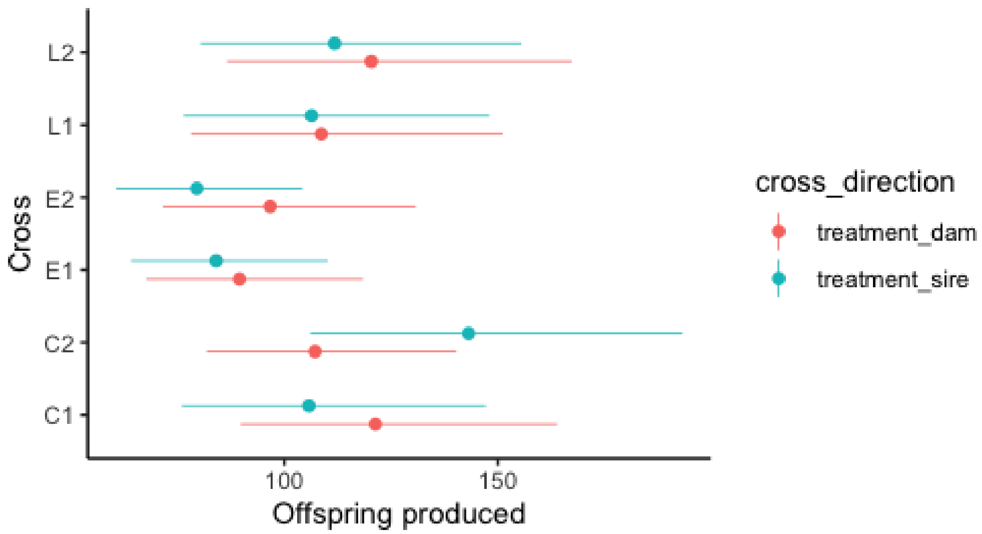


Figure S35: Fecundity measure for F2 of our sex ratio cross based on F0 treatment

Table S6: Discordant -Top 20 biological process GO terms

| GO.ID | Term | Annotated | Significant | Expected | classicFisher |
|---|---|---|---|---|---|
| GO:0050916 | sensory perception of sweet taste | 14 | 5 | 0.48 | 7.20E-05 |
| GO:0048749 | compound eye development | 377 | 24 | 12.95 | 0.00017 |
| GO:0006814 | sodium ion transport | 66 | 11 | 2.27 | 0.0002 |
| GO:0007424 | open tracheal system development | 241 | 19 | 8.28 | 0.00029 |
| GO:0050912 | detection of chemical stimulus involved in sensory perception of taste | 23 | 5 | 0.79 | 0.00094 |
| GO:0007409 | axonogenesis | 337 | 24 | 11.58 | 0.00154 |
| GO:0045572 | positive regulation of imaginal disc growth | 16 | 4 | 0.55 | 0.0018 |
| GO:0007615 | anesthesia-resistant memory | 17 | 4 | 0.58 | 0.00229 |
| GO:0051124 | synaptic growth at neuromuscular junction | 139 | 10 | 4.77 | 0.00277 |
| GO:0007614 | short-term memory | 29 | 5 | 1 | 0.0028 |
| GO:0036099 | female germ-line stem cell population ma... | 29 | 5 | 1 | 0.0028 |
| GO:0099024 | plasma membrane invagination | 26 | 4 | 0.89 | 0.00289 |
| GO:0007455 | eye-antennal disc morphogenesis | 42 | 6 | 1.44 | 0.00291 |
| GO:0007584 | response to nutrient | 19 | 4 | 0.65 | 0.00353 |
| GO:0070839 | metal ion export | 10 | 3 | 0.34 | 0.00403 |
| GO:0045187 | regulation of circadian sleep/wake cycle, sleep | 36 | 6 | 1.24 | 0.00424 |
| GO:0019367 | fatty acid elongation, saturated fatty acid | 20 | 4 | 0.69 | 0.00429 |
| GO:0034625 | fatty acid elongation, monounsaturated fatty acid | 20 | 4 | 0.69 | 0.00429 |

| | | | | | |
|---|---|---|---|---|---|
| GO:0034626 | fatty acid elongation, polyunsaturated fatty acid | 20 | 4 | 0.69 | 0.00429 |
| GO:0035158 | regulation of tube diameter, open tracheal system | 11 | 3 | 0.38 | 0.0054 |

Table S7: Large - Top 20 biological process GO terms

| GO.ID | Term | Annotated | Significant | Expected | classicFisher |
|---|---|---|---|---|---|
| GO:0048512 | circadian behavior | 110 | 8 | 3.18 | 0.00082 |
| GO:0035209 | pupal development | 17 | 4 | 0.49 | 0.00121 |
| GO:0051124 | synaptic growth at neuromuscular junction | 139 | 12 | 4.02 | 0.00125 |
| GO:0016319 | mushroom body development | 74 | 8 | 2.14 | 0.00129 |
| GO:0007616 | long-term memory | 75 | 8 | 2.17 | 0.00141 |
| GO:0010628 | positive regulation of gene expression | 191 | 14 | 5.52 | 0.00227 |
| GO:0006821 | chloride transport | 29 | 4 | 0.84 | 0.00433 |
| GO:0070828 | heterochromatin organization | 59 | 5 | 1.71 | 0.00478 |
| GO:0060070 | canonical Wnt signaling pathway | 93 | 7 | 2.69 | 0.00545 |
| GO:0007297 | ovarian follicle cell migration | 138 | 8 | 3.99 | 0.00546 |
| GO:0017156 | calcium-ion regulated exocytosis | 13 | 3 | 0.38 | 0.00553 |
| GO:0007474 | imaginal disc-derived wing vein specification | 57 | 6 | 1.65 | 0.00587 |
| GO:0007619 | courtship behavior | 120 | 7 | 3.47 | 0.00683 |
| GO:0045944 | positive regulation of transcription by RNA polymerase II | 331 | 18 | 9.57 | 0.00773 |
| GO:0030177 | positive regulation of Wnt signaling pathway | 43 | 5 | 1.24 | 0.00779 |
| GO:0035249 | synaptic transmission, glutamatergic | 26 | 4 | 0.75 | 0.00838 |
| GO:0008586 | imaginal disc-derived wing vein morphogenesis | 44 | 5 | 1.27 | 0.00846 |
| GO:0016311 | dephosphorylation | 196 | 9 | 5.67 | 0.0085 |
| GO:0051260 | protein homooligomerization | 29 | 4 | 0.84 | 0.00922 |
| GO:0031122 | cytoplasmic microtubule organization | 29 | 4 | 0.84 | 0.00922 |

Table S8: Small Top 20 biological process GO terms

| GO.ID | Term | Annotated | Significant | Expected | classicFisher |
|---|---|---|---|---|---|
| GO:0048512 | circadian behavior | 110 | 8 | 3.37 | 0.00092 |
| GO:0007615 | anesthesia-resistant memory | 17 | 4 | 0.52 | 0.0015 |
| GO:0035209 | pupal development | 17 | 4 | 0.52 | 0.0015 |
| GO:0016319 | mushroom body development | 74 | 8 | 2.27 | 0.00185 |
| GO:0016226 | iron-sulfur cluster assembly | 20 | 4 | 0.61 | 0.00284 |
| GO:0048675 | axon extension | 54 | 7 | 1.65 | 0.004 |
| GO:0006821 | chloride transport | 29 | 4 | 0.89 | 0.00508 |

| | | | | | |
|---|---|---|---|---|---|
| GO:1902808 | positive regulation of cell cycle G1/S phase transition | 12 | 3 | 0.37 | 0.0051 |
| GO:0070828 | heterochromatin organization | 59 | 6 | 1.81 | 0.00532 |
| GO:0060070 | canonical Wnt signaling pathway | 93 | 7 | 2.85 | 0.0064 |
| GO:0045455 | ecdysteroid metabolic process | 40 | 5 | 1.22 | 0.00642 |
| GO:0017156 | calcium-ion regulated exocytosis | 13 | 3 | 0.4 | 0.00648 |
| GO:0008045 | motor neuron axon guidance | 75 | 7 | 2.3 | 0.00793 |
| GO:0007616 | long-term memory | 75 | 7 | 2.3 | 0.00793 |
| GO:0007619 | courtship behavior | 120 | 7 | 3.67 | 0.00801 |
| GO:0030177 | positive regulation of Wnt signaling pathway | 43 | 5 | 1.32 | 0.00871 |
| GO:0030154 | cell differentiation | 1703 | 81 | 52.14 | 0.00937 |
| GO:0016311 | dephosphorylation | 196 | 9 | 6 | 0.01075 |
| GO:0051124 | synaptic growth at neuromuscular junction | 139 | 11 | 4.26 | 0.01082 |
| GO:0008016 | regulation of heart contraction | 29 | 4 | 0.89 | 0.01121 |

Table S9: discordant - Top 20 molecular function GO terms

| GO.ID | Term | Annotated | Significant | Expected | classicFisher |
|---|---|---|---|---|---|
| GO:0015491 | cation:cation antiporter activity | 14 | 4 | 0.48 | 0.001 |
| GO:0005319 | lipid transporter activity | 46 | 6 | 1.58 | 0.0032 |
| GO:0033041 | sweet taste receptor activity | 30 | 5 | 1.03 | 0.0033 |
| GO:0008252 | nucleotidase activity | 10 | 3 | 0.34 | 0.004 |
| GO:0022848 | acetylcholine-gated monoatomic cation-selective channel activity | 10 | 3 | 0.34 | 0.004 |
| GO:0009922 | fatty acid elongase activity | 20 | 4 | 0.69 | 0.0043 |
| GO:0015141 | succinate transmembrane transporter activity | 11 | 3 | 0.38 | 0.0054 |
| GO:0008227 | G protein-coupled amine receptor activity | 22 | 4 | 0.75 | 0.0061 |
| GO:0004596 | peptide alpha-N-acetyltransferase activiy | 13 | 3 | 0.45 | 0.0089 |
| GO:0046872 | metal ion binding | 1047 | 44 | 35.93 | 0.0107 |
| GO:0070566 | adenylyltransferase activity | 14 | 3 | 0.48 | 0.011 |
| GO:0016791 | phosphatase activity | 180 | 15 | 6.18 | 0.0119 |
| GO:0005201 | extracellular matrix structural constituent | 30 | 4 | 1.03 | 0.0185 |
| GO:0008201 | heparin binding | 17 | 3 | 0.58 | 0.0191 |
| GO:0008270 | zinc ion binding | 518 | 27 | 17.78 | 0.0201 |
| GO:0000976 | transcription cis-regulatory region binding | 296 | 21 | 10.16 | 0.0249 |
| GO:0003777 | microtubule motor activity | 65 | 7 | 2.23 | 0.0292 |
| GO:0042826 | histone deacetylase binding | 20 | 3 | 0.69 | 0.0296 |

| | | | | | |
|---|---|---|---|---|---|
| GO:0000977 | RNA polymerase II transcription regulatory region sequence-specific DNA binding | 221 | 15 | 7.58 | 0.0364 |
| GO:0000166 | nucleotide binding | 959 | 41 | 32.91 | 0.0365 |

Table S10: Large - Top 20 molecular function GO terms

| GO.ID | Term | Annotated | Significant | Expected | classicFisher |
|---|---|---|---|---|---|
| GO:0004035 | alkaline phosphatase activity | 14 | 5 | 0.4 | 2.90E-05 |
| GO:0000977 | RNA polymerase II transcription regulatory region sequence-specific DNA binding | 221 | 17 | 6.29 | 0.00034 |
| GO:0022839 | ion gated channel activity | 13 | 3 | 0.37 | 0.00529 |
| GO:0043565 | sequence-specific DNA binding | 492 | 30 | 14.01 | 0.0092 |
| GO:0001056 | RNA polymerase III activity | 16 | 3 | 0.46 | 0.00973 |
| GO:0005249 | voltage-gated potassium channel activity | 17 | 3 | 0.48 | 0.01157 |
| GO:0017147 | Wnt-protein binding | 18 | 3 | 0.51 | 0.0136 |
| GO:0000981 | DNA-binding transcription factor activity, RNA polymerase II-specific | 259 | 18 | 7.38 | 0.01481 |
| GO:0001227 | DNA-binding transcription repressor activity, RNA polymerase II-specific | 52 | 5 | 1.48 | 0.01582 |
| GO:0003714 | transcription corepressor activity | 54 | 5 | 1.54 | 0.01839 |
| GO:0016798 | hydrolase activity, acting on glycosyl bonds | 98 | 4 | 2.79 | 0.02028 |
| GO:0005516 | calmodulin binding | 37 | 4 | 1.05 | 0.02037 |
| GO:0015171 | amino acid transmembrane transporter activity | 57 | 5 | 1.62 | 0.02274 |
| GO:0016462 | pyrophosphatase activity | 493 | 21 | 14.04 | 0.02558 |
| GO:0016831 | carboxy-lyase activity | 23 | 3 | 0.66 | 0.02659 |
| GO:0004683 | calmodulin-dependent protein kinase activity | 10 | 2 | 0.28 | 0.03129 |
| GO:0043015 | gamma-tubulin binding | 10 | 2 | 0.28 | 0.03129 |
| GO:0004112 | cyclic-nucleotide phosphodiesterase activity | 10 | 2 | 0.28 | 0.03129 |
| GO:0005254 | chloride channel activity | 26 | 3 | 0.74 | 0.03668 |
| GO:0004860 | protein kinase inhibitor activity | 11 | 2 | 0.31 | 0.03753 |

Table S11: Small - Top 20 molecular function GO terms

| GO.ID | Term | Annotated | Significant | Expected | classicFisher |
|---|---|---|---|---|---|
| GO:0004035 | alkaline phosphatase activity | 14 | 5 | 0.43 | 4.10E-05 |
| GO:0043565 | sequence-specific DNA binding | 492 | 31 | 15 | 0.0022 |
| GO:0000977 | RNA polymerase II transcription regulatory region sequence-specific DNA binding | 221 | 15 | 6.74 | 0.0023 |
| GO:0004683 | calmodulin-dependent protein kinase activity | 10 | 3 | 0.3 | 0.0029 |

| | | | | | |
|---|---|---|---|---|---|
| GO:0015171 | amino acid transmembrane transporter activity | 57 | 6 | 1.74 | 0.0039 |
| GO:0005516 | calmodulin binding | 37 | 5 | 1.13 | 0.005 |
| GO:0022839 | ion gated channel activity | 13 | 3 | 0.4 | 0.0064 |
| GO:0000981 | DNA-binding transcription factor activity, RNA polymerase II-specific | 259 | 19 | 7.9 | 0.0082 |
| GO:0001056 | RNA polymerase III activity | 16 | 3 | 0.49 | 0.0117 |
| GO:0005249 | voltage-gated potassium channel activity | 17 | 3 | 0.52 | 0.0139 |
| GO:0017147 | Wnt-protein binding | 18 | 3 | 0.55 | 0.0163 |
| GO:0005525 | GTP binding | 155 | 10 | 4.73 | 0.0203 |
| GO:0001227 | DNA-binding transcription repressor activity, RNA polymerase II-specific | 52 | 5 | 1.59 | 0.0206 |
| GO:0016798 | hydrolase activity, acting on glycosyl bonds | 98 | 5 | 2.99 | 0.0229 |
| GO:0003714 | transcription corepressor activity | 54 | 5 | 1.65 | 0.0239 |
| GO:0016614 | oxidoreductase activity, acting on CH-OH group of donors | 121 | 4 | 3.69 | 0.0252 |
| GO:0016462 | pyrophosphatase activity | 493 | 24 | 15.03 | 0.0301 |
| GO:0016831 | carboxy-lyase activity | 23 | 3 | 0.7 | 0.0317 |
| GO:0003924 | GTPase activity | 145 | 9 | 4.42 | 0.0333 |
| GO:0043015 | gamma-tubulin binding | 10 | 2 | 0.3 | 0.0355 |
